# Detection of interphylum transfers of the magnetosome gene cluster in magnetotactic bacteria

**DOI:** 10.1101/2022.03.28.486042

**Authors:** Maria Uzun, Veronika Koziaeva, Marina Dziuba, Pedro Leão, Maria Krutkina, Denis Grouzdev

**Author notes:** authors with equal contribution (Maria Uzun, Veronika Koziaeva). Department of Marine Science, The University of Texas at Austin, Port Aransas, USA.

## Abstract

Magnetosome synthesis in magnetotactic bacteria (MTB) is regarded as a very ancient evolutionary process that dates back to deep-branching phyla. MTB belonging to one of such phyla, *Nitrospirota*, contain the classical genes for the magnetosome synthesis (e.g., *mam, mms*) and *man* genes, which were considered to be specific for this group. However, the recent discovery of *man* genes in MTB from the *Thermodesulfobacteriota* phylum has raised several questions about the inheritance of these genes in MTB. In this work, three new *man* genes containing MTB genomes affiliated with *Nitrospirota* and *Thermodesulfobacteriota*, were obtained. By applying reconciliation with these and the previously published MTB genomes, we demonstrate that the last common ancestor of all *Nitrospirota* was most likely not magnetotactic as assumed previously. Instead, our findings suggest that the genes for magnetosome synthesis were transmitted to the phylum *Nitrospirota* by horizontal gene transfer (HGT), which is the first case of the interphylum transfer of magnetosome genes detected to date. Furthermore, we provide evidence for the HGT of magnetosome genes from the *Magnetobacteriaceae* to the *Dissulfurispiraceae* family within *Nitrospirota*. Thus, our results imply a more significant role of HGT in the MTB evolution than deemed before and challenge the hypothesis of the ancient origin of magnetosome synthesis.

## Introduction

Bacteria that have an ability to form magnetosomes, magnetotactic bacteria (MTB), were found in phylogenetically distant taxa such as *Pseudomonadota* (former *Proteobacteria*, replaced by Oren et al. 2021 [1]), *Nitrospirota, Omnitrophota, Latescibacterota, Planctomycetota, Nitrospinota, Hydrogenedentota, Elusimicrobiota, Fibrobacterota, Riflebacteria, Bdellovibrionota*, UBA10199, *and Thermodesulfobacteriota* (the taxon *Deltaproteobacteria* has been reclassified into four phyla: *Thermodesulfobacteriota, Myxococcota*, SAR324, and *Bdellovibrionota* [2]) [3–5]. MTB are primarily found in stratified aquatic environments, where magnetosomes in couple with aerotaxis help cells react to environmental fluctuations [6]. That behavior is called magneto-aerotaxis or, simply, magnetotaxis [7]. The question of when did magnetotaxis emerge and how did it spread among such distant taxa is still a matter of debate. The first hypotheses concerning the origin of magnetotaxis appeared after discovering that the type of magnetosome mineral, magnetite or greigite, reflects phylogenetic affiliation. It has been suggested that magnetite-based and greigite-based magnetotaxis emerged independently [8]. Later, as data on the diversity of MTB increased, the idea of a polyphyletic origin was abandoned, especially after discovering *Desulfamplus magnetovallimortis* BW-1, capable of synthesizing crystals of both types [9–11]. One of the latest assumptions was that magnetotaxis is an ancient physiological trait with a single origin. The magnetosome gene cluster’s (MGC) evolutionary history mainly displays vertical inheritance, accompanied by multiple independent losses during bacterial diversification [4, 12]. According to the latest data, it has been suggested that a common ancestor with MGC most likely appeared before the divergence of the phyla *Nitrospirota* and *Pseudomonadota*, which are considered to be deep-branching phyla. Alternatively, it was proposed that MGC could be transferred undetectably early between the base of these phyla soon after divergence [13]. It could have happened during the mid-Archean Eon or even earlier [3, 5]. It has also been proposed that some ancient organisms could form primitive magnetosomes and, therefore, bacterial magnetotaxis would be a primal physiological process on Earth [12]. Magnetosomes could help ancient bacteria to be protected against environmental stresses on early Earth (e.g., ultraviolet radiation, toxic reactive oxygen species) [12]. This was supported by the presence of MTB in groups close to the last bacterial common ancestor (LBCA), which led to the assumption that the closest descendants of LBCA or even the last universal common ancestor (LUCA) potentially could synthesize primitive magnetosomes [5]. The lack of evidence for horizontal gene transfer (HGT) above the class level, including transfer between phyla, indeed supports the theory of vertical inheritance accompanied by multiple losses of MGC. At the same time, the evolutionary history of MGC at lower ranks is intricate [4]. For example, there is evidence for HGT before and after the delineation of *Magnetospirillum*, *Magnetovibrio*, and *Magnetospira* genera [14, 15] and within the order *Magnetococcales* [14], suggesting a more complex evolutionary history of MGC at lower taxonomic ranks.

Some magnetosome genes are essential for magnetosome formation and can be found in all MTB, while others are group-specific genes. For example, *mms* genes are found only in *Pseudomonadota, Nitrospinota*, and SAR324, *mad* genes – in *Thermodesulfobacteriota* and *Nitrospirota*, and *man* genes were initially found only in the genomes of *Nitrospirota* phylum [15]. However, recently several genomes of *Thermodesulfobacteriota* containing *man* genes were obtained [3].

In the current study, we sequenced and analyzed the genomes of the three magnetotactic bacteria previously identified in a freshwater lake Beloe Bordukovskoe [16]. As a result, we propose two novel *Candidatus* genera and three novel species of *man-genes* containing MTB. The comparative genome analysis and tree reconciliations revealed the instances of interfamily and interphylum HGT of magnetosome genes in the deep-branching phyla *Thermodesulfobacteriota* and *Nitrospirota*. Our findings contribute to understanding of the origin and evolution of MGC.

## Material and methods

### Sampling, microscopic observation, and DNA extraction

Water and sediment samples were collected from freshwater Lake Beloe Bordukovskoe to form a three-liter microcosm. MTB from this microcosm were concentrated using the MTB-CoSe approach as described in Koziaeva et al. 2020 [16]. Part of the magnetically concentrated cells was used for DNA isolation, and the rest were fixed in paraformaldehyde for morphology analyses. The DNA was used in the present study for metagenome sequencing. Fluorescence in situ hybridization combined with transmission electron microscopy (FISH-TEM) was conducted as described before [17] with specific probe LBB01 [16]. For high-resolution TEM (HRTEM), the same grids prepared for conventional TEM were imaged using a Tecnai G2 F20 FEG (FEI, USA) operated at 200 kV and equipped with a 4k×4k Gatan UltraScan 1000 CCD camera. Measurements and fast Fourier transform (FFT) from the HRTEM images were obtained using Digital Micrograph software (Gatan, USA).

### Genome sequencing, assembly, annotation, and metabolic pathways reconstructions

To obtain sufficient DNA for metagenomic sequencing, whole genome amplification was carried out using the multiple displacement amplification technique with the Genomiphi V2 DNA Amplification Kit (GE Healthcare, USA). This approach has been widely used previously in various works [18–20]. The amplified DNA was purified by sodium acetate precipitation. All stages of work with DNA were carried out according to the manufacturer’s recommendations.

For the DNA obtained after precipitation, metagenomic sequencing was performed. Short and long reads were obtained using the DNBSEQ (MGI) and Oxford Nanopore Technologies (ONT) platforms, respectively. A DNA library was constructed using the MGIEasy universal DNA library prep to obtain short reads. DNA library sequencing was performed using the DNBSEQ-G400 platform (MGI Tech, China) with pair-end 150-bp reads. Raw reads were quality checked with FastQC v.0.11.9 (http://www.bioinformatics.babraham.ac.uk/projects/fastqc/). Low-quality reads were removed using Trimmomatic v0.39 [21]. A DNA library was constructed using the NEBNext Companion Module for ONT Ligation Sequencing kit to obtain long reads. The library sequencing was performed on a MinION sequencing device (Oxford Nanopore Technologies, UK) using an R9.4.1 flow cell (FLO-MIN106D). Guppy v3.4.4, available from the Oxford Nanopore Technology community website, was used for basecalling, demultiplexing, and quality trimming ONT-passed long reads.

Long and short trimmed reads were hybrid *de novo* assembled using SPAdes v3.13.0 with the “-meta” flag [22]. Metagenome assembled genomes (MAGs) reconstruction was conducted using the MaxBin2 v2.2.7 [23], METABAT2 v2.15 [24], and Busy Bee Web [25] with standard parameters. The DAS Tool v1.1.3 was used for choosing consensus assemblies for the obtained MAGs [26]. The MAG LBB01 was manually curated and reassembled. Briefly, short and long reads were mapped to the resulting assembly using Bowtie 2 v2.3.5.110 [27] and Minimap2 v2.17 [28], respectively. The mapped reads were assembled into a circular genome with Unicycler v0.4.6 [29]. The quality metrics were assessed using the QUAST v5.0.2 [30]. The genome coverages were evaluated by QualiMap 2 v2.2.29 [31] and Bowtie 2 v2.3.2 [27]. Genome completeness and contamination were estimated using CheckM v1.1.3 [32]. RefineM v0.1.2 [33] was used to remove contamination based on taxonomic assignments. The identification of protein-coding sequences and primary annotation was performed using the NCBI Prokaryotic Genome Annotation Pipeline (PGAP v5.3) [34] and Rapid Annotations Subsystems Technology (RAST) online service [35]. The putative MGCs were determined using local BLAST and comparison with reference sequences of magnetotactic bacteria. The protein-coding sequences were annotated using the Kyoto Encyclopedia of Genes and Genomes (KEGG) framework [36]. Functional pathway prediction was performed using KEGG Mapper.

### Phylogenetic analyses and genome index calculation

The GTDB-Tk v1.6.0 [37] “classify_wf” command was used to find 120 single-copy bacterial marker protein sequences, construct their concatenated multiple alignments, and get the MAG’s taxonomic assignment using the GTDB r202 database [38]. For genome-based phylogenetic analyses, all available MTB genomes from different phyla and non-MTB *Nitrospirota* and *Thermodesulfobacteriota* genomes from the GTDB r95 database were selected (Supplementary table S1). The protein sequences of the same MGC gene (Mad26, Mad25, Mad24, Mad23, MamO-Cter, Man6, Man5, Man4, MamQ, MamE, MamI, MamA, Mad2, MamB, MamQ-2, Mad31, MamM, MamP, Man3, Mad10, Man2, MamK, Man1) from different MTB taxonomic groups were independently aligned using MAFFT [39]. PhyloSuite v1.2.2 [40] was used to concatenate MGC protein sequences. For MamA, -B, E, -I, -K, -M, -P, -Q protein sequences, and for the concatenated MGC protein sequences, the total number of protein sequences was reduced. This reduction was by taking off sequences from genomes that don’t belong to *Nitrospirota* or *Thermodesulfobacteriota* phyla. This procedure was necessary to reconcile in the Notung because the program does not work with trees with over 150 representatives. Maximum-likelihood phylogenetic trees were built with IQ-TREE v1.6.12 [41] using evolutionary models selected by ModelFinder [42]. Branch supports were obtained with 1000 ultrafast bootstraps [43]. Trees were visualized with iTOL v6.5 [44]. 120 single-copy bacterial marker protein sequences tree (hereinafter called «species tree») was rooted to *Fusobacteriota* [45]. Trees of the protein sequences of the same MGC gene (hereafter called «protein trees») were rooted to midpoint.

The average nucleotide identity (ANI) was calculated using the FastANI v1.33 tool [46]. Average amino acid identity (AAI) was calculated using CompareM v0.1.2 [47]. Digital DNA-DNA hybridization (dDDH) values were determined using Genome-to-Genome Distance Calculator (GGDC) v3.0 online software [48]. The pairwise percentage of conserved proteins (POCP) was calculated using the script runPOCP.sh [49], based on the previously described approach [50].

### Reconciliation

The evolution of proteins involved in magnetosome biogenesis was studied by reconciling protein trees and their concatenation with the species tree. Reconciliation is a method of annotating gene trees (protein trees in this work) with evolutionary events along with mapping them onto a species tree [51]. Two programs, Notung v.2.9 [52] and Ranger-DTL v2.0 [53], were used for reconciliation. Notung algorithm captures gene duplication (D), transfer (T), and loss (L) driving tree incongruence and infers all optimal solutions to finally report the complete and temporally feasible event histories giving the data. Notung was used with standard parameters: D = 1.5, T = 3, L = 1. Ranger-DTL, in turn, not only assigns one of the possible evolutionary events to nodes on the protein tree but also gives the probability of an ongoing evolutionary event, thereby refining the Notung results. Ranger-DTL analysis was run with *“Ranger-DTL”* command and 100 simulations with default parameters (D = 2, T = 3, L = 1). The *“AggregateRanger”* command was used to compute support values for the most frequent mappings that are the donor species. The reconciliation results of Notung were protein trees showing the most likely evolutionary paths taken. Ranger-DTL produced text files that indicated the most likely evolution events that may have occurred within the 100 simulations for each leaf node of the studied protein tree. Further, the results of all 100 simulations were aggregated into one resulting file. This file contained information on how many times out of 100 certain evolutionary events were found for each leaf node. Notung and Ranger-DTL reconciliations for each protein tree were recorded in a table (Supplementary table S2). Based on data from the table, the probability of a particular evolutionary event was calculated.

## Results

### Morphology of MTB cells and magnetosomes

In the previous study [16], sediment samples collected from freshwater Lake Beloe Bordukovskoe contained an abundant population of MTB with various morphology. Six MTB were identified using 16S rRNA and MamK phylogenetic analyses combined with the FISH-TEM approach. Among them, there were two *Nitrospirota* MTB (LBB01 and LBB02) and one *Thermodesulfobacteriota* MTB (LBB04) [16]. The population was dominated by a magnetotactic vibrio LBB01 allowing further analyses of its cell and magnetosome morphology using FISH-TEM.

The LBB01 probe hybridized only with vibrioid-shaped bacteria, as observed in Figures 1a-c. TEM images of the same area used for FISH analysis (Fig. 1d) revealed that this MTB group presented a thick chain of magnetosomes organized along the long axis of the bacterial cell body (Fig. 1e). The observed magnetosomes were anisotropic (Fig. 1f) and presented [111] as the elongation axis. In addition, the fast Fourier transform (FFT) pattern of crystalline structure corresponded to magnetite (Fig. 1g). LBB01 cells were 2.0 ± 0.4 μm long and 0.5 ± 0.1 μm wide (n = 32). They contained 33 ± 9 magnetosomes per cell (n = 32), which formed one bundle of bullet-shaped magnetosomes located close to each other. The bundle consisted of two to three twisted filaments of magnetosomes. The tips of the crystals were not always sequentially oriented and were sometimes oriented in opposite directions. The structure of the chains resembled that in strains MWB-1 and *Ca*. Magnetobacterium bavaricum TM-1 [54]. Immature crystals were found in different parts of the chains. A detailed analysis of the magnetosome size showed that they varied within 108 ± 21.1 nm × 45 ± 8.1 nm (length × width) with a shape factor of 0.45 (n = 1061; Supplementary Fig.S1).

**Figure 1.**
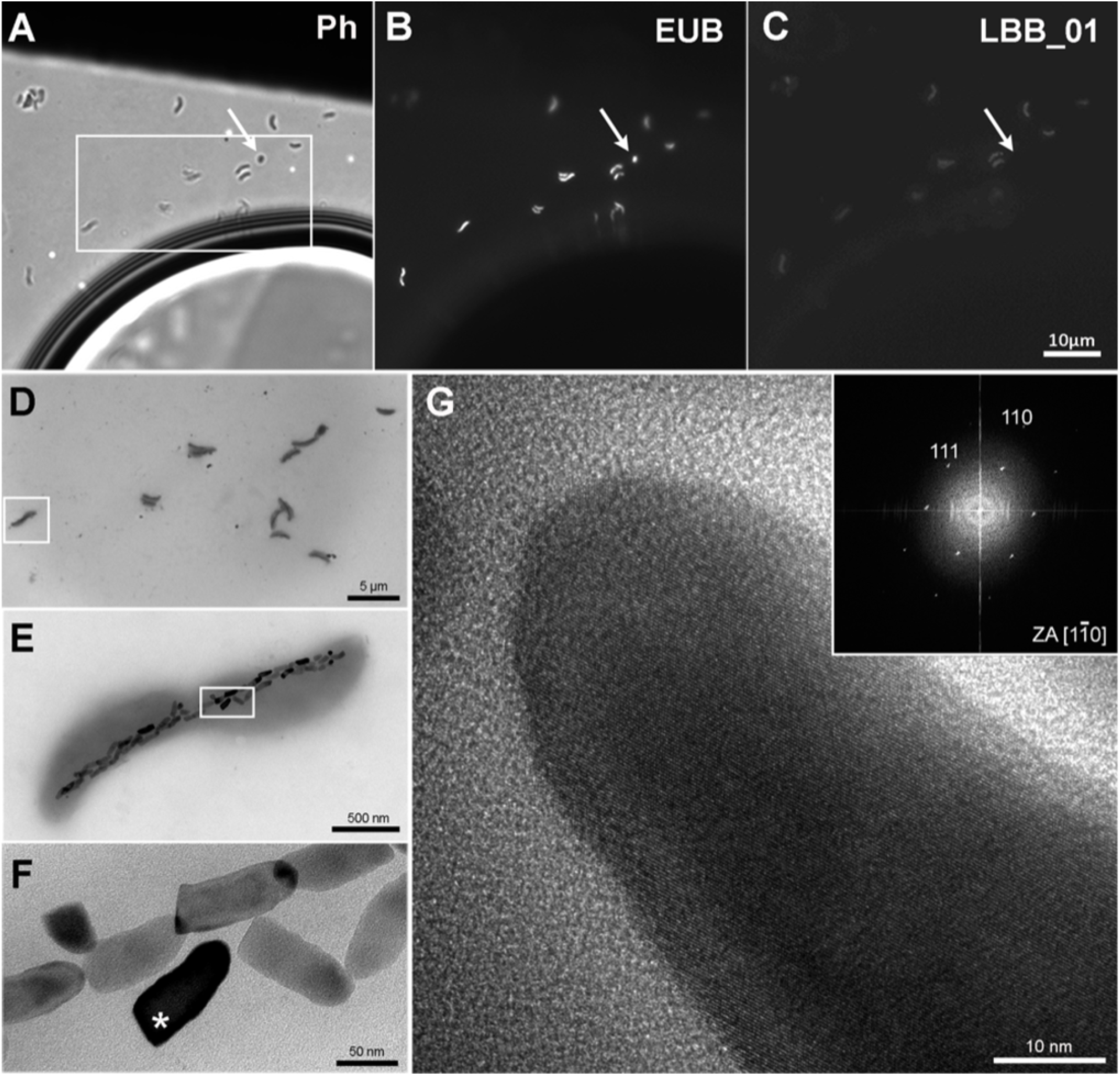
FISH-TEM of the MTB *Ca*. Magnetomonas plexicatena LBB01 and HRTEM of its magnetosomes. **A)** Phase contrast image of magnetic enrich environmental sample on Formvar coated TEM grids. **B)** Bacteria observed on image A hybridized with EUB probe. **C)** Image after hybridization with the specific design probe for *Ca*. Magnetomonas plexicatena species. The arrow highlights a small coccus present in the sample that was not hybridized with the specific probe. **D)** TEM image of the same region marked as a box in figure A. **E)** Higher magnification of the square area in figure D showing a chain of anisotropic magnetosomes organized along the long axis of the cell. **F)** Higher magnification image of the area inside the square in figure E. Magnetosomes in this cell demonstrate rough edges and a bullet shape. **G)** HRTEM of magnetosome mark with an asterisk in figure F. The FFT of the crystalline structure of the magnetosome supports that they consist of magnetite that presents an [111] elongation axis.

Besides, a small number of ovoid- and rod-shaped magnetotactic cells were present, which represented LBB02 and LBB04, respectively [16]. LBB02 were ~ 1.5 μm long and 1.2 μm wide and contained two chains of elongated magnetosomes. LBB02 is closely related to *Ca*. Magnetominusculus xianensis HCH-1, which morphology has not been identified previously [13]. Considering their high 16S rRNA identity, all representatives of *Ca*. Magnetominusculus genus might be ovoid-shaped. LBB04 is a rod-shaped cell ~ 2.5 μm long and 1.1 μm wide with disorganized elongated magnetosomes.

### Genome reconstruction and phylogenomic analyses

The genomes of the enriched MTB were assembled from metagenome sequencing. To this end, we generated 115,589,666 (2×150-bp) short paired-end (14.9 Gb) and 304,996 long reads (2.1 Gb of data, mean read length of 3,459 bp, and N_50_ of 5,251 bp). A hybrid assembly and metagenome-assembled genomes (MAG) reconstruction resulted in three MAGs that met the criteria for Genome taxonomy database (GTDB) representative genomes [38]: completeness ≥ 50% and contamination < 5%. Full or partial 16S rRNA sequences were detected, helping to link genomic data to the morphology from the previous work. First, a circular chromosome of the LBB01 genome, 3.27 Mbp long with a GC composition of 42.0%, was assembled (Supplementary table S3). Also, a high-quality draft genome for LBB02 was obtained with a GC content of 47.0% and 3.27 Mbp in size. Besides, a draft genome for LBB04, with a length of 4.49 Mbp and a GC content of 50.5%, was obtained.

According to GTDB, the reconstructed genomes of LBB01 and LBB02 were assigned to candidate family *Ca*. Magnetobacteriaceae of the phylum *Nitrospirota*, while LBB04 was assigned to the order *Syntrophales* of the *Thermodesulfobacteriota* phylum (Suppl. table S3). The phylogenomic tree based on 120 single-copy protein sequences (Fig. 2) confirmed that LBB01 and LBB02 belong to *Ca*. Magnetobacteriaceae family and LBB04 – to *Thermodesulfobacteriota* phylum. The genomes formed five clades within *Ca*. Magnetobacteriaceae, likely corresponding to five genera. One of the clades included the previously described strain *Ca*. Magnetobacterium casensis MYR-1 [55]. LBB01 genome formed a separate cluster together with the nDJH6bin1, nDJH13bin19, nDJH8bin8, and nDJH14bin5 genomes. Another clade included LBB02, *Ca*. Magnetominusculus xianensis HCH-1, nMYbin6, MYbin6, nDJH5bin4, nHCHbin2, HCHbin1, nDJH8bin6, nDJH14bin7, nDJH8bin13, and nDJH13bin15 [4, 5, 13]. The other two clades include single genomes of *Ca*. Magnetomicrobium cryptolimnococcus XYC and nDJH13bin3, respectively [56]. Two *Nitrospirota* genomes, nDJH8bin7 and nDJH14bin9, belong to a separate family. To confirm the results of the phylogenetic analyses, genomic indices were calculated. The average amino acid identity (AAI) values within the previously designated genera ranged from 75 to 100%, while values between them ranged from 55 to 62% (Supplementary table S4). It has been recently shown that there are no clear boundaries between taxa, and members of the same genus can have 65-95% AAI identity [57, 58]. Considering the branching of the phylogenomic tree and the AAI values, LBB01 can be affiliated to a separate genus, whereas LBB02 belongs to the genus *Ca*. Magnetominusculus. The percentage of orthologous conserved proteins (POCP) between genomes of the same clade was around 60% or higher, supporting the conclusions drawn from the phylogenetic and AAI analyses (Supplementary table S4). To determine taxonomy at the species level, average nucleotide identity (ANI) and digital DNA-DNA hybridization (dDDH) were calculated (Supplementary tables S5 and S6). As a result, ANI and dDDH values for LBB01 and LBB02 with closely related genomes were below the species separation threshold (< 95-96% and < 70% respectively) [59, 60], indicating that LBB01 and LBB02 represent two novel species. Therefore, we propose names *Ca*. Magnetomonas plexicatena and *Ca*. Magnetominusculus linsii for LBB01 and LBB02, respectively.

**Figure 2.**
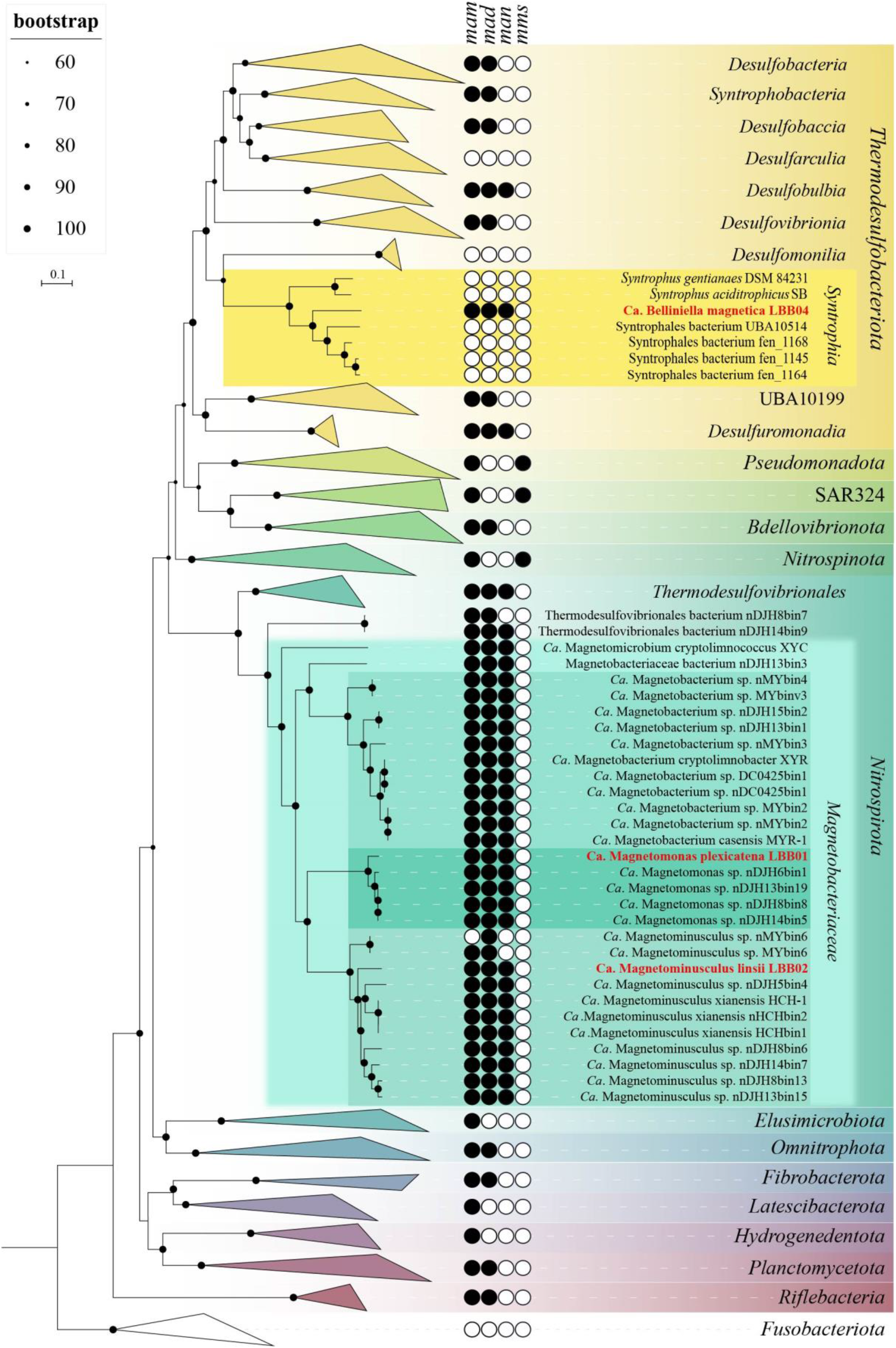
Maximum-likelihood phylogenomic tree of all previously known and the MTB genomes obtained in this work and their close non-MTB relatives. The tree was inferred from concatenated 120 bacterial single-copy marker proteins constructed with evolutionary model LG + F + I + G4. Branch supports were obtained with 1000 ultrafast bootstraps. The scale bar represents amino acid substitutions per site. The complete tree is shown in figshare data [61]. Genomes obtained in this work are highlighted in red.

As already mentioned, the genome of LBB04 clustered with representatives of the *Syntrophales* order within the *Thermodesulfobacteriota* phylum. According to the relative evolutionary divergence values and genome-based taxonomy proposed by the GTDB database, the order *Syntrophales* consists of 17 families. However, only two of them, *Syntrophaceae* and *Smithellaceae*, include validly described species. LBB04 appears phylogenetically distant from any of the known *Syntrophales* strains. The most closely related cultivated strains were *Syntrophus aciditrophicus* SB and *Syntrophus gentianae* DSM 84231 (< 94% 16S rRNA sequence similarity). Due to its distant position, we assign LBB04 to a novel genus and propose the name *Ca*. Belliniella magnetica.

### Genomic reconstruction of key metabolic functions

Genome analysis suggests that LBB01 contains the key enzymes to exploit the Wood-Ljungdahl pathway for CO2 fixation, the trait that appears common among *Nitrospirota* MTB. It has been previously hypothesized that *Ca*. Magnetobacterium casensis, a closely related species to LBB01, might utilize the reductive tricarboxylic acid cycle (rTCA) as an additional pathway for CO2 assimilation [62]. However, we found that LBB01, LBB02, *Ca*. Magnetobacterium casensis and other related *Nitrospirota* lack the key enzymes for rTCA, i.e., fumarate reductase, ATP-citrate (pro-S)-lyase (*aclAB*), and isocitrate dehydrogenase (*idh*), suggesting that Wood-Ljungdahl is likely the only CO2 fixation pathway present in these organisms. Besides, in LBB01 and LBB02, the Tricarboxylic acid cycle (TCA) appears to be incomplete as genes encoding several essential enzymes, such as succinate dehydrogenase (EC 1.3.5.1) and succinyl-CoA synthetase (EC 6.2.1.5), are not found in their genomes.

Heterotrophic utilization of glucose is possible by glycolysis (Embden-Meyerhof pathway). LBB02 contains the full set of glycolytic enzymes, whereas LBB01 lacks 6-phosphofructokinase (EC 2.7.1.11) but can alternatively utilize diphosphate-fructose-6-phosphate-1-phosphotransferase (EC 2.7.1.90) for conversion of fructose-6-phosphate to fructose-1,6-diphosphate [63].

In contrast to *Ca*. Magnetobacterium casensis, LBB01 and LBB02 appear to be unable for complete denitrification. In both organisms, the nitrate reduction to nitrite by NarGHI and nitrous oxide to nitrogen by NosZ are possible. Nitrogen fixation in LBB01 and LBB02 should not occur.

The complete set of genes associated with dissimilatory sulfur oxidation, *dsrABCHEF, aprAB*, and *sat*, was found in the genome of LBB01. Although their operation in the reverse (reductive) direction cannot be excluded entirely without the experimental evidence, the presence of *dsrEFH* is typically associated with dissimilatory sulfur oxidation. Alternatively, the pathway may be exploited to obtain energy through the disproportionation of elemental sulfur or thiosulfate, as has been recently shown for a related *Nitrospirota* bacterium [64].

Since the genome of LBB04 is highly fragmented, the reliable reconstruction of its metabolic abilities is difficult. Nonetheless, it seems to include most genes of the essential pathways (biosynthesis of amino acids, purines, pyrimidines, and several important cofactors), as well as many enzymes for TCA and glycolysis. However, LBB04 is extremely poor in genes associated with sulfur and nitrogen cycling. It may utilize acetate due to acetyl-CoA synthetase [EC:6.2.1.1] and pyruvate ferredoxin oxidoreductase [EC:1.2.7.1]. LBB04 might be able to degrade benzoate and other aromatic compounds, as it has enzymes containing benzoyl-CoA reductase domain (BcrC/BadD/HgdB). This trait is common to the cultivated *Syntrophia*, which degrade benzoate in association with hydrogen-consuming microorganisms [65].

### Magnetosome gene cluster reconstruction

Assuming the current view that MGC had vertical inheritance accompanied by many losses is true, there would be a probability of finding individual MGC genes in non-MTB genomes. Therefore, the search for MGC genes was initially carried out in all genomes of the *Nitrospirota* phylum, presented in the GTDB r95 database (Supplementary table S7), using local BLASTp and MGC protein sequences of *Ca*. Magnetomonas plexicatena LBB01 as a reference. As a result, no MGC genes have been found in non-MTB genomes.

Next, the magnetosomes synthesis genes were detected in the reconstructed MTB genomes (Supplementary Fig. S2). Considering that a circular chromosome has been assembled for *Ca*. Magnetomonas plexicatena LBB01, its MGC was taken as a reference for comparative analysis of magnetosome gene clusters containing *man* genes. The MGC of *Ca*. Magnetomonas plexicatena LBB01 consisted of three regions located on the three separate chromosome loci. The main MGC region (22 Kb) include *mad26, mad25, mad24, mad23, mamO-Cter, man6, man5, man4, mamQ, mamE, mamI, mamA, mad2, mamB, mamQ-2, mad31, mamM, mamP, man3, mad10, man2, mamK, man1* genes. A second region (2.7 Kb), located 276.8 Kb away from the main region, contains the *feoA* and *feoB* genes. These genes are homologs of the *mad30* and *mad17* genes, respectively, and are involved in iron transport for magnetosome synthesis. The third region (2.6 Kb) is located at a distance of 1.7 Mb from the second region and is composed of genes *mad28, mad29*, and a gene encoding a hypothetical protein.

The MGC of *Ca*. Magnetominusculus linsii LBB02 also includes three regions with a similar gene content to MGC of *Ca*. Magnetomonas plexicatena LBB01, except for the *mad26* gene that was located in another contig. Moreover, the gene content and synteny of the *mad26-man1* region in the assembled genomes are similar in all known magnetosome gene clusters from *Nitrospirota* phylum, confirming that the gene synteny is highly conserved in MGCs across *Nitrospirota* MTB [56]. It is also notable that two *mad28* genes were observed in the MGC of *Ca*. Magnetomonas plexicatena LBB01. Previously the duplication of this gene was detected only in *Ca*. Magnetobacterium cryptolimnobacter XYR [56]. Besides, there is a difference in MGC composition between representatives of *Ca*. Magnetobacteraceae and *Dissulfurispiraceae* families. As indicated before, the region containing the *feoAB* genes is far from the main MGC region of LBB01 that belonged to *Ca*. Magnetobacteraceae. The MGCs of the *Dissulfurispiraceae* look highly compact because the *feoAB* region is located in up to four genes from the main region. The MGC of *Ca*. Belliniella magnetica LBB04 contained only some of the genes present in *Ca*. Magnetomonas plexicatena LBB01. In the MGC of LBB04, two features attract attention. First, the *man2* and *man3* genes, previously thought to be specific for *Nitrospirota*, were detected. Second, all *mad* genes, except for *mad10*, were absent from loci where they were present in LBB01. The absence of these genes probably can be explained by the low completeness of the studied genome. However, if these genes are presented in the genome, they should occupy other loci than those in LBB01. Surprisingly, *man*-containing MGCs of other *Thermodesulfobacteriota* contained *mad2* and *mad31* genes at the same positions as in LBB01, although the same genes in the same order, like in LBB04, were presented. The part of MGCs (from *mamQ-1* to *man2*) of other *Thermodesulfobacteriota* was similar to the part of MGCs of LBB01 and other *Nitrospirota*.

### Reconstruction of the evolutionary paths for MGC of man-containing MTB

The MGCs reconstructed in this work helped investigate the evolution of MTB containing *man* genes. All *man*-containing MTB were divided into three phylogenetic groups: *Dissulfurispiraceae* group, *Magnetobacteriaceae* group, and *man*-containing *Thermodesulfobacteriota* group.

The first group, the *Dissulfurispiraceae* group, contained two families from the *Nitrospirota* phylum: the *Dissulfurispiraceae* [66] (previously named UBA9935) and the family called DUZI01 in the GTDB r202 database. According to GTDB, the *Dissulfurispiraceae* family contained ten representative genomes, four of which (nMYbin1, MYbin3, nDJH15bin8, nDJH13bin21) were affiliated to MTB (Supplementary table S1). DUZI01 family had only one genome, Nitrospirae bacterium MAG_10313_ntr_31, which belonged to MTB. The genomes of these families were combined into one group since they clustered into a separate clade on the species tree and all protein trees (figshare data “Mam_protein_trees” [61]). The *Dissulfurispiraceae* group on the species tree clustered separately from the second *man*-containing MTB group within the *Nitrospirota*, the *Magnetobacteriaceae* group. This group contained twenty-nine MTB genomes from the *Ca*. Magnetobacteriaceae family and two genomes (nDJH14bin9 and nDJH8bin7) from a new, yet unnamed in GTDB r202 database, family. As genomes of these families were clustered into a single clade on the species tree, they were combined into one group. Non-MTB representatives were not found in this group. Considering how distant *Dissulfurispiraceae* is from *Magnetobacteriaceae* on the species tree, it is crucial to understand how MGCs were inherited in the MTB of these groups. To answer this question, we conducted a reconciliation analysis. Reconciliation is a method of annotating protein trees with evolutionary events along with mapping them onto a species tree [51]. Two programs, Notung v.2.9 [52] and Ranger-DTL v2.0 [53], were used for reconciliation (see materials and methods for details). As a result, in 90.9 % of cases (20 of 22 MGC protein tree reconciliations) according to Notung and in 95.5 % (21 of 22 trees) according to Ranger-DTL, MGC genes of the *Dissulfurispiraceae* group were acquired horizontally from the *Magnetobacteriaceae* group (Supplementary Fig. S3, Supplementary table S2, figshare data “Reconciliation_results” [61]). The reconciliation results of the concatenated protein tree also confirmed the horizontal inheritance of MGC in *Dissulfurispiraceae* group from the *Magnetobacteriaceae* group. MGC from the *Magnetobacteriaceae* group was most likely transferred to the last common ancestor of the *Dissulfurispiraceae* group. This MGC was then inherited vertically by future lineages and eventually lost in the non-MTB lineages of the *Dissulfurispiraceae* group. Alternatively, in a less likely scenario, the MGC from the *Magnetobacteriaceae* group was transferred specifically to the family *Dissulfurispiraceae*, and from there, by HGT, to the family DUZI01. Thus, based on the results obtained, the MGC genes in the *Dissulfurispiraceae* group were acquired horizontally from the *Magnetobacteriaceae* group.

The reconciliation analysis also revealed that MGCs in the *Magnetobacteriaceae* group were obtained by HGT in 54.5% of cases (12 of 22 trees) according to Ranger-DTL and 59% (13 of 22 trees) according to Notung. Of all HGT cases, the most common were transfers from the *Thermodesulfobacteriota* phylum, which accounted for 46.2% (6 of 13 trees) according to Ranger-DTL and 69.2% (9 of 13 trees) according to Notung. Transfers from other phyla (*Bdellovibrionota, Riflebacteria*, *Planctomycetota*, *Omnitrophota*) were much less likely to occur. The vertical inheritance of genes from the last common *man*-containing magnetotactic ancestor was observed in 45.5% of cases (in 10 out of 22 trees) according to Ranger-DTL and 41% (9 of 22 trees) according to Notung. However, a vertical inheritance pattern was observed in protein trees containing representatives solely from *Nitrospirota* (Man1, −2, −4, −5 trees) or *Nitrospirota* and *Thermodesulfobacteriota* (Mad2, −10, −24, −25, −31 trees) phyla. Since these proteins can only be found in a limited number of taxa, their vertical inheritance results may be explained by the fact that the programs cannot analyze these evolutionary paths in sufficient detail. In the meantime, a reconciliation of the concatenated protein tree and individual Mam protein trees suggests that MTB from the *Magnetobacteriaceae* group likely acquired their MGCs by HGT.

The third group, the *Thermodesulfobacteriota* group, includes five MTB genomes affiliated to classes *Syntrophia* (LBB04), *Desulfobulbia* (MAG_13126_9_058, MAG_21600_9_004, MAG_21601_9_030), and *Desulfuromonadia* (MAG_22309_dsfv_022) from the *Thermodesulfobacteriota* phylum. It is noteworthy that MTB are rare in these classes. Despite being affiliated to different classes, investigated genomes were combined into one group because they were affiliated to one phylum. Interestingly, on the concatenated magnetosome protein trees, MTB from this group clustered with MTB affiliated with the *Nitrospirota* phylum (figshare data “Mam_protein_trees” [61]), but not with *Thermodesulfobacteriota*. Analysis of the reconciliation data revealed that MGCs of the *man*-containing *Thermodesulfobacteriota* group were acquired horizontally with 100.0% probability (19 of 19 trees) according to Ranger-DTL and 94.7% (18 of 19 trees) according to Notung. Moreover, MGCs were transferred to this group from the *Magnetobacteriaceae* group with a 73.6% probability (14 of 19 trees) according to Ranger-DTL and 52.6% (10 of 19 trees) according to Notung. The likelihood of vertical inheritance was extremely low: 0% according to Ranger-DTL and 5.3% (1 tree of 19) - to Notung. The reconciliation data of the concatenated protein tree also confirmed that the MGC in the man-containing *Thermodesulfobacteriota* group was obtained by HGT. According to Notung, this transfer originated from *Bdellovibrionota*, while Ranger-DTL showed inheritance from the *Magnetobacteriaceae* group. At the same time, within the *man*-containing *Thermodesulfobacteriota* group, it is not entirely clear how MGCs were inherited. It could be a horizontal transfer to the ancestor of all *Thermodesulfobacteriota* followed by further vertical inheritance and multiple losses of MGC in most genomes of the entire phyla. Alternatively, it could be a horizontal transfer to one specific bacterium after delineation of the major classes, followed by further transfers to other species. Since MGC horizontal gene transfer has been independently detected from the *Magentobacteriaceae* group to the *Thermodesulfobacteriota* group and vice versa, this reinforces the HGT of MGC between the two groups. (Fig. 3). This is, to our knowledge, the first detected case of inter-phylum HGT of MGC. However, more data is required for clarification, which of the groups served as a donor and which as a recipient.

**Figure 3.**
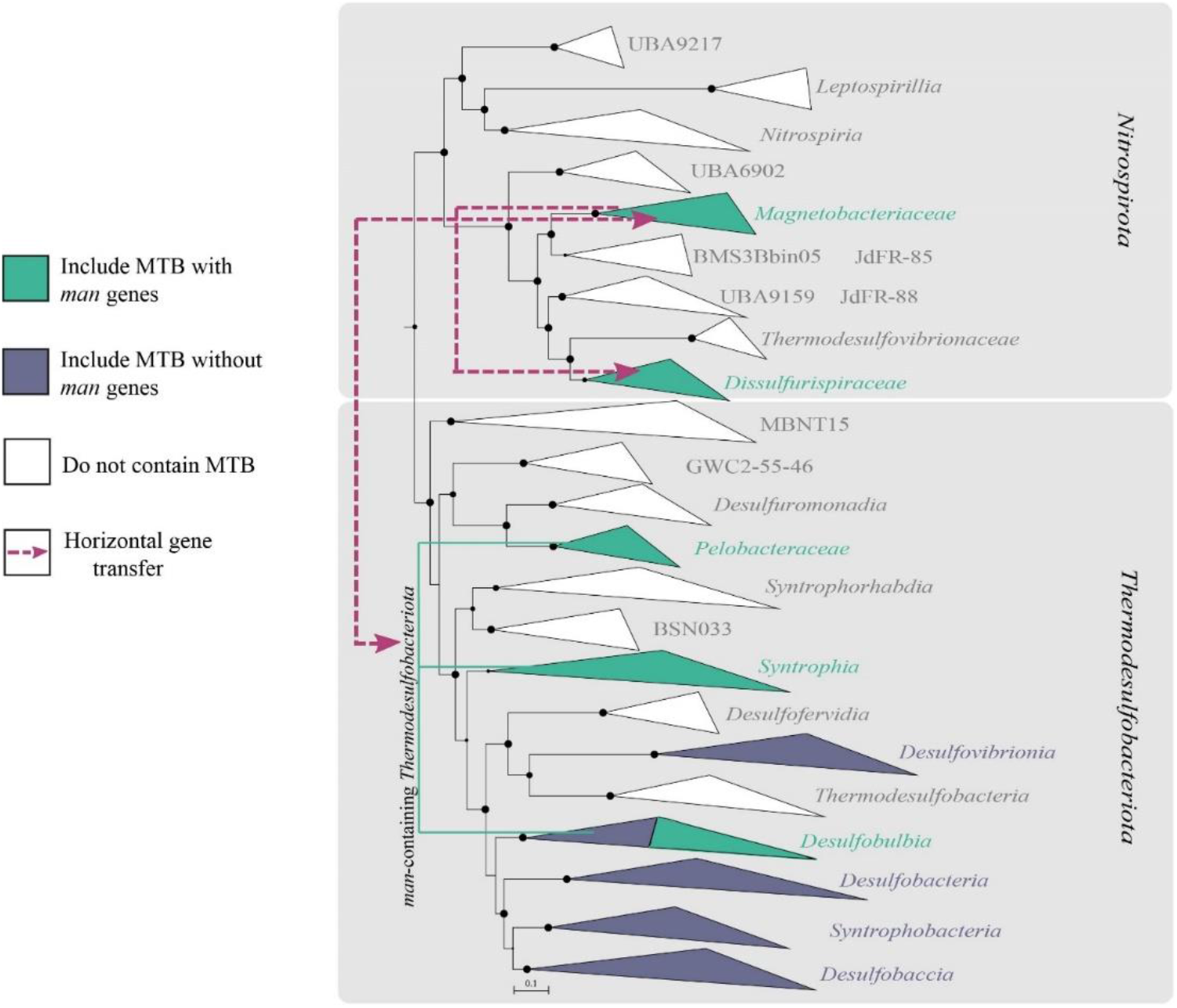
Results of reconciliation for *man*-containing genomes from the *Nitrospirota* and *Thermodesulbacteriota* phyla. A maximum likelihood phylogenomic tree was built from concatenated 120 bacterial single-copy marker proteins using evolutionary model LG + F + I + G4. Branch supports were obtained with 1000 ultrafast bootstraps. The scale bar represents amino acid substitutions per site. Branches colored in green indicate groups that include MTB representatives. Violet-colored branches include MTB representatives without *man* genes in MGCs. White branches do not have MTB members. Purple lines indicate the direction of horizontal gene transfer (HGT) of MGC.

## Discussion

Our knowledge about magnetotactic *Nitrospirota* bacteria increased significantly in the last years since *Ca*. Magnetobacterium bavaricum, the first *Nitrospirota* MTB, was discovered [67]. *Nitrospirota* representatives have been found worldwide in freshwater and marine ecosystems [68–71]. Unfortunately, there is no available pure culture of *Nitrospirota* MTB, making it difficult to accurately characterize their physiology, ecology, and magnetosome formation process. All known magnetotactic *Nitrospirota* were studied using culture-independent techniques [62, 67, 68, 70–74], and the morphology of many *Nitrospirota* MTB is known through the FISH-TEM approaches [56, 68, 70, 73]. In this study, two novel *Nitrospirota* genomes were reconstructed, which expanded MTB’s genomic diversity. For one of them, the genome of LBB01, the whole circular chromosome was successfully obtained, the first complete genome of magnetotactic *Nitrospirota*. Using the LBB01 genome as a reference allowed further comparative analyses of magnetosome genes and metabolic features. Comparative analysis showed common traits for *Nitrospirota* MTB. MGC of all genomes, with some exceptions, consisted of three separate clusters that were highly conservative, which was also noted by Zhang et al. [56]. No principal differences were found in metabolic features between the magnetotactic *Nitrospirota* genomes. Consistent with previous studies, all of the magnetotactic representatives appear to have the ability for autotrophy using Wood-Ljungdahl and heterotrophy by glycolysis via Embden-Meyerhof pathways[56, 62]. In addition, all *Nitrospirota* MTB are potentially capable of dissimilatory sulfur oxidation. At the same time, MTB that belong to *Ca*. Magnetobacteriaceae showed high diversity in cell morphology and magnetosome chain organization. So, MTB of the genus *Ca*. Magnetobacterium represent rod-shaped bacteria of various sizes that contain up to thousand magnetosomes per cell [62, 67], *Ca*. Magnetominusculus are small ovoid-shaped bacteria with several bundles of magnetosome chains, *Ca*. Magnetomonas are vibrioid-shaped bacteria with a single bundle of chains, *Ca*. Magnetoovum and *Ca*. Cryptolimnococcus are large ovoid-shaped MTB forming multiple bundles of chains [19, 73]. Accordingly, the results of phylogenetic inference supported by differences in genomic features and morphology, LBB01 and LBB02 have been defined and described as a novel genus and two novel species within *Ca*. Magnetobacteriaceae family, respectively.

Compared to *Nitrospirota*, MTB from *Thermodesulfobacteriota* phylum represented a minor fraction of the Lake Beloe Bordukovskoe MTB community. Nonetheless, we obtained a genome of strain LBB04 affiliated with the order *Syntrophales* and detected *man* genes in its MGC. Previously, *man* genes have already been detected in the MTB of *Thermodesulfobacteriota* phylum [3]. However, these genes were initially considered to be found exclusively in the *Nitrospirota* phylum [62]. The obtained new data allowed us to investigate several issues about the origin and evolution of *man*-containing MGCs.

As mentioned earlier, two theories have been proposed regarding the appearance of MGC in the MTB from the *Nitrospirota* phylum [13]. In the first and most frequently cited hypothesis, MGC in *Nitrospirota* was acquired vertically from the last common ancestor of *Nitrospirota* and *Pseudomonadota*. According to the second, less speculated view, MGC was horizontally transferred undetectably early between these two phyla soon after their delineation. These suggestions were proposed a couple of years ago when mentioned phyla clustered together on the magnetotactic phylogenetic tree. Since then, many new data on MTB diversity has emerged, providing an opportunity to explore these two suggestions in more detail.

Assuming the first suggestion about the vertical MGC inheritance is true, there should have been MGC losses within the *Nitrospirota* phylum since this phylum has many non-MTB representatives. Wang and Chen have already made this assumption during the analysis of the 16S rRNA sequences tree of the *Nitrospirota* and *Pseudomonadota* representatives known at that time [75]. However, they concluded that the number of losses must have been enormously high, which had never been documented before. Using the new genomic data, we tested this hypothesis by building the species tree (Fig. 2, figshare data [61]). On the resulting tree, MTB represented the minor part of *Nitrospirota* members, and a huge number of losses indeed should have occurred in case of the common inheritance of magnetotactic trait within this phylum. Thus, our result confirms the conclusions of Wang and Chen, also suggesting that such a scenario is unlikely to occur. Apart from that, currently, *Nitrospirota* and *Pseudomonadota* phyla are located distantly on the species tree, with many phyla that include non-MTB representatives positioned between them. In the case of a common inheritance of their MGCs, the number of losses that should have occurred increases many times, which is even more unlikely. Also, we suggest that if MGC losses within the whole *Nitrospirota* phylum did occur, it would be possible to detect the parts of the MGCs that remained after the losses. However, the search for MGC residues in non-MTB genomes of the *Nitrospirota* phylum did not reveal even a single gene for magnetosome synthesis. This may indicate that either the MGCs are always entirely removed in case of losses, or the genes were never present in these genomes. Considering all these results, we conclude that with a high probability, the last common ancestor of all *Nitrospirota* was not magnetotactic.

Next, we considered the second hypothesis, which assumes the horizontal inheritance of the *man*-containing MGC. For this, reconciliation tools were used, and statistical calculations of obtained results were carried out. Eventually, several new HGT events were detected. First, the reconciliation analysis shows that the *Magnetobacteriaceae* group donated MGC to the *Dissulfurispiraceae* group by HGT with high probability. Second, it has been found that in the *Magnetobacteriaceae* group, MGCs with a high probability were also obtained by HGT from other phyla, which, however, cannot be yet identified. The fact that *man*-containing MGCs in this group are highly conserved also provides further evidence for the recent transfer of magnetosome synthesis genes. Current results suggest that, with different probabilities, the donors for MGC could be MTB from *Thermodesulfobacteriota*, *Bdellovibrionota*, *Riflebacteria*, *Planctomycetota*, *Omnitrophota* phyla. However, none of the reconciliation results showed the probability of MGC inheritance in the *Magnetobacteriaceae* group from the MTB of the *Pseudomonadota* phylum, as suggested earlier. Within the *Magnetobacteriaceae* group, reconciliation results revealed vertical inheritance of MGCs. This is the first reported case of HGT of magnetosome synthesis genes among MTB families within the phylum *Nitrospirota*. Also, to the best of our knowledge, the first case of interphylum horizontal genes transfer of MGC was reported in this work. These results significantly refine the second, previously much less studied, theory of MGC inheritance in the *Nitrospirota* phylum.

Given the presence of *man* genes in the *Thermodesulfobacteriota* phylum, the next question was how the members of this group inherited *man*-containing MGCs. According to the reconciliation results, MGCs in the *Thermodesulfobacteriota* phylum were most likely obtained by HGT from the *Magnetobacteriaceae* group of the phylum *Nitrospirota*. Thus, our study reveals that magnetosome synthesis genes were horizontally transferred between MTB belonging to different phyla. The detection of the interphylum HGT calls into question the anciency of the origin of magnetosome synthesis genes, as the MGCs could have been transferred horizontally into these deep-branching phyla in later evolutionary periods. Although the timepoint when magnetosome synthesis emerged cannot be determined based on the currently available data, our findings suggest that it could occur after the delineation of *Nitrospirota* and *Pseudomonadota*, as at least one of these phyla acquired the MGC horizontally. This implies that the origin might be dated to a later time point than previously suggested mid-Archaeon [13]. Based on the obtained results, it can be assumed that the horizontal transfer of MGCs plays a more significant role in MTB evolution than thought previously. The possibility of interphylum transfers should be considered in further analyses of MTB evolution. Future works should focus on gathering further MTB genome sequences and more thorough reconstructions of the evolutionary history of MGC in different phylogenetic groups.

### Taxonomic consideration

#### *Candidatus* Magnetomonas

Magnetomonas (Ma.gne.to.mo’nas. Gr. n. *magnes*, - *etos* a magnet; N.L. pref. *magneto*-pertaining to a magnet; N L. fem. *n.monas* unit, monad; N.L. fem. n. *Magnetomonas* a magnetic monad)

#### *Candidatus* Magnetomonas plexicatena

Magnetomonas plexicatena (ple.xi.ca.te’na.L. past *part.plexus* interwoven; L. fem. n.*catena* chain; N.L. fem. n. *plexicatena* an interwoven chain)

Vibrioid-shaped, cell size 2.0 ± 0.4 μm long and 0.5 ± 0.1 μm wide, form magnetite magnetosomes organized in chains along the long axis of the bacterial cell body. Magnetosomes present a mean length of 108 ± 21.1 nm and a mean width of 45 ± 8.1 nm. Potentially capable of chemolithoautotrophy with the oxidation of sulfur compounds and carbon assimilation by Wood-Ljungdahl pathway. Potentially capable of heterotrophy by glycolysis. Not capable of nitrogen fixation. The reference strain is LBB01. The genome reference sequence of LBB01 is CP049016. G+C content 42.0%

#### *Candidatus* Magnetominusculus linsii

Magnetominusculus linsii (lin’si.i. N.L. gen. masc. n. *linsii*, of Lins, named after Ulysses Lins, a Brazilian microbiologist, who made a significant contribution to the study of magnetotactic bacteria)

Small ovoid cells 1.5 μm long and 1.2 μm wide, form two bundles of bullet-shaped magnetite magnetosomes. Potentially capable of chemolithoautotrophy with the oxidation of sulfur compounds and carbon assimilation by Wood-Ljungdahl pathway. Potentially capable of heterotrophy by glycolysis. Not capable of nitrogen fixation. The reference strain is LBB02. The genome reference sequence of LBB02 is JAKOEO000000000. G+C content 47.0%

#### *Candidatus* Belliniella

Belliniella (Bel.li.ni.el’la. N.L. fem. n. Belliniella, named in honour of Salvatore Bellini, an Italian microbiologist, who was one of the discoverers of magnetotactic bacteria)

#### *Candidatus* Belliniella magnetica

Belliniella magnetica (mag.ne’ti.ca. L. fem. adj. *magnetica*, of magnetic, referring to intracellular magnetite particles)

Rod-shaped cells ~ 2.5 μm long and 1.1 μm wide, form elongated magnetosomes not organized in chains. May utilize acetate. Potentially capable for glycolysis and degrading of benzoate and other aromatic compounds. The genome reference sequence of LBB04 is JAKOEP000000000. G+C content 50.4%

## Acknowledgments

We thank Prof. Aharon Oren for his expert guidance in nomenclature. Bioinformatic analyses were performed using computing resources at the Core Research Facility ‘Bioengineering’ (Research Center of Biotechnology RAS) and SciBear OU (https://sci-bear.com/). The reported study was partially funded by RFBR (project number 20-34-90116) and by the Ministry of Science and Higher Education of the Russian Federation.

## Data Availability

The genome sequences of LBB01, LBB02 and LBB04 genomes have been deposited in GenBank under the accession numbers CP049016.1, JAKOEO000000000 and JAKOEP000000000 in BioProject numbers PRJNA527025, PRJNA802353, PRJNA802356, respectively. The raw metagenomic read data have been deposited in the NCBI Sequence Read Archive under the accession numbers SRR18039296 and SRR18039297. All data generated and analyzed in this study are also available in figshare [61] and the supplementary information accompanying this paper.

