## Supplementary for "Detection of interphylum transfers of the magnetosome gene cluster in magnetotactic bacteria"

*authors with equal contribution

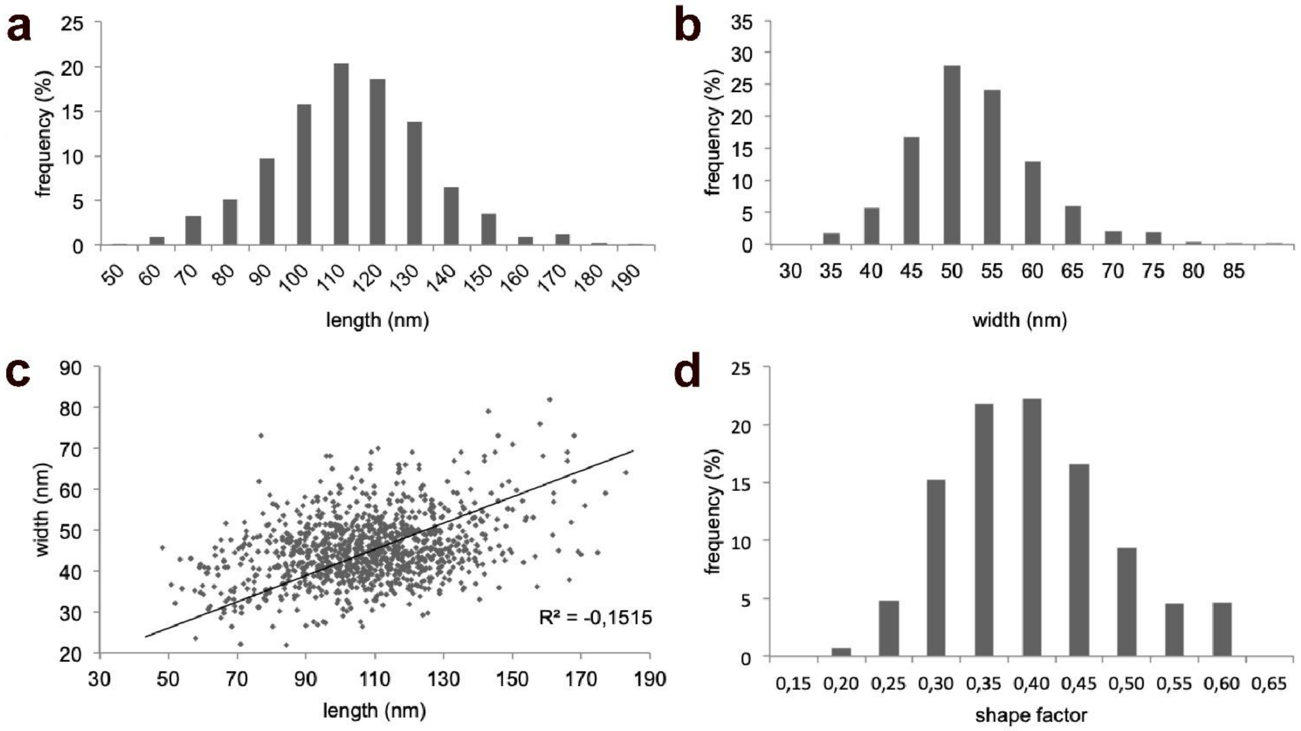

**Supplementary figure S1.** Crystal size metrics for *Candidatus* Magnetomonas plexicatena LBB01. **a** Length distribution; **b** width distribution; **c** length against width (aspect ratio); **d** shape factor distribution (n = 1061)

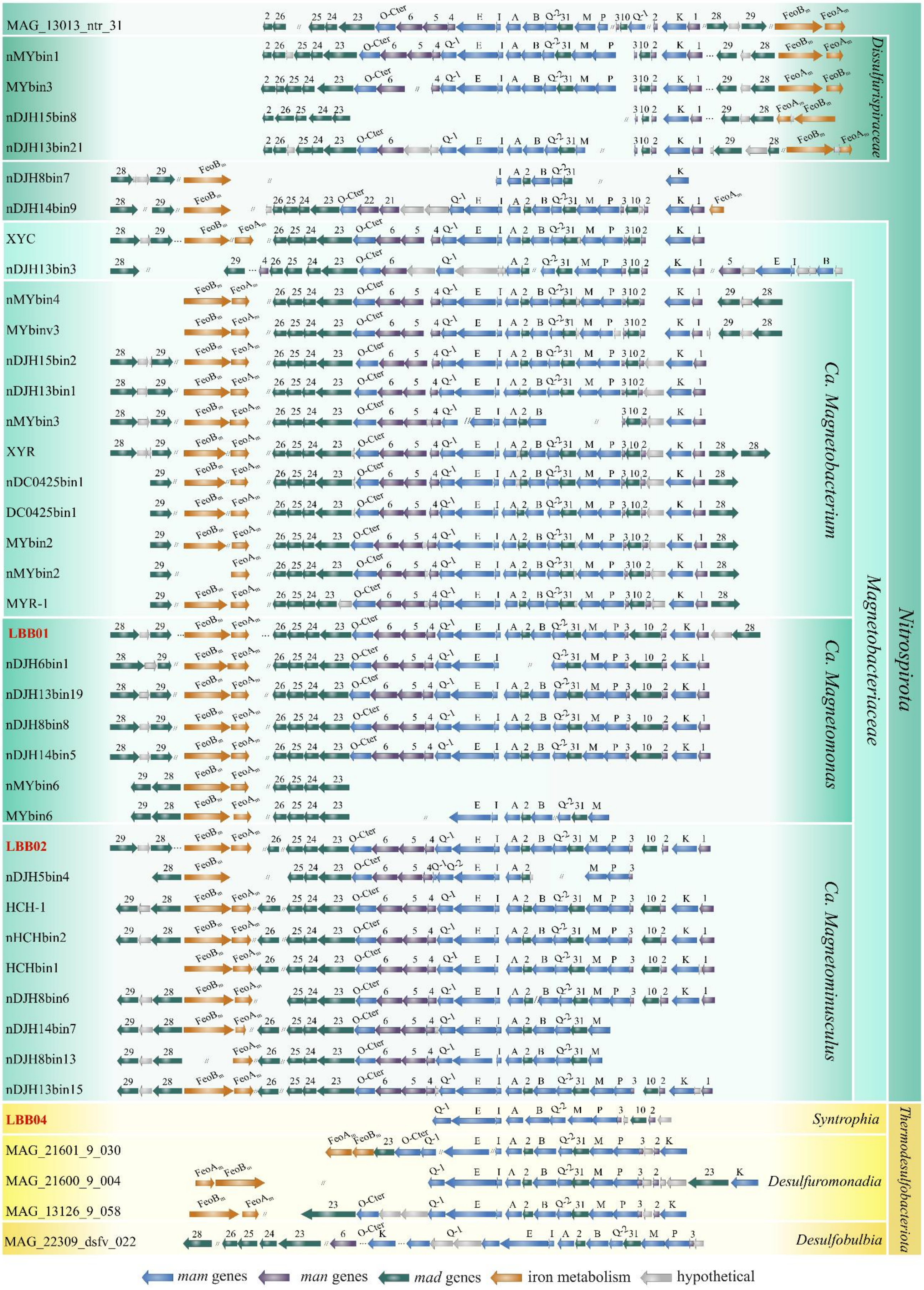

**Supplementary figure S2.** Comparison of the MGC regions in MTB genomes affiliated with the *Nitrospirota* and *Thermodesulfobacteriota* phyla. Genomes obtained in this work are highlighted in red. Full names for strains can be found in Supplementary Table S1.

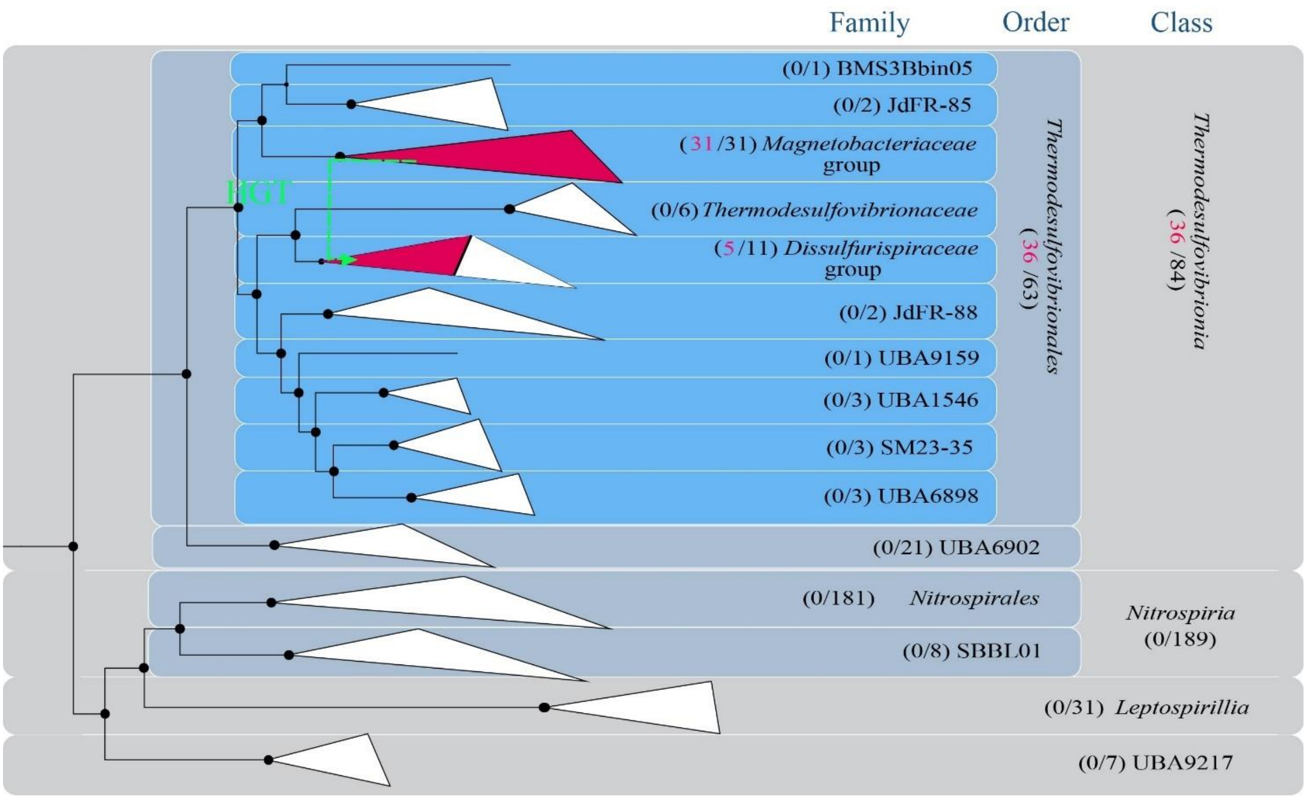

**Supplementary figure S3.** Phylogenomic analysis and reconciliation results for genomes from the *Nitrospirota* phylum including 36 MTB genomes. A maximum likelihood phylogenomic tree was inferred from concatenated 120 bacterial single-copy marker proteins, which was constructed with evolutionary model LG + F + I + G4. Branch supports were obtained with 1000 ultrafast bootstraps. The scale bar represents amino acid substitutions per site. Red-colored branches indicate groups that include MTB representatives. White-colored branches do not have MTB representatives. Green lines indicate the direction of horizontal gene transfer (HGT) of MGC

**Supplementary table S1.** Statistics for all bacterial genomes used in this work

| Genome name | NCBI/IMG accession | Size, Mbp | Scaff, no. | GC, % | GTDB Taxonomy | Completeness, % | Contamination, % |
|---|---|---|---|---|---|---|---|
| Alphaproteobacteria bacterium nDJH14bin10 | GCA_015233585.1 | 3.33 | 435 | 60.2 | d__Bacteria;p__Proteobacteria;c__Alphaproteobacteria;o__Rhodospirillales;f__WOUV01;g__;s__ | 96.63 | 0.00 |
| Alphaproteobacteria bacterium nJC1bin3 | GCA_015232485.1 | 3.96 | 698 | 67.27 | d__Bacteria;p__Proteobacteria;c__Alphaproteobacteria;o__Rhodospirillales;f__Magnetospirillaceae;g__;s__ | 86.90 | 2.94 |
| Alphaproteobacteria bacterium nJC2bin54 | GCA_015232435.1 | 3.89 | 694 | 67.6 | d__Bacteria;p__Proteobacteria;c__Alphaproteobacteria;o__Rhodospirillales;f__Magnetospirillaceae;g__;s__ | 97.48 | 2.10 |
| Alphaproteobacteria bacterium nKLKbin1 | GCA_015232335.1 | 3.43 | 89 | 59.34 | d__Bacteria;p__Proteobacteria;c__Alphaproteobacteria;o__Rhodospirillales;f__WMHbin7;g__WMHbin7;s__ | 99.50 | 0.00 |
| Alphaproteobacteria bacterium nN2-2bin1 | GCA_015232095.1 | 4.33 | 129 | 65.17 | d__Bacteria;p__Proteobacteria;c__Alphaproteobacteria;o__Rhodospirillales;f__Magnetospirillaceae;g__Magnetospirillum;s__Magnetospirillum moscoviense | 99.50 | 0.50 |
| Alphaproteobacteria b`acterium nN2-2bin2 | GCA_015232055.1 | 4.10 | 602 | 67.51 | d__Bacteria;p__Proteobacteria;c__Alphaproteobacteria;o__Rhodospirillales;f__Magnetospirillaceae;g__;s__ | 96.97 | 1.82 |
| Alphaproteobacteria bacterium nN3bin14 | GCA_015232025.1 | 3.74 | 499 | 67.66 | p__Proteobacteria;c__Alphaproteobacteria;o__Rhodospirillales;f__;g__;s__ | 61.19 | 2.10 |
| Alphaproteobacteria bacterium nSSYDbin4 | GCA_015231605.1 | 3.18 | 213 | 63.34 | p__Proteobacteria;c__Alphaproteobacteria;o__Rhodospirillales;f__Magnetovibrionaceae;g__;s__ | 78.78 | 1.58 |
| Alphaproteobacteria bacterium nTSbin12 | GCA_015231505.1 | 2.39 | 329 | 63.33 | p__Proteobacteria;c__Alphaproteobacteria;o__Rhodospirillales;f__Magnetovibrionaceae;g__;s__ | 78.43 | 0.00 |
| Alphaproteobacteria bacterium nWMHbin5 | GCA_015229215.1 | 3.56 | 80 | 59.35 | p__Proteobacteria;c__Alphaproteobacteria;o__Rhodospirillales;f__WMHbin7;g__WMHbin7;s__WMHbin7 sp002753155 | 99.84 | 7.58 |
| Alphaproteobacteria bacterium nWRX3bin4 | GCA_015228915.1 | 2.59 | 411 | 66.09 | p__Proteobacteria;c__Alphaproteobacteria;o__Rhodospirillales;f__WOUV01;g__;s__ | 97.66 | 0.00 |
| Alphaproteobacteria bacterium nYQH56bin3 | GCA_015228695.1 | 2.80 | 436 | 65.57 | p__Proteobacteria;c__Alphaproteobacteria;o__Rhodospirillales;f__WOUV01;g__;s__ | 97.17 | 0.86 |
| Alphaproteobacteria bacterium WMHbin7 | GCA_002753155.1 | 2.98 | 73 | 59.84 | p__Proteobacteria;c__Alphaproteobacteria;o__Rhodospirillales;f__WMHbin7;g__WMHbin7;s__WMHbin7 sp002753155 | 96.43 | 3.87 |
| Bilophila wadsworthia ATCC 49260 | GCF_000701705.1 | 4.63 | 198 | 59.25 | p__Desulfobacterota;c__Desulfovibrionia;o__Desulfovibrionales;f__Desulfovibrionaceae;g__Bilophila;s__Bilophila wadsworthia | 93.15 | 3.04 |
| Caenispirillum salinarum AK4 | GCF_000315795.1 | 4.95 | 61 | 68.79 | p__Proteobacteria;c__Alphaproteobacteria;o__Rhodospirillales;f__Rhodospirillaceae;g__Caenispirillum;s__Caenispirillum salinarum | 55.90 | 0.00 |
| Caldimicrobium thiodismutans TF1 | GCF_001548275.1 | 1.81 | 1 | 38.3 | p__Desulfobacterota;c__Thermodesulfobacteria;o__Thermodesulfobacteriales;f__Thermodesulfobacteriaceae;g__Caldimicrobium;s__Caldimicrobium thiodismutans | 99.95 | 1.29 |
| Caldimicrobium thiodismutans ZAV-15 | GCA_002877875.1 | 1.30 | 103 | 36.9 | p__Desulfobacterota;c__Thermodesulfobacteria;o__Thermodesulfobacteriales;f__Thermodesulfobacteriaceae;g__Caldimicrobium;s__Caldimicrobium thiodismutans_A | 84.30 | 3.23 |
| Ca. Adiutrix intracellularis Adiu1 | GCA_001577715.1 | 2.08 | 155 | 43.26 | p__Desulfobacterota;c__Desulfarculia_A;o__Adiutricales;f__Adiutricaceae;g__Adiutrix;s__Adiutrix intracellularis | 73.45 | 2.14 |
| Ca. Desulfofervidus auxilii HS1 | GCF_001577525.1 | 2.54 | 1 | 37.17 | p__Desulfobacterota;c__Desulfofervidia;o__Desulfofervidales;f__Desulfofervidaceae;g__Desulfofervidus;s__Desulfofervidus auxilii | 83.53 | 3.23 |
| Ca. Electrothrix aarhusiensis MCF | GCA_004028505.1 | 3.73 | 143 | 47.47 | p__Desulfobacterota;c__Desulfobulbia;o__Desulfobulbales;f__Desulfobulbaceae;g__Electrothrix;s__Electrothrix aarhusiensis | 58.77 | 1.75 |
| Ca. Electrothrix marina A5 | GCA_004028495.1 | 2.06 | 472 | 49.7 | p__Desulfobacterota;c__Desulfobulbia;o__Desulfobulbales;f__Desulfobulbaceae;g__Electrothrix;s__Electrothrix marina | 91.29 | 3.71 |
| Ca. Endomicrobium trichonymphae Rs-D17 | GCA_002355835.1 | 1.13 | 4 | 35.23 | p__Elusimicrobiota;c__Endomicrobia;o__Endomicrobiales;f__Endomicrobiaceae;g__Endomicrobium_A;s__Endomicrobium_A trichonymphae | 90.32 | 0.65 |
| Ca. Hydrogenedens terephthalicus JGI OTU-1 | GCA_000493945.1 | 2.87 | 106 | 36.9 | p__Hydrogenedentota;c__Hydrogenedentia;o__Hydrogenedentiales;f__Hydrogenedentaceae;g__Hydrogenedens;s__Hydrogenedens terephthalicus | 49.08 | 0.97 |
| Ca. Hydrogenedentes bacterium MAG_17963_hgd_111 | GCA_013349485.1 | 3.02 | 288 | 60.18 | p__Hydrogenedentota;c__Hydrogenedentia;o__Hydrogenedentiales;f__DUZN01;g__DUZN01;s__DUZN01 sp013349485 | 67.25 | 2.56 |
| Ca. Hydrogenedentes bacterium MAG_17971_hgd_130 | GCA_013349535.1 | 2.68 | 240 | 60.43 | p__Hydrogenedentota;c__Hydrogenedentia;o__Hydrogenedentiales;f__DUZN01;g__DUZN01;s__DUZN01 sp013349535 | 90.20 | 3.64 |
| Ca. Hydrogenedentes bacterium NORP43 | GCA_002746185.1 | 4.23 | 202 | 49.96 | p__Hydrogenedentota;c__Hydrogenedentia;o__Hydrogenedentiales;f__GCA-2746185;g__GCA-2746185;s__GCA-2746185 sp002746185 | 89.44 | 1.24 |
| Ca. Hydrogenedentes bacterium UBA6118 | GCA_002422525.1 | 4.51 | 94 | 67.12 | p__Hydrogenedentota;c__Hydrogenedentia;o__Hydrogenedentiales;f__SLHB01;g__UBA6118;s__UBA6118 sp002422525 | 89.59 | 2.53 |
| Ca. Lambdaproteobacteria bacterium PCRbin3 | GCA_002753255.1 | 5.06 | 530 | 41.83 | p__SAR324;c__SAR324;o__SAR324;f__GCA-2753255;g__GCA-2753255;s__GCA-2753255 sp002753255 | 97.67 | 2.73 |
| Ca. Lambdaproteobacteria bacterium RIFOXYC1_FULL_56_13 | GCA_001783655.1 | 3.09 | 66 | 56.53 | p__SAR324;c__SAR324;o__XYD2-FULL-50-16;f__XYD2-FULL-50-16;g__XYC1-FULL-56-13;s__XYC1-FULL-56-13 sp001783715 | 100.00 | 0.97 |
| Ca. Lambdaproteobacteria bacterium RIFOXYD2_FULL_50_16 | GCA_001783695.1 | 3.14 | 54 | 49.65 | p__SAR324;c__SAR324;o__XYD2-FULL-50-16;f__XYD2-FULL-50-16;g__XYD2-FULL-50-16;s__XYD2-FULL-50-16 sp001783695 | 98.71 | 2.26 |
| Ca. Latescibacter anaerobius SCGC AAA252-E07 | 2264867254 | 2.29 | 164 | 42.12 | p__Latescibacterota;c__Latescibacteria;o__Latescibacterales;f__Latescibacteraceae;g__Latescibacter;s__Latescibacter anaerobius | 57.63 | 1.10 |
| Ca. Latescibacteria bacterium 4484_107 | GCA_002059205.1 | 1.60 | 192 | 55.50 | p__Latescibacterota;c__Latescibacteria_A;o__4484-107;f__4484-107;g__4484-107;s__4484-107 sp002059205 | 76.68 | 0.00 |
| Ca. Latescibacteria bacterium 4484_181 | GCA_002049985.1 | 1.00 | 102 | 49.86 | p__Latescibacterota;c__MVCY01;o__MVCY01;f__MVCY01;g__MVCY01;s__MVCY01 sp002049985 | 76.91 | 0.25 |
| Ca. Latescibacteria bacterium 4484_7 | GCA_002085285.1 | 1.53 | 161 | 50.94 | p__Krumholzibacteriota;c__Krumholzibacteria;o__Krumholzibacteriales;f__Krumholzibacteriaceae;g__4484-7;s__4484-7 sp002085285 | 69.78 | 2.20 |
| Ca. Magnetaquicoccus inordinatus UR-1 | GCA_004217665.1 | 4.14 | 546 | 52.51 | p__Proteobacteria;c__Magnetococcia;o__Magnetococcales;f__Magnetaquicoccaceae;g__Magnetaquicoccus;s__Magnetaquicoccus inordinatus | 99.50 | 1.00 |
| Ca. Magnetobacterium casensis MYR-1 | GCF_000714715.1 | 3.42 | 70 | 48.87 | p__Nitrospirota;c__Thermodesulfovibrionia;o__Thermodesulfovibrionales;f__Magnetobacteriaceae;g__Magnetobacterium;s__Magnetobacterium casensis | 95.76 | 1.36 |
| Ca. Magnetobacterium sp. XYC | GCA_018606805.1 | 3.59 | 91 | 37.75 | p__Nitrospirota;c__Thermodesulfovibrionia;o__Thermodesulfovibrionales;f__Magnetobacteriaceae;g__;s__ | 92.94 | 0.42 |
| Ca. Magnetobacterium sp. XYR | GCA_018606785.1 | 4.23 | 195 | 48.61 | p__Nitrospirota;c__Thermodesulfovibrionia;o__Thermodesulfovibrionales;f__Magnetobacteriaceae;g__Magnetobacterium;s__Magnetobacterium sp002753685 | 73.26 | 1.37 |
| Ca. Magnetomorum sp nER2bin1 | GCA_015232875.1 | 6.16 | 393 | 38.94 | p__Desulfobacterota;c__Desulfobacteria;o__Desulfobacterales;f__Magnetomoraceae;g__Magnetomorum;s__Magnetomorum sp002753725 | 90.97 | 3.55 |
| Ca. Magnetomorum sp. HK-1 | GCA_001292585.1 | 14.15 | 3036 | 34.61 | p__Desulfobacterota;c__Desulfobacteria;o__Desulfobacterales;f__Magnetomoraceae;g__Magnetomorum;s__ | 90.31 | 4.20 |
| Ca. Nitrospira inopinata ENR4 | GCF_001458695.1 | 3.30 | 1 | 59.23 | p__Nitrospirota;c__Nitrospiria;o__Nitrospirales;f__Nitrospiraceae;g__Nitrospira_F;s__Nitrospira_F inopinata | 78.32 | 0.18 |
| Ca. Nitrospira nitrificans COMA2 | GCF_001458775.1 | 4.12 | 36 | 56.59 | p__Nitrospirota;c__Nitrospiria;o__Nitrospirales;f__Nitrospiraceae;g__Nitrospira_F;s__Nitrospira_F nitrificans | 97.48 | 2.94 |
| Ca. Omnitrophica bacterium Cal1bin1 | GCA_002753745.1 | 2.54 | 240 | 49.55 | p__Omnitrophota;c__Koll11;o__UBA10015;f__GCA-002753745;g__GCA-2753745;s__GCA-2753745 sp002753745 | 76.15 | 3.23 |
| Ca. Omnitrophica bacterium MBPbin6 | GCA_002753465.1 | 2.18 | 150 | 49.49 | p__Omnitrophota;c__Koll11;o__UBA10015;f__GCA-002753745;g__GCA-2753465;s__GCA-2753465 sp002753465 | 94.84 | 0.83 |
| Ca. Omnitrophica bacterium nCal1bin2 | GCA_015233765.1 | 2.95 | 196 | 49.66 | p__Omnitrophota;c__Koll11;o__UBA10015;f__GCA-002753745;g__GCA-2753745;s__GCA-2753745 sp002753745 | 66.90 | 1.72 |
| Ca. Omnitrophica bacterium nCal2bin1 | GCA_015234065.1 | 1.75 | 176 | 51.48 | p__Omnitrophota;c__Koll11;o__UBA10015;f__GCA-002753745;g__;s__ | 82.80 | 3.23 |
| Ca. Omnitrophica bacterium nDJH13bin13 | GCA_015233705.1 | 1.17 | 97 | 54.15 | p__Omnitrophota;c__Koll11;o__UBA10015;f__GCA-002753745;g__;s__ | 95.77 | 0.00 |
| Ca. Omnitrophica bacterium nDJH13bin20 | GCA_015233675.1 | 2.12 | 31 | 42.8 | p__Omnitrophota;c__Koll11;o__UBA10015;f__GCA-002753745;g__JABDGZ01;s__ | 69.64 | 0.00 |
| Ca. Omnitrophica bacterium nDJH15bin13 | GCA_015233495.1 | 2.10 | 111 | 49.08 | p__Omnitrophota;c__Koll11;o__UBA10015;f__Kpj58rc;g__UBA12451;s__ | 97.32 | 0.00 |
| Ca. Omnitrophica bacterium nDJH15bin6 | GCA_015233405.1 | 2.38 | 47 | 43.25 | p__Omnitrophota;c__Koll11;o__UBA10015;f__Kpj58rc;g__;s__ | 66.54 | 0.97 |
| Ca. Omnitrophica bacterium nDJH2bin13 | GCA_015233375.1 | 1.76 | 209 | 42.08 | p__Omnitrophota;c__Koll11;o__UBA10015;f__GCA-002753745;g__;s__ | 81.25 | 0.65 |
| Ca. Omnitrophica bacterium nDJH2bin18 | GCA_015233335.1 | 1.85 | 38 | 44.09 | p__Omnitrophota;c__Koll11;o__UBA10015;f__GCA-002753745;g__JABDGZ01;s__ | 95.17 | 1.97 |
| Ca. Omnitrophica bacterium nDJH6bin13 | GCA_015233235.1 | 1.59 | 175 | 47.17 | p__Omnitrophota;c__Koll11;o__UBA1560;f__SKK-01;g__;s__ | 95.95 | 0.91 |
| Ca. Omnitrophica bacterium nDJH6bin14 | GCA_015233195.1 | 1.98 | 50 | 37.49 | p__Omnitrophota;c__Koll11;o__UBA10015;f__Kpj58rc;g__UBA12451;s__ | 70.95 | 0.00 |
| Ca. Omnitrophica bacterium nDJH6bin18 | GCA_015233165.1 | 2.29 | 187 | 41.64 | p__Omnitrophota;c__Koll11;o__UBA10015;f__GCA-002753745;g__;s__ | 71.27 | 1.82 |
| Ca. Omnitrophica bacterium nDJH6bin28 | GCA_015233145.1 | 1.81 | 58 | 47.32 | p__Omnitrophota;c__Koll11;o__UBA10015;f__GCA-002753745;g__;s__ | 98.99 | 3.99 |

| Genome name | NCBI/IMG accession | Size, Mbp | Scaff, no. | GC, % | GTDB Taxonomy | Completeness, % | Contamination, % |
|---|---|---|---|---|---|---|---|
| Ca. Omnitrophica bacterium nDJH6bin5 | GCA_015233115.1 | 2.39 | 37 | 40.68 | p_Omnitrophota;c_Koll11;o_UBA10015;f_Kpj58rc;g_UBA12451;s_ | 64.82 | 1.82 |
| Ca. Omnitrophica bacterium nHGRbin18 | GCA_015232565.1 | 2.64 | 80 | 51.11 | p_Omnitrophota;c_Koll11;o_UBA10015;f_GCA-002753745;g_GCA-2753745;s_ | 77.53 | 1.69 |
| Ca. Omnitrophica bacterium nHLHbin2 | GCA_015232535.1 | 2.21 | 210 | 42.78 | p_Omnitrophota;c_Koll11;o_UBA10015;f_Kpj58rc;g_;s_ | 67.24 | 0.65 |
| Ca. Omnitrophica bacterium nMBPbin3 | GCA_015232235.1 | 1.97 | 41 | 50.38 | p_Omnitrophota;c_Koll11;o_UBA10015;f_GCA-002753745;g_GCA-2753465;s_GCA-2753465 sp002753465 | 65.52 | 0.00 |
| Ca. Omnitrophica bacterium nS315bin20 | GCA_015231685.1 | 1.38 | 200 | 56.96 | p_Omnitrophota;c_Koll11;o_2-02-FULL-51-18;f_;g_;s_ | 98.03 | 0.91 |
| Ca. Omnitrophica bacterium nS315bin24 | GCA_015231715.1 | 1.50 | 115 | 37.11 | p_Omnitrophota;c_Koll11;o_UBA1560;f_SKK-01;g_;s_ | 92.02 | 0.91 |
| Ca. Omnitrophica bacterium nW3bin14 | GCA_015231405.1 | 3.66 | 93 | 49.49 | p_Omnitrophota;c_Koll11;o_UBA1560;f_SKK-01;g_;s_ | 79.69 | 2.15 |
| Ca. Omnitrophica bacterium nXXbin4 | GCA_015228785.1 | 2.02 | 123 | 46.42 | p_Omnitrophota;c_Koll11;o_UBA10015;f_GCA-002753745;g_JAAYZZ01;s_ | 73.12 | 0.10 |
| Ca. Omnitrophus fodinae SCGC AAA011-A17 | GCA_000405945.1 | 2.04 | 150 | 49.56 | p_Omnitrophota;c_Omnitrophia;o_Omnitrophales;f_Omnitrophaceae;g_Omnitrophus;s_Omnitrophus fodinae | 83.33 | 1.08 |
| Ca. Riflebacteria bacterium nDJH14bin3 | GCA_015233565.1 | 8.30 | 185 | 40.61 | p_Riflebacteria;c_Ozemobacteria;o_Ozemobacterales;f_Ozemobacteraceae;g_;s_ | 98.70 | 2.37 |
| Ca. Riflebacteria bacterium nDJH6bin19 | GCA_015233155.1 | 6.31 | 185 | 49.32 | p_Riflebacteria;c_Ozemobacteria;o_Ozemobacterales;f_Ozemobacteraceae;g_;s_ | 99.91 | 0.60 |
| Ca. Riflebacteria bacterium nHGRbin4 | GCA_015232575.1 | 7.05 | 54 | 40.28 | p_Riflebacteria;c_Ozemobacteria;o_Ozemobacterales;f_Ozemobacteraceae;g_;s_ | 98.06 | 0.00 |
| Ca. Terasakiella magnetica PR-1 | GCF_900093605.1 | 3.68 | 48 | 45.97 | p_Proteobacteria;c_Alphaproteobacteria;o_Rhodospirillales;f_Terasakiellaceae;g_Terasakiella;s_Terasakiella magnetica_A | 89.26 | 3.64 |
| delta proteobacterium MLS_D | GCA_002030015.1 | 2.51 | 70 | 55.2 | p_Desulfobacterota;c_Syntrophia;o_Syntrophales;f_UBA2210;g_MLS-D;s_MLS-D sp002030015 | 96.76 | 3.18 |
| Delta proteobacterium NaphS2 | GCA_000179315.1 | 6.55 | 810 | 49.83 | p_Desulfobacterota;c_Desulfobacteria;o_Desulfatiglandales;f_Desulfatiglandaceae;g_NaphS2;s_NaphS2 sp000179315 | 97.85 | 0.00 |
| Deltaproteobacteria bacterium 37-65-8 | GCA_002279275.1 | 1.83 | 193 | 65.33 | p_Desulfobacterota_E;c_MBNT15;o_MBNT15;f_MBNT15;g_CG2-30-66-27;s_CG2-30-66-27 sp002279275 | 70.32 | 0.65 |
| Deltaproteobacteria bacterium B111_G9 | GCA_003646995.1 | 3.85 | 542 | 47.56 | p_Desulfobacterota;c_Desulfobacteria;o_Desulfatiglandales;f_Desulfatiglandaceae;g_B111-G9;s_B111-G9 sp003646995 | 61.93 | 0.33 |
| Deltaproteobacteria bacterium B119_G9 | GCA_003646715.1 | 2.74 | 559 | 43.47 | p_Desulfobacterota;c_Syntrophobacteria;o_B119-G9;f_B119-G9;g_B119-G9;s_B119-G9 sp003646715 | 99.50 | 0.50 |
| Deltaproteobacteria bacterium B13_G4 | GCA_003647525.1 | 1.69 | 215 | 39.2 | p_Desulfobacterota;c_Desulfobacteria;o_Desulfobacterales;f_ETH-SRB1;g_B13-G4;s_B13-G4 sp003647525 | 64.43 | 2.58 |
| Deltaproteobacteria bacterium B17_G16 | GCA_003646975.1 | 3.25 | 554 | 58.82 | p_Desulfobacterota;c_Desulfobacteria;o_Desulfatiglandales;f_Desulfatiglandaceae;g_B17-G16;s_B17-G16 sp003646975 | 91.59 | 0.00 |
| Deltaproteobacteria bacterium B25_G16 | GCA_003646935.1 | 2.38 | 429 | 50.52 | p_Desulfobacterota;c_Desulfobacteria;o_Desulfatiglandales;f_B25-G16;g_B25-G16;s_B25-G16 sp003646935 | 99.87 | 0.89 |
| Deltaproteobacteria bacterium B3_G2 | GCA_003647385.1 | 3.07 | 611 | 41.15 | p_Desulfobacterota;c_Desulfobacteria;o_Desulfobacterales;f_Desulfobacteraceae;g_Desulfobacula;s_Desulfobacula sp003647385 | 91.29 | 0.75 |
| Deltaproteobacteria bacterium B30_G6 | GCA_003647375.1 | 5.41 | 703 | 47.76 | p_Desulfobacterota;c_Desulfobacteria;o_Desulfobacterales;f_B30-G6;g_B30-G6;s_B30-G6 sp003647375 | 72.33 | 2.58 |
| Deltaproteobacteria bacterium B33_G16 | GCA_003646875.1 | 2.20 | 384 | 41.86 | p_Desulfobacterota;c_Desulfobacteria;o_Desulfatiglandales;f_B25-G16;g_B33-G16;s_B33-G16 sp003646875 | 96.94 | 3.57 |
| Deltaproteobacteria bacterium B37_G16 | GCA_003646865.1 | 2.47 | 349 | 48.37 | p_Desulfobacterota;c_Desulfobacteria;o_Desulfatiglandales;f_Desulfatiglandaceae;g_B111-G9;s_B111-G9 sp003646865 | 85.66 | 6.77 |
| Deltaproteobacteria bacterium B46_G9 | GCA_003646815.1 | 3.47 | 410 | 51.5 | p_Desulfobacterota;c_Desulfobacteria;o_Desulfatiglandales;f_Desulfatiglandaceae;g_B46-G9;s_B46-G9 sp003646815 | 78.99 | 0.84 |
| Deltaproteobacteria bacterium CG_4_8_14_3_um_filter_43_13 | GCA_002782605.1 | 2.57 | 311 | 43.25 | p_Desulfobacterota;c_SM23-61;o_UBA8473;f_UBA8473;g_UBA8473;s_UBA8473 sp002782605 | 88.06 | 0.70 |
| Deltaproteobacteria bacterium CG03_land_8_20_14_0_80_45_14 | GCA_002780715.1 | 3.24 | 253 | 45.5 | p_Desulfobacterota;c_BSN033;o_BSN033;f_UBA1163;g_UBA1163;s_UBA1163 sp002780715 | 94.52 | 2.39 |
| Deltaproteobacteria bacterium CG07_land_8_20_14_0_80_60_11 | GCA_002779455.1 | 2.42 | 285 | 59.82 | p_Desulfobacterota;c_Desulfobaccia;o_Desulfobaccales;f_0-14-0-80-60-11;g_0-14-0-80-60-11;s_0-14-0-80-60-11 sp002779455 | 100.00 | 0.00 |
| Deltaproteobacteria bacterium CSP1-8 | GCA_001443455.1 | 2.18 | 46 | 63.49 | p_Desulfobacterota_E;c_MBNT15;o_MBNT15;f_MBNT15;g_CSP1-8;s_CSP1-8 sp001443455 | 95.45 | 2.00 |
| Deltaproteobacteria bacterium D1FN1.001 | GCA_005774545.1 | 3.10 | 115 | 65.26 | p_Desulfobacterota_F;c_Desulfuromonadia;o_Desulfuromonadales;f_BM103;g_VAUL01;s_VAUL01 sp005774545 | 73.02 | 0.74 |
| Deltaproteobacteria bacterium DOLJORAL78_54_23 | GCA_002747175.1 | 1.85 | 61 | 53.53 | p_Desulfobacterota;c_Desulfobacteria;o_Desulfobacterales;f_Desulfosarcinaceae;g_Desulfosarcina;s_Desulfosarcina sp002747175 | 54.03 | 0.00 |
| Deltaproteobacteria bacterium DOLZORAL124_49_6 | GCA_002747805.1 | 1.55 | 89 | 48.77 | p_Desulfobacterota;c_Desulfobulbia;o_Desulfobulbales;f_Desulfocapsaceae;g_GCA-2747805;s_GCA-2747805 sp002747805 | 89.16 | 4.96 |
| Deltaproteobacteria bacterium ER2bin7 | GCA_002753725.1 | 5.97 | 653 | 38.47 | p_Desulfobacterota;c_Desulfobacteria;o_Desulfobacterales;f_Magnetomoraceae;g_Magnetomorum;s_Magnetomorum sp002753725 | 86.90 | 1.29 |
| Deltaproteobacteria bacterium GWA2_55_10 | GCA_001797415.1 | 2.24 | 161 | 55.63 | p_Desulfobacterota_F;c_GWC2-55-46;o_GWC2-55-46;f_GWC2-55-46;g_GWC2-55-46_A;s_GWC2-55-46_A sp001797415 | 96.09 | 0.65 |
| Deltaproteobacteria bacterium GWC2_55_46 | GCA_001595385.3 | 2.79 | 3 | 55.4 | p_Desulfobacterota_F;c_GWC2-55-46;o_GWC2-55-46;f_GWC2-55-46;g_GWC2-55-46;s_GWC2-55-46 sp001595385 | 68.08 | 1.45 |
| Deltaproteobacteria bacterium GWF2_42_12 | GCA_001797665.1 | 2.44 | 126 | 42.05 | p_Desulfobacterota_F;c_GWC2-55-46;o_GWC2-55-46;f_UBA9637;g_UBA9637;s_UBA9637 sp001797665 | 96.77 | 1.29 |
| Deltaproteobacteria bacterium HGW-Deltaproteobacteria-1 | GCA_002840635.1 | 3.30 | 558 | 47.91 | p_Desulfobacterota;c_Syntrophia;o_Syntrophales;f_Smithellaceae;g_UBA8904;s_UBA8904 sp002840635 | 92.22 | 2.78 |
| Deltaproteobacteria bacterium HGW-Deltaproteobacteria-12 | GCA_002840565.1 | 3.99 | 64 | 47.28 | p_Desulfobacterota;c_Syntrophia;o_Syntrophales;f_Smithellaceae;g_Fen-1166;s_Fen-1166 sp002840565 | 90.39 | 0.65 |
| Deltaproteobacteria bacterium HGW-Deltaproteobacteria-13 | GCA_002840555.1 | 3.51 | 29 | 45.65 | p_Desulfobacterota;c_Syntrophia;o_Syntrophales;f_Smithellaceae;g_Smithella;s_Smithella sp002840555 | 98.35 | 1.29 |
| Deltaproteobacteria bacterium HGW-Deltaproteobacteria-15 | GCA_002840535.1 | 6.98 | 106 | 54.47 | p_Desulfobacterota;c_Desulfobacteria;o_Desulfatiglandales;f_HGW-15;g_HGW-15;s_HGW-15 sp002840535 | 85.04 | 5.94 |
| Deltaproteobacteria bacterium HGW-Deltaproteobacteria-19 | GCA_002841865.1 | 3.98 | 87 | 59.86 | p_Desulfobacterota;c_Syntrophia;o_Syntrophales;f_PHBD01;g_PHBD01;s_PHBD01 sp002841865 | 75.18 | 0.65 |
| Deltaproteobacteria bacterium HGW-Deltaproteobacteria-2 | GCA_002840505.1 | 3.28 | 11 | 42.84 | p_Desulfobacterota;c_Syntrophia;o_Syntrophales;f_Smithellaceae;g_Smithella;s_Smithella sp002840505 | 83.50 | 3.87 |
| Deltaproteobacteria bacterium HGW-Deltaproteobacteria-3 | GCA_002841785.1 | 2.85 | 388 | 57.95 | p_Desulfobacterota;c_Desulfobulbia;o_Desulfobulbales;f_Desulfurivibrionaceae;g_UBA2262;s_UBA2262 sp002841785 | 86.27 | 2.10 |
| Deltaproteobacteria bacterium HGW-Deltaproteobacteria-4 | GCA_002841765.1 | 3.02 | 30 | 55.77 | p_Desulfobacterota_F;c_Desulfuromonadia;o_Desulfuromonadales;f_Trichloromonadaceae;g_UBA2197;s_UBA2197 sp002841765 | 96.76 | 3.92 |
| Deltaproteobacteria bacterium HGW-Deltaproteobacteria-6 | GCA_002840435.1 | 4.09 | 53 | 49.46 | p_Desulfobacterota;c_Syntrophia;o_Syntrophales;f_Smithellaceae;g_UBA8904;s_UBA8904 sp002840435 | 96.02 | 0.65 |
| Deltaproteobacteria bacterium M0040 | GCA_006226895.1 | 1.63 | 270 | 53 | p_Desulfobacterota_F;c_Desulfuromonadia;o_Desulfuromonadales;f_BM103;g_M0040;s_M0040 sp006226895 | 98.71 | 3.23 |
| Deltaproteobacteria bacterium MAG 48 | GCA_003973265.1 | 3.27 | 113 | 52.35 | p_Desulfobacterota;c_Desulfobacteria;o_Desulfobacterales;f_QNYZ01;g_QNYZ01;s_QNYZ01 sp003973265 | 87.24 | 3.49 |
| Deltaproteobacteria bacterium MAG_00134_naph_006 | GCA_013349875.1 | 1.50 | 692 | 49.54 | p_Desulfobacterota;c_Desulfobacteria;o_Desulfatiglandales;f_Desulfatiglandaceae;g_DUZB01;s_DUZB01 sp013349725 | 94.84 | 1.94 |
| Deltaproteobacteria bacterium MAG_00241_naph_010 | GCA_013349745.1 | 1.55 | 324 | 49.45 | p_Desulfobacterota;c_Desulfobacteria;o_Desulfatiglandales;f_Desulfatiglandaceae;g_DUZB01;s_DUZB01 sp013349725 | 86.38 | 1.29 |
| Deltaproteobacteria bacterium MAG_00792_naph_016 | GCA_013349725.1 | 3.03 | 409 | 49.74 | p_Desulfobacterota;c_Desulfobacteria;o_Desulfatiglandales;f_Desulfatiglandaceae;g_DUZB01;s_DUZB01 sp013349725 | 91.00 | 4.32 |
| Deltaproteobacteria bacterium MAG_09788_naph_37 | GCA_013349635.1 | 0.90 | 137 | 47.23 | p_Desulfobacterota;c_Desulfobacteria;o_Desulfatiglandales;f_Desulfatiglandaceae;g_JACNIU01;s_ | 91.46 | 4.78 |
| Deltaproteobacteria bacterium MAG_15370_dsfb_81 | GCA_013349495.1 | 3.87 | 334 | 48.4 | p_Desulfobacterota;c_Desulfobacteria;o_Desulfobacterales;f_SURF-15;g_DUZK01;s_DUZK01 sp013349495 | 74.22 | 3.23 |
| Deltaproteobacteria bacterium MAG_17929_sntb_26 | GCA_013349515.1 | 2.78 | 276 | 53.1 | p_Desulfobacterota;c_Syntrophobacteria;o_Syntrophobacterales;f_Syntrophobacteraceae;g_SbD1;s_ | 90.45 | 5.43 |
| Deltaproteobacteria bacterium MAG_17996_sntb_20 | GCA_013349395.1 | 1.69 | 454 | 53.11 | p_Desulfobacterota;c_Syntrophobacteria;o_Syntrophobacterales;f_Syntrophobacteraceae;g_SbD1;s_ | 68.18 | 2.76 |
| Deltaproteobacteria bacterium MAG_22204_dsfv_001 | GCA_013349285.1 | 2.68 | 75 | 52.74 | p_Desulfobacterota;c_Desulfobulbia;o_Desulfobulbales;f_Desulfurivibrionaceae;g_DUZU01;s_DUZU01 sp013349285 | 81.84 | 0.00 |
| Deltaproteobacteria bacterium MAG_22309_dsfv_022 | GCA_013349295.1 | 2.90 | 66 | 55.15 | p_Desulfobacterota;c_Desulfobulbia;o_Desulfobulbales;f_Desulfurivibrionaceae;g_UBA2262;s_UBA2262 sp013349295 | 95.00 | 3.86 |
| Deltaproteobacteria bacterium nDJH13bin5 | GCA_015233625.1 | 4.59 | 619 | 51.85 | p_Desulfobacterota;c_Desulfobaccia_A;o_;f_;g_;s_ | 90.97 | 5.16 |
| Deltaproteobacteria bacterium nDJH2bin9 | GCA_015233355.1 | 4.76 | 633 | 51.72 | p_Desulfobacterota;c_Desulfobaccia_A;o_;f_;g_;s_ | 99.58 | 0.84 |

| Genome name | NCBI/IMG accession | Size, Mbp | Scaff, no. | GC, % | GTDB Taxonomy | Completeness, % | Contamination, % |
|---|---|---|---|---|---|---|---|
| Deltaproteobacteria bacterium nDJH5bin8 | GCA_015233285.1 | 3.75 | 521 | 51.53 | p_Desulfobacterota;c_Desulfobacteria;o_;f_;g_;s_ | 98.39 | 2.58 |
| Deltaproteobacteria bacterium nDJH6bin12 | GCA_015233295.1 | 3.95 | 569 | 51.41 | p_Desulfobacterota;c_Desulfobaccia_A;o_;f_;g_;s_ | 99.35 | 1.61 |
| Deltaproteobacteria bacterium nDJH6bin20 | GCA_015233135.1 | 2.61 | 64 | 45.14 | p_UBA10199;c_UBA10199;o_DSSB01;f_;g_;s_ | 95.85 | 3.18 |
| Deltaproteobacteria bacterium nDJH8bin5 | GCA_015233055.1 | 4.05 | 527 | 51.56 | p_Desulfobacterota;c_Desulfobaccia_A;o_;f_;g_;s_ | 93.19 | 0.39 |
| Deltaproteobacteria bacterium nXXbin1 | GCA_015228875.1 | 4.58 | 228 | 47.04 | p_UBA10199;c_UBA10199;o_;f_;g_;s_ | 99.70 | 0.89 |
| Deltaproteobacteria bacterium Phox-21 | GCA_001896555.1 | 3.31 | 63 | 57.7 | p_Desulfobacterota;c_Desulfomonilia;o_UBA1062;f_UBA1062;g_UBA1062;s_UBA1062 sp001896555 | 83.26 | 2.47 |
| Deltaproteobacteria bacterium PowLak16_MAG11 | GCA_007280345.1 | 2.43 | 223 | 50.5 | p_Desulfobacterota;c_QYQD01;o_QYQD01;f_QYQD01;g_QYQD01;s_QYQD01 sp007280345 | 77.26 | 1.68 |
| Deltaproteobacteria bacterium PowLak16_MAG18 | GCA_007280195.1 | 1.67 | 140 | 65.04 | p_Desulfobacterota_E;c_MBNT15;o_MBNT15;f_MBNT15;g_CG2-30-66-27;s_CG2-30-66-27 sp007280195 | 81.61 | 1.45 |
| Deltaproteobacteria bacterium RBG_13_43_22 | GCA_001797675.1 | 3.23 | 39 | 48.06 | p_Desulfobacterota;c_Desulfobaccia_A;o_RBG-13-43-22;f_RBG-13-43-22;g_RBG-13-43-22;s_RBG-13-43-22 sp001797675 | 58.23 | 1.35 |
| Deltaproteobacteria bacterium RBG_13_49_15 | GCA_001797695.1 | 2.59 | 201 | 49.16 | p_Desulfobacterota;c_Desulfobacteria;o_Desulfobacterales;f_UBA2156;g_RBG-13-49-15;s_RBG-13-49-15 sp001797695 | 94.44 | 0.85 |
| Deltaproteobacteria bacterium RBG_13_52_11 | GCA_001797745.1 | 2.54 | 107 | 52.05 | p_Desulfobacterota;c_BSN033;o_B13-G15;f_RBG-16-54-18;g_RBG-13-52-11;s_RBG-13-52-11 sp001797745 | 100.00 | 0.00 |
| Deltaproteobacteria bacterium RBG_16_44_11 | GCA_001797845.1 | 2.43 | 47 | 43.54 | p_Desulfobacterota;c_Syntrophia;o_Syntrophales;f_Smithellaceae;g_UBA4810;s_UBA4810 sp001797845 | 97.74 | 1.64 |
| Deltaproteobacteria bacterium RBG_16_54_11 | GCA_001797905.1 | 2.37 | 139 | 53.8 | p_Desulfobacterota;c_BSN033;o_B13-G15;f_RBG-16-54-18;g_RBG-16-54-11;s_RBG-16-54-11 sp001797905 | 92.95 | 6.42 |
| Deltaproteobacteria bacterium RIFCSPHIGHO2_02_FULL_43_33 | GCA_001798165.1 | 2.19 | 126 | 42.54 | p_Desulfobacterota_F;c_GWC2-55-46;o_GWC2-55-46;f_UBA9637;g_UBA10170;s_UBA10170 sp001798165 | 91.10 | 3.38 |
| Deltaproteobacteria bacterium RIFOXYD12_FULL_50_9 | GCA_001799225.1 | 3.77 | 274 | 50.16 | p_Desulfobacterota;c_Desulfobulbia;o_Desulfobulbales;f_Desulfurivibrionaceae;g_XYD12-FULL-50-9;s_XYD12-FULL-50-9 sp001799225 | 76.34 | 1.13 |
| Deltaproteobacteria bacterium RIFOXYD12_FULL_55_16 | GCA_001799255.1 | 2.46 | 90 | 55.67 | p_Desulfobacterota;c_Desulfobulbia;o_Desulfobulbales;f_Desulfurivibrionaceae;g_UBA2262;s_UBA2262 sp001799255 | 95.45 | 0.60 |
| Deltaproteobacteria bacterium RIFOXYD12_FULL_56_24 | GCA_001799285.1 | 3.04 | 67 | 56.42 | p_Desulfobacterota;c_Desulfobulbia;o_Desulfobulbales;f_Desulfurivibrionaceae;g_UBA2262;s_UBA2262 sp001799285 | 74.86 | 1.66 |
| Deltaproteobacteria bacterium RIFOXYD12_FULL_57_12 | GCA_001799275.1 | 3.84 | 126 | 56.76 | p_Desulfobacterota;c_Desulfobulbia;o_Desulfobulbales;f_BM004;g_YD12-FULL-57-12;s_YD12-FULL-57-12 sp001799275 | 86.90 | 2.10 |
| Deltaproteobacteria bacterium SM23_61 | GCA_001304105.1 | 4.24 | 286 | 56.18 | p_Desulfobacterota;c_SM23-61;o_SM23-61;f_SM23-61;g_SM23-61;s_SM23-61 sp001304105 | 98.32 | 1.68 |
| Deltaproteobacteria bacterium SURF_34 | GCA_003599365.1 | 1.92 | 108 | 43.36 | p_Desulfobacterota;c_SM23-61;o_UBA8473;f_UBA8473;g_UBA8473;s_UBA8473 sp003599365 | 94.12 | 2.10 |
| Deltaproteobacteria bacterium SURF_52 | GCA_003599195.1 | 3.92 | 179 | 57.38 | p_Desulfobacterota;c_Desulfobaccia;o_Desulfobaccales;f_0-14-0-80-60-11;g_SURF-52;s_SURF-52 sp003599195 | 98.88 | 0.00 |
| Deltaproteobacteria bacterium tcs-42 | GCA_002049545.1 | 2.55 | 121 | 48.34 | p_Desulfobacterota_F;c_Desulfuromonadia;o_Desulfuromonadales;f_Geopsychrobacteraceae;g_Desulfuromusa;s_Desulfuromusa sp002049545 | 91.94 | 7.31 |
| Deltaproteobacteria bacterium UBA11853 | GCA_003513855.1 | 2.01 | 259 | 64.22 | p_Desulfobacterota_E;c_MBNT15;o_MBNT15;f_MBNT15;g_CSP1-8;s_CSP1-8 sp003513855 | 97.42 | 5.16 |
| Deltaproteobacteria bacterium UBA1386 | GCA_002305765.1 | 3.57 | 121 | 57.67 | p_Desulfobacterota_G;c_Syntrophorhabdia;o_Syntrophorhabdales;f_Syntrophorhabdaceae;g_Delta-02;s_Delta-02 sp002305765 | 76.60 | 1.68 |
| Deltaproteobacteria bacterium YD0425bin50 | GCA_002753105.1 | 4.97 | 589 | 36.91 | p_Desulfobacterota;c_Desulfobacteria;o_Desulfobacterales;f_YD0425bin50;g_YD0425bin50;s_YD0425bin50 sp002753105 | 96.94 | 1.45 |
| Deltaproteobacteria bacterium YD0425bin51 | GCA_002753225.1 | 5.22 | 134 | 32.23 | p_Desulfobacterota;c_Desulfobacteria;o_Desulfobacterales;f_YD0425bin51;g_YD0425bin51;s_YD0425bin51 sp002753225 | 96.77 | 1.02 |
| Desulfacinum hydrothermale DSM 13146 | GCF_900176285.1 | 3.70 | 59 | 60.97 | p_Desulfobacterota;c_Syntrophobacteria;o_Syntrophobacterales;f_DSM-9756;g_Desulfacinum;s_Desulfacinum hydrothermale | 89.58 | 1.69 |
| Desulfacinum infernum DSM 9756 | GCF_900129305.1 | 4.24 | 81 | 61.97 | p_Desulfobacterota;c_Syntrophobacteria;o_Syntrophobacterales;f_DSM-9756;g_Desulfacinum;s_Desulfacinum infernum | 70.52 | 0.67 |
| Desulfamplus magnetovallimortis PRJEB14757 | GCF_900170035.1 | 6.68 | 108 | 40.72 | p_Desulfobacterota;c_Desulfobacteria;o_Desulfobacterales;f_Desulfobacteraceae;g_Desulfamplus;s_Desulfamplus magnetovallimortis | 92.20 | 4.77 |
| Desulfamplus sp nDH2bin3 | GCA_015233755.1 | 2.54 | 407 | 35.65 | p_Desulfobacterota;c_Desulfobacteria;o_Desulfobacterales;f_Desulfobacteraceae;g_Desulfamplus;s_ | 95.45 | 0.91 |
| Desulfamplus sp nER1bin2 | GCA_015232935.1 | 2.88 | 486 | 42.66 | p_Desulfobacterota;c_Desulfobacteria;o_Desulfobacterales;f_Desulfobacteraceae;g_Desulfamplus;s_ | 93.18 | 1.42 |
| Desulfamplus sp nHGRbin17 | GCA_015232555.1 | 3.82 | 106 | 37.82 | p_Desulfobacterota;c_Desulfobacteria;o_Desulfobacterales;f_Desulfobacteraceae;g_Desulfamplus;s_ | 83.99 | 1.08 |
| Desulfamplus sp nHLHbin7 | GCA_015232465.1 | 3.02 | 147 | 37.75 | p_Desulfobacterota;c_Desulfobacteria;o_Desulfobacterales;f_Desulfobacteraceae;g_Desulfamplus;s_ | 80.42 | 0.00 |
| Desulfamplus sp nJC1bin9 | GCA_015232455.1 | 4.35 | 82 | 35.99 | p_Desulfobacterota;c_Desulfobacteria;o_Desulfobacterales;f_Desulfobacteraceae;g_Desulfamplus;s_ | 67.05 | 0.91 |
| Desulfamplus sp nN2-2bin5 | GCA_015232035.1 | 3.18 | 215 | 37.93 | p_Desulfobacterota;c_Desulfobacteria;o_Desulfobacterales;f_Desulfobacteraceae;g_Desulfamplus;s_ | 96.36 | 0.00 |
| Desulfamplus sp nS315bin3 | GCA_015231705.1 | 3.60 | 282 | 40.67 | p_Desulfobacterota;c_Desulfobacteria;o_Desulfobacterales;f_Desulfobacteraceae;g_Desulfamplus;s_ | 96.77 | 0.32 |
| Desulfamplus sp nS315bin44 | GCA_015231615.1 | 3.86 | 200 | 36.17 | p_Desulfobacterota;c_Desulfobacteria;o_Desulfobacterales;f_Desulfobacteraceae;g_Desulfamplus;s_ | 83.90 | 1.69 |
| Desulfamplus sp nS315bin9 | GCA_015231655.1 | 3.17 | 195 | 37.59 | p_Desulfobacterota;c_Desulfobacteria;o_Desulfobacterales;f_Desulfobacteraceae;g_Desulfamplus;s_ | 94.12 | 0.81 |
| Desulfamplus sp nTSbin15 | GCA_015231525.1 | 5.49 | 851 | 40.46 | p_Desulfobacterota;c_Desulfobacteria;o_Desulfobacterales;f_Desulfobacteraceae;g_Desulfamplus;s_ | 91.91 | 0.89 |
| Desulfamplus sp nTSbin20 | GCA_015231465.1 | 3.63 | 402 | 47.21 | p_Desulfobacterota;c_Desulfobacteria;o_Desulfobacterales;f_Desulfobacteraceae;g_Desulfamplus;s_ | 94.28 | 0.00 |
| Desulfamplus sp. nTSbin4 | GCA_015231415.1 | 3.23 | 196 | 37.6 | p_Desulfobacterota;c_Desulfobacteria;o_Desulfobacterales;f_Desulfobacteraceae;g_Desulfamplus;s_ | 93.21 | 1.19 |
| Desulfamplus sp. nXXbin12 | GCA_015228835.1 | 2.59 | 369 | 37.76 | p_Desulfobacterota;c_Desulfobacteria;o_Desulfobacterales;f_Desulfobacteraceae;g_Desulfamplus;s_ | 84.99 | 1.59 |
| Desulfamplus_sp nTSbin21 | GCA_015231395.1 | 2.81 | 548 | 39.13 | p_Desulfobacterota;c_Desulfobacteria;o_Desulfobacterales;f_Desulfobacteraceae;g_Desulfamplus;s_ | 90.33 | 0.60 |
| Desulfarculaceae bacterium JdFR-95 | GCA_002011835.1 | 2.81 | 499 | 68.22 | p_Desulfobacterota;c_Desulfarculia;o_Desulfarculales;f_Desulfarculaceae;g_JdFR-95;s_JdFR-95 sp002011835 | 95.45 | 0.00 |
| Desulfarculales bacterium UBA696 | GCA_002298995.1 | 4.17 | 77 | 53.8 | p_Desulfobacterota;c_Syntrophobacteria;o_Syntrophobacterales;f_Syntrophobacteraceae;g_UBA696;s_UBA696 sp002298995 | 82.83 | 1.79 |
| Desulfarculales bacterium UBA702 | GCA_002298165.1 | 4.63 | 119 | 52.03 | p_Desulfobacterota;c_Syntrophobacteria;o_Syntrophobacterales;f_Syntrophobacteraceae;g_UBA696;s_UBA696 sp002298165 | 61.13 | 1.44 |
| Desulfarculus baarsii DSM 2075 | GCF_000143965.1 | 3.66 | 1 | 65.7 | p_Desulfobacterota;c_Desulfarculia;o_Desulfarculales;f_Desulfarculaceae;g_Desulfarculus;s_Desulfarculus baarsii | 100.00 | 1.77 |
| Desulfarculus sp. SURF_10 | GCA_003605055.1 | 3.60 | 78 | 66.98 | p_Desulfobacterota;c_Desulfarculia;o_Desulfarculales;f_Desulfarculaceae;g_SURF-10;s_SURF-10 sp003605055 | 100.00 | 0.00 |
| Desulfatibacillum aliphaticivorans DSM 15576 | GCF_000429905.1 | 6.47 | 64 | 54.43 | p_Desulfobacterota;c_Desulfobacteria;o_Desulfobacterales;f_Desulfatibacillaceae;g_Desulfatibacillum;s_Desulfatibacillum aliphaticivorans | 76.75 | 0.65 |
| Desulfatibacillum alkenivorans DSM 16219 | GCF_900142135.1 | 6.47 | 102 | 54.95 | p_Desulfobacterota;c_Desulfobacteria;o_Desulfobacterales;f_Desulfatibacillaceae;g_Desulfatibacillum;s_Desulfatibacillum alkenivorans | 77.77 | 1.45 |
| Desulfatiglans anilini DSM 4660 | GCF_000422285.1 | 4.67 | 107 | 58.78 | p_Desulfobacterota;c_Desulfobacteria;o_Desulfatiglandales;f_Desulfatiglandaceae;g_Desulfatiglans;s_Desulfatiglans anilini | 98.18 | 0.91 |
| Desulfatirhabdium butyrativorans DSM 18734 | GCF_000429925.1 | 4.48 | 73 | 54.92 | p_Desulfobacterota;c_Desulfobacteria;o_Desulfobacterales;f_Desulfatirhabdiaceae;g_Desulfatirhabdium;s_Desulfatirhabdium butyrativorans | 80.00 | 5.34 |
| Desulfatitalea sp. BRH_c12 | GCA_000961655.1 | 6.00 | 91 | 55.05 | p_Desulfobacterota;c_Desulfobacteria;o_Desulfobacterales;f_Desulfosarcinaceae;g_Desulfatitalea;s_Desulfatitalea sp000961655 | 98.18 | 0.83 |
| Desulfatitalea tepidiphila S28bF | GCF_001293685.1 | 5.61 | 7 | 56.68 | p_Desulfobacterota;c_Desulfobacteria;o_Desulfobacterales;f_Desulfosarcinaceae;g_Desulfatitalea;s_Desulfatitalea tepidiphila | 88.71 | 3.23 |
| Desulfobacca acetoxidans DSM 11109 | GCF_000195295.1 | 3.28 | 1 | 52.89 | p_Desulfobacterota;c_Desulfobaccia;o_Desulfobaccales;f_Desulfobaccaceae;g_Desulfobacca;s_Desulfobacca acetoxidans | 66.01 | 1.08 |
| Desulfobacca sp. 4484_104 | GCA_002049795.1 | 2.65 | 181 | 53.64 | p_Desulfobacterota;c_Desulfobaccia;o_Desulfobaccales;f_Desulfobaccaceae;g_Desulfobacca_B;s_Desulfobacca_B sp002049795 | 88.32 | 2.25 |
| Desulfobacter curvatus DSM 3379 | GCF_000373985.1 | 5.64 | 234 | 46.9 | p_Desulfobacterota;c_Desulfobacteria;o_Desulfobacterales;f_Desulfobacteraceae;g_Desulfobacter;s_Desulfobacter curvatus | 93.58 | 2.78 |
| Desulfobacter hydrogenophilus AcRS1 | GCF_004319545.1 | 5.16 | 3 | 46.53 | p_Desulfobacterota;c_Desulfobacteria;o_Desulfobacterales;f_Desulfobacteraceae;g_Desulfobacter;s_Desulfobacter hydrogenophilus | 56.31 | 0.00 |

| Genome name | NCBI/IMG accession | Size, Mbp | Scaff, no. | GC, % | GTDB Taxonomy | Completeness, % | Contamination, % |
|---|---|---|---|---|---|---|---|
| Desulfobacter postgatei 2ac9 | GCF_000233695.2 | 3.97 | 1 | 47.2 | p_Desulfobacterota;c_Desulfobacteria;o_Desulfobacterales;f_Desulfobacteraceae;g_Desulfobacter;s_Desulfobacter postgatei | 54.15 | 0.65 |
| Desulfobacter postgatei DOLJORAL78_47_202 | GCA_002747145.1 | 1.48 | 65 | 47 | p_Desulfobacterota;c_Desulfobacteria;o_Desulfobacterales;f_Desulfobacteraceae;g_Desulfobacter;s_Desulfobacter postgatei_A | 71.94 | 0.09 |
| Desulfobacter vibrioformis DSM 8776 | GCF_000745975.1 | 4.47 | 86 | 48.64 | p_Desulfobacterota;c_Desulfobacteria;o_Desulfobacterales;f_Desulfobacteraceae;g_Desulfobacter;s_Desulfobacter vibrioformis | 89.77 | 0.00 |
| Desulfobacteraceae bacterium 4484_190.2 | GCA_002050025.1 | 2.37 | 232 | 45.82 | p_Desulfobacterota;c_Desulfobacteria;o_Desulfatiglandales;f_Desulfatiglandaceae;g_4484-190-2;s_4484-190-2 sp002050025 | 62.78 | 3.98 |
| Desulfobacteraceae bacterium 4572_123 | GCA_002084545.1 | 3.18 | 168 | 47.51 | p_Desulfobacterota;c_Desulfobacteria;o_Desulfobacterales;f_4572-123;g_4572-123;s_4572-123 sp002084545 | 90.99 | 1.68 |
| Desulfobacteraceae bacterium 4572_130 | GCA_002084425.1 | 1.94 | 24 | 29.47 | p_Desulfobacterota;c_Desulfobacteria;o_Desulfobacterales;f_Desulfobacteraceae;g_4572-130;s_4572-130 sp002084425 | 88.04 | 0.65 |
| Desulfobacteraceae bacterium 4572_35.2 | GCA_002084665.1 | 1.90 | 129 | 46.14 | p_Desulfobacterota_F;c_Desulfuromonadia;o_Desulfuromonadales;f_Desulfuromonadaceae;g_Desulfuromonas;s_Desulfuromonas sp002084665 | 61.51 | 1.66 |
| Desulfobacteraceae bacterium 4572_89 | GCA_002085465.1 | 1.70 | 98 | 43.64 | p_Desulfobacterota;c_Desulfobacteria;o_Desulfobacterales;f_Desulfobacteraceae;g_NBML01;s_NBML01 sp002085465 | 98.49 | 1.94 |
| Desulfobacteraceae bacterium B1Sed10_16 | GCA_003551985.1 | 2.12 | 442 | 54.15 | p_Desulfobacterota;c_Desulfobacteria;o_Desulfobacterales;f_SURF-3;g_B1SED10-16;s_B1SED10-16 sp003551985 | 95.85 | 3.69 |
| Desulfobacteraceae bacterium BM002 | GCA_002899795.1 | 3.80 | 638 | 50.24 | p_Desulfobacterota;c_Syntrophobacteria;o_BM002;f_BM002;g_BM002;s_BM002 sp002899795 | 93.64 | 6.39 |
| Desulfobacteraceae bacterium BM005 | GCA_002868985.1 | 3.05 | 577 | 43.45 | p_Desulfobacterota;c_Desulfobacteria;o_Desulfobacterales;f_UBA11574;g_UBA11574;s_UBA11574 sp002868985 | 54.79 | 0.00 |
| Desulfobacteraceae bacterium CG2_30_51_40 | GCA_001874005.1 | 3.44 | 229 | 50.72 | p_Desulfobacterota;c_Desulfobacteria;o_Desulfatiglandales;f_Desulfatiglandaceae;g_CG2-30-51-40;s_CG2-30-51-40 sp001874005 | 97.10 | 0.65 |
| Desulfobacteraceae bacterium CSSed165cm_505 | GCA_007131845.1 | 2.05 | 410 | 55.22 | p_Desulfobacterota;c_Desulfobacteria;o_Desulfobacterales;f_SURF-3;g_B1SED10-16;s_B1SED10-16 sp007131845 | 95.85 | 3.69 |
| Desulfobacteraceae bacterium Eth-SRB2 | GCA_004193595.1 | 5.26 | 83 | 43.72 | p_Desulfobacterota;c_Desulfobacteria;o_Desulfobacterales;f_UBA11574;g_S5133MH16;s_S5133MH16 sp004193595 | 81.21 | 0.32 |
| Desulfobacteraceae bacterium maxbin2.1429 | GCA_003819975.1 | 5.14 | 978 | 48.48 | p_Desulfobacterota;c_Desulfobacteria;o_Desulfatiglandales;f_HGW-15;g_RPPU01;s_RPPU01 sp003819975 | 92.80 | 1.68 |
| Desulfobacteraceae bacterium RAAP-1 | GCA_001443525.1 | 4.09 | 193 | 49.94 | p_Desulfobacterota;c_Desulfobacteria;o_Desulfobacterales;f_Desulfatirhabdiaceae;g_RAAP-1;s_RAAP-1 sp001443525 | 93.87 | 1.33 |
| Desulfobacteraceae bacterium SURF_15 | GCA_003599475.1 | 5.29 | 268 | 54.29 | p_Desulfobacterota;c_Desulfobacteria;o_Desulfobacterales;f_SURF-15;g_SURF-15;s_SURF-15 sp003599475 | 95.48 | 1.61 |
| Desulfobacteraceae bacterium SURF_3 | GCA_003599885.1 | 4.51 | 90 | 49.04 | p_Desulfobacterota;c_Desulfobacteria;o_Desulfobacterales;f_SURF-3;g_SURF-3;s_SURF-3 sp003599885 | 92.06 | 4.60 |
| Desulfobacteraceae bacterium SURF_33 | GCA_003597945.1 | 3.72 | 137 | 52.78 | p_Desulfobacterota;c_Desulfobacteria;o_Desulfobacterales;f_SURF-3;g_SURF-33;s_SURF-33 sp003597945 | 80.65 | 0.00 |
| Desulfobacteraceae bacterium SURF_4 | GCA_003599575.1 | 6.45 | 197 | 55.75 | p_Desulfobacterota;c_Desulfobacteria;o_Desulfobacterales;f_Desulfosarcinaceae;g_Desulfatitalea;s_Desulfatitalea sp003599575 | 68.49 | 0.86 |
| Desulfobacteraceae bacterium SURF_67 | GCA_003599015.1 | 3.00 | 149 | 49.3 | p_Desulfobacterota;c_Desulfobacteria;o_Desulfatiglandales;f_Desulfatiglandaceae;g_CG2-30-51-40;s_CG2-30-51-40 sp003599015 | 98.81 | 0.62 |
| Desulfobacteraceae bacterium UBA11574 | GCA_003486165.1 | 2.82 | 505 | 43.35 | p_Desulfobacterota;c_Desulfobacteria;o_Desulfobacterales;f_UBA11574;g_UBA11574;s_UBA11574 sp003486165 | 98.74 | 0.84 |
| Desulfobacteraceae bacterium UBA2156 | GCA_002328165.1 | 2.94 | 78 | 45.87 | p_Desulfobacterota;c_Desulfobacteria;o_Desulfobacterales;f_UBA2156;g_UBA2156;s_UBA2156 sp002328165 | 95.80 | 3.64 |
| Desulfobacteraceae bacterium UBA2771 | GCA_002352605.1 | 3.03 | 196 | 54.14 | p_Desulfobacterota;c_Desulfobacteria;o_Desulfobacterales;f_Desulfosarcinaceae;g_Desulfosarcina;s_Desulfosarcina sp002352605 | 93.88 | 7.28 |
| Desulfobacteraceae bacterium UBA4064 | GCA_002382065.1 | 4.77 | 300 | 50.19 | p_Desulfobacterota;c_Desulfobacteria;o_Desulfobacterales;f_Desulfatirhabdiaceae;g_UBA4064;s_UBA4064 sp002382065 | 82.58 | 4.52 |
| Desulfobacteraceae bacterium UBA5616 | GCA_002423615.1 | 3.66 | 121 | 51.61 | p_Desulfobacterota;c_Desulfobacteria;o_Desulfobacterales;f_UBA5616;g_UBA5616;s_UBA5616 sp002423615 | 65.27 | 0.00 |
| Desulfobacteraceae bacterium UBA5623 | GCA_002424495.1 | 4.02 | 456 | 47.46 | p_Desulfobacterota;c_Desulfobacteria;o_Desulfatiglandales;f_Desulfatiglandaceae;g_UBA5623;s_UBA5623 sp002424495 | 96.77 | 1.29 |
| Desulfobacteraceae bacterium UBA8212 | GCA_003538835.1 | 5.56 | 584 | 53.41 | p_Desulfobacterota;c_Desulfobacteria;o_Desulfobacterales;f_Desulfobacteraceae;g_Desulfobacter;s_Desulfobacter sp003538835 | 88.71 | 0.20 |
| Desulfobacterales bacterium CG2_30_60_27 | GCA_001873115.1 | 2.52 | 157 | 60.25 | p_Desulfobacterota;c_Desulfobulbia;o_Desulfobulbales;f_Desulfurivibrionaceae;g_CG2-30-60-27;s_CG2-30-60-27 sp001873115 | 97.42 | 0.32 |
| Desulfobacterales bacterium CG23 | GCA_002771315.1 | 2.75 | 309 | 52.32 | p_Desulfobacterota;c_Desulfobacteria;o_Desulfobacterales;f_UBA2156;g_GCA-002779465;s_GCA-002779465 sp002771315 | 94.09 | 2.73 |
| Desulfobacterales bacterium CSSed165cm_255 | GCA_007134875.1 | 2.88 | 231 | 55.67 | p_Desulfobacterota;c_Desulfobacteria;o_Desulfobacterales;f_SURF-3;g_B1SED10-16;s_B1SED10-16 sp007134875 | 91.82 | 5.73 |
| Desulfobacterales bacterium GWB2_56_26 | GCA_001799365.1 | 4.51 | 263 | 55.56 | p_Desulfobacterota;c_Desulfobulbia;o_Desulfobulbales;f_Desulfocapsaceae;g_Desulforhopalus;s_Desulforhopalus sp001799365 | 91.40 | 3.41 |
| Desulfobacterales bacterium nHLHbin5 | GCA_015232515.1 | 4.58 | 421 | 32.52 | p_Desulfobacterota;c_Desulfobacteria;o_Desulfobacterales;f_YD0425bin51;g_YD0425bin51;s_YD0425bin51 sp002753225 | 92.58 | 3.37 |
| Desulfobacterales bacterium nTSbin1 | GCA_015231595.1 | 5.13 | 210 | 32.05 | p_Desulfobacterota;c_Desulfobacteria;o_Desulfobacterales;f_YD0425bin50;g_;s_ | 81.38 | 2.66 |
| Desulfobacterales bacterium nYD0425bin5 | GCA_015228685.1 | 5.63 | 133 | 32.38 | p_Desulfobacterota;c_Desulfobacteria;o_Desulfobacterales;f_YD0425bin51;g_YD0425bin51;s_YD0425bin51 sp002753225 | 94.49 | 0.71 |
| Desulfobacterales bacterium nYD0425bin6 | GCA_015228675.1 | 5.42 | 467 | 36.82 | p_Desulfobacterota;c_Desulfobacteria;o_Desulfobacterales;f_YD0425bin50;g_YD0425bin50;s_YD0425bin50 sp002753105 | 90.90 | 0.65 |
| Desulfobacterales bacterium S5133MH16 | GCA_001751005.1 | 3.02 | 574 | 43.59 | p_Desulfobacterota;c_Desulfobacteria;o_Desulfobacterales;f_UBA11574;g_S5133MH16;s_S5133MH16 sp001751005 | 80.59 | 4.52 |
| Desulfobacterium autotrophicum HRM2 | GCF_000020365.1 | 5.66 | 2 | 48.76 | p_Desulfobacterota;c_Desulfobacteria;o_Desulfobacterales;f_Desulfobacteraceae;g_Desulfobacterium_B;s_Desulfobacterium_B autotrophicum | 84.88 | 2.58 |
| Desulfobacterium vacuolatum DSM 3385 | GCF_900176365.1 | 5.04 | 80 | 46.54 | p_Desulfobacterota;c_Desulfobacteria;o_Desulfobacterales;f_Desulfobacteraceae;g_Desulfobacterium_A;s_Desulfobacterium_A vacuolatum | 91.07 | 0.60 |
| Desulfobacula toluolica Tol2 | GCF_000307105.1 | 5.20 | 1 | 41.45 | p_Desulfobacterota;c_Desulfobacteria;o_Desulfobacterales;f_Desulfobacteraceae;g_Desulfobacula;s_Desulfobacula toluolica | 95.81 | 0.00 |
| Desulfobotulus mexicanus PAR22N | GCF_006175995.1 | 3.83 | 69 | 48.98 | p_Desulfobacterota;c_Desulfobacteria;o_Desulfobacterales;f_Desulforegulaceae;g_Desulfobotulus;s_Desulfobotulus mexicanus | 98.18 | 1.82 |
| Desulfobulbaceae bacterium BM004 | GCA_002868955.1 | 2.66 | 189 | 47.45 | p_Desulfobacterota;c_Desulfobulbia;o_Desulfobulbales;f_BM004;g_BM004;s_BM004 sp002868955 | 88.17 | 2.90 |
| Desulfobulbaceae bacterium BM506 | GCA_002868945.1 | 3.71 | 83 | 56.29 | p_Desulfobacterota;c_Desulfobulbia;o_Desulfobulbales;f_Desulfurivibrionaceae;g_BM506;s_BM506 sp002868945 | 92.58 | 2.58 |
| Desulfobulbaceae bacterium BRH_c16a | GCA_000961725.1 | 6.52 | 72 | 51.68 | p_Desulfobacterota;c_Desulfobulbia;o_Desulfobulbales;f_Desulfocapsaceae;g_Desulforhopalus;s_Desulforhopalus sp000961725 | 93.12 | 2.78 |
| Desulfobulbaceae bacterium CSSed10_361 | GCA_003557565.1 | 2.23 | 357 | 59.21 | p_Desulfobacterota;c_Desulfobulbia;o_Desulfobulbales;f_Desulfurivibrionaceae;g_Desulfurivibrio;s_Desulfurivibrio sp003557565 | 57.04 | 0.12 |
| Desulfobulbaceae bacterium DB1 | GCA_001914235.1 | 3.87 | 28 | 53.78 | p_Desulfobacterota;c_Desulfobulbia;o_Desulfobulbales;f_Desulfurivibrionaceae;g_DB1;s_DB1 sp001914235 | 90.88 | 0.59 |
| Desulfobulbaceae bacterium Del10 | GCA_004332195.1 | 1.66 | 21 | 54.26 | p_Desulfobacterota;c_Desulfobulbia;o_Desulfobulbales;f_Desulfurivibrionaceae;g_Desulfurivibrio;s_Desulfurivibrio sp004332195 | 74.06 | 1.82 |
| Desulfobulbaceae bacterium nTSbin18 | GCA_015231515.1 | 3.41 | 225 | 45.02 | p_Desulfobacterota;c_Desulfobulbia;o_Desulfobulbales;f_Desulfurivibrionaceae;g_;s_ | 84.17 | 1.94 |
| Desulfobulbaceae bacterium SZUA-575 | GCA_003249645.1 | 3.74 | 182 | 45.85 | p_Desulfobacterota;c_Desulfobulbia;o_Desulfobulbales;f_Desulfocapsaceae;g_SZUA-575;s_SZUA-575 sp003249645 | 96.77 | 5.48 |
| Desulfobulbaceae bacterium SZUA-615 | GCA_003249165.1 | 3.54 | 497 | 45.36 | p_Desulfobacterota;c_Desulfobulbia;o_Desulfobulbales;f_Desulfurivibrionaceae;g_SURF-16;s_SURF-16 sp003249165 | 84.70 | 5.65 |
| Desulfobulbaceae bacterium UBA10518 | GCA_003508005.1 | 3.01 | 95 | 57.91 | p_Desulfobacterota;c_Desulfobulbia;o_Desulfobulbales;f_Desulfobulbaceae;g_UBA10518;s_UBA10518 sp003508005 | 76.96 | 2.58 |
| Desulfobulbaceae bacterium UBA11700 | GCA_003517965.1 | 2.52 | 81 | 53.21 | p_Desulfobacterota;c_Desulfobulbia;o_Desulfobulbales;f_Desulfurivibrionaceae;g_UBA2262;s_UBA2262 sp003517965 | 90.30 | 3.64 |
| Desulfobulbaceae bacterium UBA2213 | GCA_002327225.1 | 2.46 | 214 | 51.85 | p_Desulfobacterota;c_Desulfobulbia;o_Desulfobulbales;f_Desulfocapsaceae;g_UBA2270;s_UBA2270 sp002327225 | 91.35 | 6.85 |
| Desulfobulbaceae bacterium UBA2262 | GCA_002347185.1 | 2.85 | 141 | 61.68 | p_Desulfobacterota;c_Desulfobulbia;o_Desulfobulbales;f_Desulfurivibrionaceae;g_UBA2262;s_UBA2262 sp002347185 | 80.42 | 1.29 |
| Desulfobulbaceae bacterium UBA2270 | GCA_002347745.1 | 3.95 | 171 | 48.8 | p_Desulfobacterota;c_Desulfobulbia;o_Desulfobulbales;f_Desulfocapsaceae;g_UBA2270;s_UBA2270 sp002347745 | 89.19 | 0.65 |
| Desulfobulbaceae bacterium UBA2273 | GCA_002347095.1 | 3.20 | 33 | 57.57 | p_Desulfobacterota;c_Desulfobulbia;o_Desulfobulbales;f_Desulfurivibrionaceae;g_UBA2262;s_UBA2262 sp002347095 | 77.42 | 3.23 |
| Desulfobulbaceae bacterium UBA2276 | GCA_002347085.1 | 1.79 | 68 | 51.37 | p_Desulfobacterota;c_Desulfobulbia;o_Desulfobulbales;f_Desulfocapsaceae;g_UBA2270;s_UBA2270 sp002347085 | 94.84 | 3.46 |
| Desulfobulbaceae bacterium UBA5121 | GCA_002414225.1 | 3.53 | 59 | 63.19 | p_Desulfobacterota;c_Desulfobulbia;o_Desulfobulbales;f_Desulfurivibrionaceae;g_UBA5123;s_UBA5123 sp002414225 | 53.42 | 0.00 |
| Desulfobulbaceae bacterium UBA5123 | GCA_002415465.1 | 2.28 | 240 | 61.01 | p_Desulfobacterota;c_Desulfobulbia;o_Desulfobulbales;f_Desulfurivibrionaceae;g_UBA5123;s_UBA5123 sp002415465 | 86.93 | 2.07 |

| Genome name | NCBI/IMG accession | Size, Mbp | Scaff, no. | GC, % | GTDB Taxonomy | Completeness, % | Contamination, % |
|---|---|---|---|---|---|---|---|
| Desulfobulbaceae bacterium UBA5611 | GCA_002424635.1 | 2.95 | 95 | 50.29 | p__Desulfobacterota;c__Desulfobulbia;o__Desulfobulbales;f__Desulfocapsaceae;g__UBA2270;s__UBA2270 sp002424635 | 96.63 | 0.00 |
| Desulfobulbaceae bacterium UBA5628 | GCA_002421565.1 | 1.80 | 197 | 60.1 | p__Desulfobacterota;c__Desulfobulbia;o__Desulfobulbales;f__Desulfurivibrionaceae;g__UBA5628;s__UBA5628 sp002421565 | 98.90 | 1.10 |
| Desulfobulbus mediterraneus DSM 13871 | GCF_000429965.1 | 4.80 | 68 | 57.66 | p__Desulfobacterota;c__Desulfobulbia;o__Desulfobulbales;f__Desulfobulbaceae;g__Desulfobulbus_A;s__Desulfobulbus_A mediterraneus | 85.16 | 0.65 |
| Desulfobulbus propionicus DOLZORAL124_48_34 | GCA_002746685.1 | 1.61 | 53 | 47.78 | p__Desulfobacterota;c__Desulfobulbia;o__Desulfobulbales;f__Desulfobulbaceae;g__Desulfobulbus_A;s__Desulfobulbus_A propionicus_C | 85.58 | 0.07 |
| Desulfobulbus sp. SURF_48 | GCA_003604995.1 | 2.64 | 108 | 59.65 | p__Desulfobacterota;c__Desulfobulbia;o__Desulfobulbales;f__Desulfobulbaceae;g__UBA10518;s__UBA10518 sp003604995 | 89.65 | 2.58 |
| Desulfocarbo indianensis SCBM | GCF_001184205.1 | 5.11 | 334 | 63.05 | p__Desulfobacterota;c__Desulfarculia;o__Desulfarculales;f__Desulfarculaceae;g__Desulfocarbo;s__Desulfocarbo indianensis | 98.45 | 0.00 |
| Desulfococcus multivorans DSM 2059 | GCF_001854245.1 | 4.46 | 1 | 56.83 | p__Desulfobacterota;c__Desulfobacteria;o__Desulfobacterales;f__Desulfococcaceae;g__Desulfococcus;s__Desulfococcus multivorans | 98.02 | 0.15 |
| Desulfoluna spongiiphila AA1 | GCF_900101345.1 | 6.54 | 52 | 57.24 | p__Desulfobacterota;c__Desulfobacteria;o__Desulfobacterales;f__Desulfolunaceae;g__Desulfoluna;s__Desulfoluna spongiiphila | 94.55 | 1.82 |
| Desulfonatronospira sp. CSSed162cmB_39 | GCA_007127015.1 | 1.31 | 200 | 49.04 | p__Desulfobacterota;c__Desulfovibrionia;o__Desulfovibrionales;f__Desulfonatronovibrionaceae;g__Desulfonatronospira;s__Desulfonatronospira sp007127015 | 70.18 | 0.88 |
| Desulfonatronospira sp. CSSed162cmB_565 | GCA_007125075.1 | 2.17 | 149 | 49.19 | p__Desulfobacterota;c__Desulfovibrionia;o__Desulfovibrionales;f__Desulfonatronovibrionaceae;g__Desulfonatronospira;s__Desulfonatronospira sp007125075 | 86.57 | 0.00 |
| Desulfonatronovibrio hydrogenovorans DSM 9292 | GCF_000686525.1 | 2.94 | 16 | 50.35 | p__Desulfobacterota;c__Desulfovibrionia;o__Desulfovibrionales;f__Desulfonatronovibrionaceae;g__Desulfonatronovibrio;s__Desulfonatronovibrio hydrogenovorans | 76.63 | 1.54 |
| Desulfonatronovibrio magnus AHT22 | GCF_000934755.1 | 4.81 | 134 | 44.86 | p__Desulfobacterota;c__Desulfovibrionia;o__Desulfovibrionales;f__Desulfonatronovibrionaceae;g__Desulfonatronovibrio;s__Desulfonatronovibrio magnus | 95.81 | 8.63 |
| Desulfonatronovibrio sp. CSSed10_448R1 | GCA_003558575.1 | 1.76 | 360 | 43.61 | p__Desulfobacterota;c__Desulfovibrionia;o__Desulfovibrionales;f__Desulfonatronovibrionaceae;g__Desulfonatronovibrio;s__Desulfonatronovibrio sp003558575 | 96.13 | 3.55 |
| Desulfonauticus sp. 38_4375 | GCA_001507915.1 | 2.13 | 107 | 38.21 | p__Desulfobacterota;c__Desulfovibrionia;o__Desulfovibrionales;f__Desulfonauticaceae;g__Desulfonauticus;s__Desulfonauticus sp001507915 | 82.99 | 2.90 |
| Desulfonauticus submarinus DSM 15269 | GCF_900104045.1 | 2.10 | 23 | 32.47 | p__Desulfobacterota;c__Desulfovibrionia;o__Desulfovibrionales;f__Desulfonauticaceae;g__Desulfonauticus;s__Desulfonauticus submarinus | 95.48 | 2.71 |
| Desulfonema ishimotonii Tokyo 01 | GCF_003851005.1 | 6.64 | 9 | 53.5 | p__Desulfobacterota;c__Desulfobacteria;o__Desulfobacterales;f__Desulfococcaceae;g__Desulfonema;s__Desulfonema ishimotonii | 97.73 | 4.55 |
| Desulfopila aestuarii DSM 18488 | GCF_900143695.1 | 6.07 | 103 | 49.63 | p__Desulfobacterota;c__Desulfobulbia;o__Desulfobulbales;f__Desulfocapsaceae;g__Desulfopila;s__Desulfopila aestuarii | 93.23 | 1.29 |
| Desulfopila sp. IMCC35005 | GCA_005116655.1 | 6.69 | 63 | 44.28 | p__Desulfobacterota;c__Desulfobulbia;o__Desulfobulbales;f__Desulfocapsaceae;g__Desulfopila;s__Desulfopila sp005116655 | 96.45 | 0.65 |
| Desulfopila sp. IMCC35006 | GCF_005116645.1 | 5.62 | 95 | 49.75 | p__Desulfobacterota;c__Desulfobulbia;o__Desulfobulbales;f__Desulfocapsaceae;g__Desulforhopalus;s__Desulforhopalus sp005116645 | 99.34 | 0.20 |
| Desulforegula conservatrix Mb1Pa | GCF_000426225.1 | 4.47 | 178 | 42.39 | p__Desulfobacterota;c__Desulfobacteria;o__Desulfobacterales;f__Desulforegulaceae;g__Desulforegula;s__Desulforegula conservatrix | 98.18 | 0.91 |
| Desulforhabdus sp. SDB_sulfate2 | GCA_007244385.1 | 5.20 | 119 | 54.39 | p__Desulfobacterota;c__Syntrophobacteria;o__Syntrophobacterales;f__Syntrophobacteraceae;g__Desulforhabdus;s__Desulforhabdus sp007244385 | 78.46 | 1.04 |
| Desulforhopalus sp. IMCC35007 | GCF_005116575.1 | 5.74 | 127 | 45.99 | p__Desulfobacterota;c__Desulfobulbia;o__Desulfobulbales;f__Desulfocapsaceae;g__Desulforhopalus;s__Desulforhopalus sp005116575 | 66.62 | 1.49 |
| Desulfosarcina cetonica JCM 12296 | GCF_001311845.1 | 7.09 | 558 | 55.73 | p__Desulfobacterota;c__Desulfobacteria;o__Desulfobacterales;f__Desulfosarcinaceae;g__Desulfosarcina;s__Desulfosarcina cetonica | 98.71 | 0.00 |
| Desulfosediminicola ganghwensis IMCC35004 | GCA_005116675.2 | 5.65 | 1 | 48.43 | p__Desulfobacterota;c__Desulfobulbia;o__Desulfobulbales;f__Desulfocapsaceae;g__Desulfopila;s__Desulfopila sp005116675 | 95.45 | 2.73 |
| Desulfotalea sp. NORP6 | GCA_002733995.1 | 4.12 | 52 | 44.41 | p__Desulfobacterota;c__Desulfobulbia;o__Desulfobulbales;f__Desulfocapsaceae;g__Desulforhopalus;s__Desulforhopalus sp002733995 | 92.12 | 0.00 |
| Desulfotignum balticum DSM 7044 | GCF_000421285.1 | 5.12 | 5 | 51.24 | p__Desulfobacterota;c__Desulfobacteria;o__Desulfobacterales;f__Desulfobacteraceae;g__Desulfotignum;s__Desulfotignum balticum | 99.58 | 0.84 |
| Desulfovibrio alkalitolerans DSM 16529 | GCF_000422245.1 | 3.20 | 32 | 64.48 | p__Desulfobacterota;c__Desulfovibrionia;o__Desulfovibrionales;f__Desulfovibrionaceae;g__Desulfohalovibrio;s__Desulfohalovibrio alkalitolerans | 100.00 | 4.98 |
| Desulfovibrio aminophilus DSM 12254 | GCF_000422565.1 | 3.42 | 25 | 66.25 | p__Desulfobacterota;c__Desulfovibrionia;o__Desulfovibrionales;f__Desulfovibrionaceae;g__Aminidesulfovibrio;s__Aminidesulfovibrio aminophilus | 84.52 | 1.33 |
| Desulfovibrio brasiliensis JCM 12178 | GCF_001311825.1 | 3.57 | 182 | 59.65 | p__Desulfobacterota;c__Desulfovibrionia;o__Desulfovibrionales;f__Desulfovibrionaceae;g__Pseudodesulfovibrio;s__Pseudodesulfovibrio brasiliensis | 98.18 | 0.91 |
| Desulfovibrio carbinolicus DSM 3852 | GCF_004135975.1 | 4.69 | 6 | 64.62 | p__Desulfobacterota;c__Desulfovibrionia;o__Desulfovibrionales;f__Desulfovibrionaceae;g__Solidesulfovibrio;s__Solidesulfovibrio carbinolicus | 68.40 | 2.15 |
| Desulfovibrio carbinoliphilus subsp. oakridgensis FW-101-2B | GCA_000177215.2 | 4.22 | 3 | 66.44 | p__Desulfobacterota;c__Desulfovibrionia;o__Desulfovibrionales;f__Desulfovibrionaceae;g__Solidesulfovibrio;s__Solidesulfovibrio carbinoliphilus | 74.36 | 2.52 |
| Desulfovibrio cuneatus DSM 11391 | GCF_000430005.1 | 3.36 | 76 | 53.55 | p__Desulfobacterota;c__Desulfovibrionia;o__Desulfovibrionales;f__Desulfovibrionaceae;g__Frigidesulfovibrio;s__Frigidesulfovibrio cuneatus | 59.35 | 0.65 |
| Desulfovibrio fairfieldensis CCUG 45958 | GCF_001553605.1 | 3.70 | 1 | 60.9 | p__Desulfobacterota;c__Desulfovibrionia;o__Desulfovibrionales;f__Desulfovibrionaceae;g__Desulfovibrio;s__Desulfovibrio fairfieldensis | 70.44 | 2.64 |
| Desulfovibrio gracilis DSM 16080 | GCF_900167125.1 | 3.18 | 53 | 58.43 | p__Desulfobacterota;c__Desulfovibrionia;o__Desulfovibrionales;f__Desulfovibrionaceae;g__Paucidesulfovibrio;s__Paucidesulfovibrio gracilis | 96.49 | 0.91 |
| Desulfovibrio hydrothermalis AM13 = DSM 14728 | GCF_000331025.1 | 3.71 | 2 | 45.15 | p__Desulfobacterota;c__Desulfovibrionia;o__Desulfovibrionales;f__Desulfovibrionaceae;g__Maridesulfovibrio;s__Maridesulfovibrio hydrothermalis | 95.33 | 0.84 |
| Desulfovibrio legallii H1T | GCF_004309735.1 | 2.67 | 35 | 62.92 | p__Desulfobacterota;c__Desulfovibrionia;o__Desulfovibrionales;f__Desulfovibrionaceae;g__Desulfovibrio;s__Desulfovibrio legallii | 66.85 | 1.71 |
| Desulfovibrio legallii KHC7 | GCF_900102485.1 | 2.70 | 36 | 64.81 | p__Desulfobacterota;c__Desulfovibrionia;o__Desulfovibrionales;f__Desulfovibrionaceae;g__Desulfovibrio;s__Desulfovibrio legallii_A | 67.17 | 3.42 |
| Desulfovibrio longus DSM 6739 | GCF_000420485.1 | 3.70 | 19 | 63.66 | p__Desulfobacterota;c__Desulfovibrionia;o__Desulfovibrionales;f__Desulfovibrionaceae;g__Paucidesulfovibrio;s__Paucidesulfovibrio longus | 94.24 | 3.18 |
| Desulfovibrio magneticus str. Maddingley MBC34 | GCA_000307955.1 | 4.39 | 489 | 65.71 | p__Desulfobacterota;c__Desulfovibrionia;o__Desulfovibrionales;f__Desulfovibrionaceae;g__Solidesulfovibrio;s__Solidesulfovibrio magneticus_A | 98.99 | 4.73 |
| Desulfovibrio magneticus UBA7700 | GCA_002482565.1 | 4.22 | 217 | 65.61 | p__Desulfobacterota;c__Desulfovibrionia;o__Desulfovibrionales;f__Desulfovibrionaceae;g__Solidesulfovibrio;s__Solidesulfovibrio magneticus_B | 96.57 | 3.36 |
| Desulfovibrio mexicanus DSM 13116 | GCF_900188225.1 | 3.54 | 15 | 65.68 | p__Desulfobacterota;c__Desulfovibrionia;o__Desulfovibrionales;f__Desulfovibrionaceae;g__Humidesulfovibrio;s__Humidesulfovibrio mexicanus | 91.54 | 0.12 |
| Desulfovibrio oxyclinae DSM 11498 | GCF_000375485.1 | 3.32 | 32 | 59.12 | p__Desulfobacterota;c__Desulfovibrionia;o__Desulfovibrionales;f__Desulfovibrionaceae;g__Pseudodesulfovibrio;s__Pseudodesulfovibrio oxyclinae | 74.00 | 2.73 |
| Desulfovibrio piger ATCC 29098 | GCF_000156375.1 | 2.87 | 47 | 63.05 | p__Desulfobacterota;c__Desulfovibrionia;o__Desulfovibrionales;f__Desulfovibrionaceae;g__Desulfovibrio;s__Desulfovibrio piger | 99.91 | 0.63 |
| Desulfovibrio sp. DV | GCF_001936595.1 | 4.85 | 219 | 63.38 | p__Desulfobacterota;c__Desulfovibrionia;o__Desulfovibrionales;f__Desulfovibrionaceae;g__Solidesulfovibrio;s__Solidesulfovibrio sp001936595 | 98.25 | 4.20 |
| Desulfovibrio sp. TomC | GCF_000801335.2 | 5.07 | 84 | 61.71 | p__Desulfobacterota;c__Desulfovibrionia;o__Desulfovibrionales;f__Desulfovibrionaceae;g__Solidesulfovibrio;s__Solidesulfovibrio sp000801335 | 99.11 | 1.68 |
| Desulfovibrio sp. UBA4079 | GCA_002382645.1 | 2.86 | 208 | 68.01 | p__Desulfobacterota;c__Desulfovibrionia;o__Desulfovibrionales;f__Desulfovibrionaceae;g__Aminidesulfovibrio;s__Aminidesulfovibrio sp002382645 | 87.10 | 0.86 |
| Desulfovibrio sp. UBA6121 | GCA_002422505.1 | 3.18 | 178 | 65.01 | p__Desulfobacterota;c__Desulfovibrionia;o__Desulfovibrionales;f__Desulfovibrionaceae;g__Humidesulfovibrio;s__Humidesulfovibrio sp002422505 | 93.53 | 2.10 |
| Desulfovibrio sp. X2 | GCF_000422205.1 | 3.91 | 66 | 67.99 | p__Desulfobacterota;c__Desulfovibrionia;o__Desulfovibrionales;f__Desulfovibrionaceae;g__Desulfohalovibrio;s__Desulfohalovibrio sp000422205 | 62.87 | 1.00 |
| Desulfovibrio zosterae DSM 11974 | GCF_000425265.1 | 4.09 | 15 | 41.76 | p__Desulfobacterota;c__Desulfovibrionia;o__Desulfovibrionales;f__Desulfovibrionaceae;g__Maridesulfovibrio;s__Maridesulfovibrio zosterae | 85.35 | 2.73 |
| Desulfovibrionaceae bacterium nDJH2bin10 | GCA_015233395.1 | 3.37 | 158 | 62.81 | p__Desulfobacterota;c__Desulfovibrionia;o__Desulfovibrionales;f__Desulfovibrionaceae;g__;s__ | 94.49 | 1.19 |
| Desulfovibrionaceae bacterium nDJH8bin10 | GCA_015233065.1 | 3.29 | 255 | 62.71 | p__Desulfobacterota;c__Desulfovibrionia;o__Desulfovibrionales;f__Desulfovibrionaceae;g__;s__ | 90.74 | 2.16 |
| Desulfovibrionaceae bacterium UBA930 | GCA_002293605.1 | 2.10 | 253 | 58.27 | p__Desulfobacterota;c__Desulfovibrionia;o__Desulfovibrionales;f__Desulfovibrionaceae;g__Frigidesulfovibrio;s__Frigidesulfovibrio sp002293605 | 70.44 | 0.13 |
| Desulfurivibrio alkaliphilus AHT 2 | GCF_000092205.1 | 3.10 | 1 | 60.29 | p__Desulfobacterota;c__Desulfobulbia;o__Desulfobulbales;f__Desulfurivibrionaceae;g__Desulfurivibrio;s__Desulfurivibrio alkaliphilus | 93.55 | 1.29 |
| Desulfurivibrio sp. SURF_16 | GCA_003605035.1 | 4.22 | 118 | 57.21 | p__Desulfobacterota;c__Desulfobulbia;o__Desulfobulbales;f__Desulfurivibrionaceae;g__SURF-16;s__SURF-16 sp003605035 | 92.58 | 1.29 |
| Desulfuromonadaceae bacterium CSSed165cm_319 | GCA_007134025.1 | 2.25 | 149 | 58.5 | p__Desulfobacterota_F;c__Desulfuromonadia;o__Desulfuromonadales;f__Syntrophotaleaceae;g__SLLR01;s__SLLR01 sp007134025 | 86.13 | 1.61 |
| Desulfuromonadaceae bacterium GWB2_53_15 | GCA_001824455.1 | 1.83 | 189 | 53.47 | p__Desulfobacterota_F;c__Desulfuromonadia;o__Geobacterales;f__Pseudopelobacteraceae;g__Pseudopelobacter;s__Pseudopelobacter sp001824455 | 95.22 | 0.00 |
| Desulfuromonadaceae bacterium SZUA-401 | GCA_003246035.1 | 1.69 | 120 | 63.06 | p__Desulfobacterota_F;c__Desulfuromonadia;o__Desulfuromonadales;f__SZUA-401;g__SZUA-401;s__SZUA-401 sp003246035 | 97.40 | 3.78 |
| Desulfuromonadaceae bacterium UBA5613 | GCA_002424625.1 | 2.38 | 144 | 56.32 | p__Desulfobacterota_F;c__Desulfuromonadia;o__Desulfuromonadales;f__Trichloromonadaceae;g__UBA2197;s__UBA2197 sp002424625 | 93.35 | 3.64 |
| Desulfuromonadaceae bacterium UBA6124 | GCA_002423365.1 | 3.37 | 399 | 61.42 | p__Desulfobacterota_F;c__Desulfuromonadia;o__Desulfuromonadales;f__Trichloromonadaceae;g__Trichloromonas;s__Trichloromonas sp002423365 | 96.76 | 3.64 |

| Genome name | NCBI/IMG accession | Size, Mbp | Scaff , no. | GC, % | GTDB Taxonomy | Completeness, % | Contamination, % |
|---|---|---|---|---|---|---|---|
| Desulfuromonadales bacterium C00003068 | GCA_001751155.1 | 2.74 | 461 | 46.28 | p_Desulfobacterota_F;c_Desulfuromonadia;o_Desulfuromonadales;f_Desulfuromonadaceae;g_Desulfuromonas;s_Desulfuromonas sp001751155 | 92.26 | 0.32 |
| Desulfuromonadales bacterium CSSed11_297R1 | GCA_003561875.1 | 2.22 | 239 | 58.23 | p_Desulfobacterota_F;c_Desulfuromonadia;o_Desulfuromonadales;f_Syntrophotaleaceae;g_SLLR01;s_SLLR01 sp003561875 | 90.06 | 7.28 |
| Desulfuromonadales bacterium GT-UBC1 | GCA_003712145.1 | 4.04 | 110 | 59.89 | p_Desulfobacterota_F;c_Desulfuromonadia;o_Geobacterales;f_Geobacteraceae;g_Geobacter;s_Geobacter sp003712145 | 100.00 | 0.60 |
| Desulfuromonadales bacterium MAG_13126_9_058 | GCA_013349325.1 | 3.58 | 72 | 52.01 | p_Desulfobacterota_F;c_Desulfuromonadia;o_Geobacterales;f_Pseudopelobacteraceae;g_Pseudopelobacter;s_Pseudopelobacter sp013349325 | 65.68 | 0.00 |
| Desulfuromonadales bacterium MAG_21600_9_004 | GCA_013349405.1 | 3.43 | 60 | 51.5 | p_Desulfobacterota_F;c_Desulfuromonadia;o_Geobacterales;f_Pseudopelobacteraceae;g_Pseudopelobacter;s_Pseudopelobacter sp013349405 | 96.13 | 1.19 |
| Desulfuromonadales bacterium MAG_21601_9_030 | GCA_013349415.1 | 2.54 | 232 | 54.11 | p_Desulfobacterota_F;c_Desulfuromonadia;o_Geobacterales;f_Pseudopelobacteraceae;g_Pseudopelobacter;s_ | 95.83 | 3.01 |
| Desulfuromonas acetexigens DSM 1397 | GCF_900111775.1 | 3.68 | 41 | 60.34 | p_Desulfobacterota_F;c_Desulfuromonadia;o_Desulfuromonadales;f_Trichloromonadaceae;g_Trichloromonas;s_Trichloromonas acetexigens | 93.58 | 4.80 |
| Desulfuromonas acetoxidans DSM 684 | GCF_000167355.1 | 3.83 | 51 | 51.83 | p_Desulfobacterota_F;c_Desulfuromonadia;o_Desulfuromonadales;f_Desulfuromonadaceae;g_Desulfuromonas;s_Desulfuromonas acetoxidans | 94.17 | 0.15 |
| Desulfuromonas sp. BM302 | GCA_002868925.1 | 2.57 | 216 | 56.03 | p_Desulfobacterota_F;c_Desulfuromonadia;o_Desulfuromonadales;f_UBA2294;g_BM707;s_BM707 sp002868925 | 96.14 | 8.18 |
| Desulfuromonas sp. BM508 | GCA_002868865.1 | 2.66 | 109 | 55.18 | p_Desulfobacterota_F;c_Desulfuromonadia;o_Desulfuromonadales;f_Geopsychrobacteraceae;g_BM509;s_BM509 sp002868865 | 79.80 | 1.35 |
| Desulfuromonas sp. BM509 | GCA_002869695.1 | 3.29 | 219 | 54.98 | p_Desulfobacterota_F;c_Desulfuromonadia;o_Desulfuromonadales;f_Geopsychrobacteraceae;g_BM509;s_BM509 sp002869695 | 81.70 | 0.65 |
| Desulfuromonas sp. BM513 | GCA_002869685.1 | 3.12 | 186 | 52.53 | p_Desulfobacterota_F;c_Desulfuromonadia;o_Desulfuromonadales;f_Desulfuromonadaceae;g_Desulfuromonas;s_Desulfuromonas sp002869685 | 95.71 | 7.19 |
| Desulfuromonas sp. BM707 | GCA_002869615.1 | 3.08 | 121 | 53.07 | p_Desulfobacterota_F;c_Desulfuromonadia;o_Desulfuromonadales;f_UBA2294;g_BM707;s_BM707 sp002869615 | 98.18 | 2.73 |
| Desulfuromonas sp. BM709 | GCA_002869605.1 | 3.03 | 226 | 57.94 | p_Desulfobacterota_F;c_Desulfuromonadia;o_Desulfuromonadales;f_SZUA-401;g_SZUA-401;s_SZUA-401 sp002869605 | 90.97 | 2.26 |
| Desulfuromusa kysingii DSM 7343 | GCF_900107645.1 | 3.74 | 27 | 46.66 | p_Desulfobacterota_F;c_Desulfuromonadia;o_Desulfuromonadales;f_Geopsychrobacteraceae;g_Desulfuromusa;s_Desulfuromusa kysingii | 88.39 | 6.33 |
| Dethiosulfatarculus sandiegensis SPR | GCF_000931935.2 | 5.93 | 82 | 52.06 | p_Desulfobacterota;c_Desulfarculia;o_Desulfarculales;f_Desulfarculaceae;g_Dethiosulfatarculus;s_Dethiosulfatarculus sandiegensis | 97.67 | 2.27 |
| Dissulfurispira thermophila T55J | GCA_014701235.1 | 2.37 | 1 | 38.74 | p_Nitrospirota;c_Thermodesulfovibrionia;o_Thermodesulfovibrionales;f_UBA9935;g_UBA665;s_ | 63.54 | 0.93 |
| Elusimicrobia bacterium GWA2_69_24 | GCA_001799695.1 | 4.20 | 132 | 68.59 | p_Elusimicrobiota;c_Elusimicrobia;o_UBA1565;f_UBA1565;g_UBA10166;s_UBA10166 sp001799695 | 89.47 | 0.36 |
| Elusimicrobia bacterium NORP122 | GCA_002401485.1 | 2.91 | 191 | 54.93 | p_Elusimicrobiota;c_Elusimicrobia;o_UBA1565;f_UBA1565;g_UBA1565;s_UBA1565 sp002401485 | 91.61 | 1.29 |
| Elusimicrobia bacterium UBA10166 | GCA_003505215.1 | 4.76 | 305 | 68.17 | p_Elusimicrobiota;c_Elusimicrobia;o_UBA1565;f_UBA1565;g_UBA10166;s_UBA10166 sp001799695 | 91.60 | 1.79 |
| Elusimicrobia bacterium UBA1565 | GCA_002322475.1 | 3.60 | 408 | 62.79 | p_Elusimicrobiota;c_Elusimicrobia;o_UBA1565;f_UBA1565;g_UBA1565;s_UBA1565 sp002790095 | 91.17 | 5.22 |
| Elusimicrobia bacterium UBA9639 | GCA_003450735.1 | 4.17 | 84 | 67.93 | p_Elusimicrobiota;c_Elusimicrobia;o_UBA1565;f_UBA1565;g_UBA9639;s_UBA9639 sp003450735 | 84.82 | 3.65 |
| Elusimicrobium minutum Pei191 | GCF_000020145.1 | 1.64 | 1 | 39.95 | p_Elusimicrobiota;c_Elusimicrobia;o_Elusimicrobiales;f_Elusimicrobiaceae;g_Elusimicrobium;s_Elusimicrobium minutum | 94.54 | 1.26 |
| Endomicrobium proavitum Rsa215 | GCF_001027545.1 | 1.59 | 1 | 39.34 | p_Elusimicrobiota;c_Endomicrobia;o_Endomicrobiales;f_Endomicrobiaceae;g_Endomicrobium;s_Endomicrobium proavitum | 50.53 | 2.33 |
| Fibrobacteria bacterium nGRbin1 | GCA_015232855.1 | 5.38 | 350 | 40.73 | p_Fibrobacterota;c_Fibrobacteria;o_UBA11236;f_;g_;s_ | 78.15 | 8.39 |
| Fusobacterium periodonticum ATCC 33693 | GCA_000160475.1 | 2.62 | 53 | 27.37 | p_Fusobacteriota;c_Fusobacteriia;o_Fusobacteriales;f_Fusobacteriaceae;g_Fusobacterium;s_Fusobacterium periodonticum | 90.91 | 3.41 |
| Fusobacterium russii ATCC 25533 593A | GCA_000381725.1 | 1.94 | 41 | 28.64 | p_Fusobacteriota;c_Fusobacteriia;o_Fusobacteriales;f_Fusobacteriaceae;g_Fusobacterium;s_Fusobacterium russii | 71.11 | 1.46 |
| Gammaproteobacteria bacterium MAG_00150_gam_010 | GCA_013349855.1 | 2.85 | 486 | 49.09 | p_Proteobacteria;c_Gammaproteobacteria;o_GCF-002020875;f_GCF-002020875;g_NIOZ-UU100;s_NIOZ-UU100 sp013349825 | 97.97 | 2.58 |
| Gammaproteobacteria bacterium MAG_00160_gam_009 | GCA_013349845.1 | 2.90 | 318 | 49.11 | p_Proteobacteria;c_Gammaproteobacteria;o_GCF-002020875;f_GCF-002020875;g_NIOZ-UU100;s_NIOZ-UU100 sp013349825 | 99.41 | 0.59 |
| Gammaproteobacteria bacterium MAG_00172_gam_018 | GCA_013349825.1 | 2.87 | 274 | 48.98 | p_Proteobacteria;c_Gammaproteobacteria;o_GCF-002020875;f_GCF-002020875;g_NIOZ-UU100;s_NIOZ-UU100 sp013349825 | 99.40 | 0.00 |
| Gammaproteobacteria bacterium MAG_00188_gam_006 | GCA_013349835.1 | 2.66 | 565 | 48.83 | p_Proteobacteria;c_Gammaproteobacteria;o_GCF-002020875;f_GCF-002020875;g_NIOZ-UU100;s_NIOZ-UU100 sp013349825 | 98.71 | 0.00 |
| Gammaproteobacteria bacterium MAG_00212_gam_1 | GCA_013349805.1 | 2.10 | 955 | 48.4 | p_Proteobacteria;c_Gammaproteobacteria;o_GCF-002020875;f_GCF-002020875;g_NIOZ-UU100;s_NIOZ-UU100 sp013349825 | 99.35 | 0.00 |
| Gammaproteobacteria bacterium MAG_00215_gam_020 | GCA_013349775.1 | 2.93 | 507 | 49.02 | p_Proteobacteria;c_Gammaproteobacteria;o_GCF-002020875;f_GCF-002020875;g_NIOZ-UU100;s_NIOZ-UU100 sp013349825 | 99.08 | 3.67 |
| Gammaproteobacteria bacterium nER1bin10 | GCA_015232955.1 | 2.60 | 485 | 54.7 | p_Proteobacteria;c_Gammaproteobacteria;o_Thiohalomonadales;f_Thiohalomonadaceae;g_;s_ | 94.49 | 1.82 |
| Gammaproteobacteria bacterium NORP174 | GCA_002400775.1 | 1.50 | 52 | 46.59 | p_Proteobacteria;c_Gammaproteobacteria;o_GCA-2400775;f_GCA-2400775;g_GCA-2400775;s_GCA-2400775 sp002400775 | 99.68 | 0.65 |
| Gammaproteobacteria bacterium nPCRbin5 | GCA_015231865.1 | 3.54 | 299 | 34.87 | p_Proteobacteria;c_Gammaproteobacteria;o_GCF-002020875;f_GRL18;g_;s_ | 98.06 | 0.65 |
| Gammaproteobacteria bacterium nTSbin2 | GCA_015231435.1 | 2.87 | 223 | 52.9 | p_Proteobacteria;c_Gammaproteobacteria;o_Thiohalomonadales;f_Thiohalomonadaceae;g_;s_ | 100.00 | 0.00 |
| Gammaproteobacteria bacterium nS315bin38 | GCA_015231665.1 | 2.47 | 429 | 62.18 | p_Proteobacteria;c_Gammaproteobacteria;o_Chromatiales;f_Sedimenticolaceae;g_YD12-FULL-61-37;s_ | 99.35 | 1.29 |
| Gemmata obscuriglobus UQM 2246 | GCA_000171775.1 | 9.15 | 922 | 67.19 | p_Planctomycetota;c_Planctomycetes;o_Gemmatales;f_Gemmataceae;g_Gemmata;s_Gemmata obscuriglobus | 60.34 | 0.00 |
| Geoalkalibacter ferrihydriticus DSM 17813 | GCF_000820505.1 | 3.84 | 23 | 57.95 | p_Desulfobacterota_F;c_Desulfuromonadia;o_Desulfuromonadales;f_Geoalkalibacteraceae;g_Geoalkalibacter;s_Geoalkalibacter ferrihydriticus | 81.29 | 0.65 |
| Geoalkalibacter subterraneus Red1 | GCF_000827125.1 | 3.72 | 2 | 56.68 | p_Desulfobacterota_F;c_Desulfuromonadia;o_Desulfuromonadales;f_Geoalkalibacteraceae;g_Geoalkalibacter_A;s_Geoalkalibacter_A subterraneus | 84.82 | 0.67 |
| Geobacter anodireducens SD-1 | GCA_001628815.1 | 3.67 | 2 | 61.47 | p_Desulfobacterota_F;c_Desulfuromonadia;o_Geobacterales;f_Geobacteraceae;g_Geobacter;s_Geobacter anodireducens | 100.00 | 5.45 |
| Geobacter daltonii FRC-32 | GCF_000022265.1 | 4.30 | 1 | 53.47 | p_Desulfobacterota_F;c_Desulfuromonadia;o_Geobacterales;f_Geobacteraceae;g_Geotalea;s_Geotalea daltonii | 92.92 | 0.91 |
| Geobacter lovleyi SZ | GCF_000020385.1 | 3.99 | 2 | 54.74 | p_Desulfobacterota_F;c_Desulfuromonadia;o_Geobacterales;f_Pseudopelobacteraceae;g_Trichlorobacter;s_Trichlorobacter lovleyi | 85.00 | 0.81 |
| Geobacter metallireducens GS-15 | GCF_000012925.1 | 4.01 | 2 | 59.49 | p_Desulfobacterota_F;c_Desulfuromonadia;o_Geobacterales;f_Geobacteraceae;g_Geobacter;s_Geobacter metallireducens | 95.24 | 0.00 |
| Geobacter pelophilus Drf2 | GCF_002117975.1 | 4.34 | 2 | 61.03 | p_Desulfobacterota_F;c_Desulfuromonadia;o_Geobacterales;f_Geobacteraceae;g_Geomonas;s_Geomonas pelophila | 95.85 | 3.18 |
| Geobacter sp. L1geo | GCA_003574895.1 | 4.48 | 39 | 54.15 | p_Desulfobacterota_F;c_Desulfuromonadia;o_Geobacterales;f_Pseudopelobacteraceae;g_Pseudopelobacter;s_Pseudopelobacter sp003574895 | 90.91 | 2.73 |
| Geobacter sp. palsa_1151 | GCA_003151775.1 | 2.95 | 145 | 56.96 | p_Desulfobacterota_F;c_Desulfuromonadia;o_Geobacterales;f_Pseudopelobacteraceae;g_Pseudopelobacter;s_Pseudopelobacter sp003151775 | 95.35 | 3.64 |
| Geobacter sp. UBA1603 | GCA_002322035.1 | 2.35 | 108 | 56.78 | p_Desulfobacterota_F;c_Desulfuromonadia;o_Geobacterales;f_Pseudopelobacteraceae;g_Pseudopelobacter;s_Pseudopelobacter sp002322035 | 92.92 | 4.55 |
| Geobacter sp. UBA698 | GCA_002298925.1 | 4.33 | 152 | 55.97 | p_Desulfobacterota_F;c_Desulfuromonadia;o_Geobacterales;f_Pseudopelobacteraceae;g_Pseudopelobacter;s_Pseudopelobacter sp002298925 | 99.35 | 0.65 |
| Geobacter sulfurreducens PCA | GCF_000007985.2 | 3.81 | 1 | 60.94 | p_Desulfobacterota_F;c_Desulfuromonadia;o_Geobacterales;f_Geobacteraceae;g_Geobacter;s_Geobacter sulfurreducens | 99.91 | 0.89 |
| Geobacter thiogenes ATCC BAA-34 | GCF_900167465.1 | 3.57 | 42 | 52.83 | p_Desulfobacterota_F;c_Desulfuromonadia;o_Geobacterales;f_Pseudopelobacteraceae;g_Trichlorobacter;s_Trichlorobacter thiogenes | 70.75 | 2.98 |
| Geobacter toluenoxydans JCM 15764 | GCA_001311985.1 | 4.21 | 77 | 53.59 | p_Desulfobacterota_F;c_Desulfuromonadia;o_Geobacterales;f_Geobacteraceae;g_Geotalea;s_Geotalea toluenoxydans | 94.94 | 2.78 |
| Geobacter uraniireducens Rf4 | GCF_000016745.1 | 5.14 | 1 | 54.24 | p_Desulfobacterota_F;c_Desulfuromonadia;o_Geobacterales;f_Geobacteraceae;g_Geotalea;s_Geotalea uraniireducens | 94.29 | 3.69 |
| Geobacteraceae bacterium GWC2_53_11 | GCA_001802645.1 | 4.28 | 91 | 53 | p_Desulfobacterota_F;c_Desulfuromonadia;o_Geobacterales;f_Pseudopelobacteraceae;g_Pseudopelobacter;s_Pseudopelobacter sp001802645 | 91.55 | 7.83 |
| Geobacteraceae bacterium GWC2_55_20 | GCA_001802125.1 | 4.55 | 186 | 54.41 | p_Desulfobacterota_F;c_Desulfuromonadia;o_Geobacterales;f_Pseudopelobacteraceae;g_Pseudopelobacter;s_Pseudopelobacter sp001802125 | 91.36 | 2.73 |
| Geomonas oryzae S43 | GCF_004117875.1 | 4.93 | 18 | 61.24 | p_Desulfobacterota_F;c_Desulfuromonadia;o_Geobacterales;f_Geobacteraceae;g_Geomonas;s_Geomonas oryzae | 95.85 | 2.83 |
| Geothermobacter sp. EPR-M | GCF_002093115.1 | 3.73 | 59 | 59.39 | p_Desulfobacterota_F;c_Desulfuromonadia;o_Desulfuromonadales;f_Geothermobacteraceae;g_PPFX01;s_PPFX01 sp002093115 | 95.59 | 3.28 |
| Geothermobacter sp. HR-1 | GCF_002898515.1 | 3.84 | 73 | 59.23 | p_Desulfobacterota_F;c_Desulfuromonadia;o_Desulfuromonadales;f_Geothermobacteraceae;g_PPFX01;s_PPFX01 sp002898515 | 99.26 | 0.65 |

| Genome name | NCBI/IMG accession | Size, Mbp | Scaff, no. | GC, % | GTDB Taxonomy | Completeness, % | Contamination, % |
|---|---|---|---|---|---|---|---|
| Ghiorsea bivora TAG-1 | GCA_000744415.1 | 2.16 | 13 | 42.68 | p__Proteobacteria;c__Zetaproteobacteria;o__Mariprofundales;f__Mariprofundaceae;g__Ghiorsea;s__Ghiorsea bivora | 99.35 | 1.29 |
| Gimesia maris DSM 8797 | GCA_000181475.1 | 7.78 | 125 | 50.45 | p__Planctomycetota;c__Planctomycetes;o__Planctomycetales;f__Planctomycetaceae;g__Gimesia;s__Gimesia maris | 85.91 | 0.45 |
| Halodesulfovibrio aestuarii DSM 17919 | GCF_000384815.1 | 3.55 | 21 | 45.11 | p__Desulfobacterota;c__Desulfovibrionia;o__Desulfovibrionales;f__Desulfovibrionaceae;g__Halodesulfovibrio;s__Halodesulfovibrio aestuarii | 96.42 | 1.68 |
| Halodesulfovibrio marinisediminis DSM 17456 | GCF_900129975.1 | 3.71 | 13 | 44.96 | p__Desulfobacterota;c__Desulfovibrionia;o__Desulfovibrionales;f__Desulfovibrionaceae;g__Halodesulfovibrio;s__Halodesulfovibrio marinisediminis | 80.69 | 2.26 |
| Halodesulfovibrio spirochaetisodalis JC271 | GCF_001672295.1 | 3.61 | 59 | 46.19 | p__Desulfobacterota;c__Desulfovibrionia;o__Desulfovibrionales;f__Desulfovibrionaceae;g__Halodesulfovibrio;s__Halodesulfovibrio spirochaetisodalis | 95.80 | 2.52 |
| Ilyobacter polytropus DSM 2926 | GCA_000165505.1 | 3.13 | 3 | 34.37 | p__Fusobacteriota;c__Fusobacteriia;o__Fusobacteriales;f__Fusobacteriaceae;g__Ilyobacter;s__Ilyobacter polytropus | 89.68 | 5.59 |
| Isosphaera pallida ATCC 43644 | GCA_000186345.1 | 5.53 | 2 | 62.49 | p__Planctomycetota;c__Planctomycetes;o__Isosphaerales;f__Isosphaeraceae;g__Isosphaera;s__Isosphaera pallida | 66.54 | 0.00 |
| Latescibacteria bacterium SCGC AAA252-B13 | 2264867252 | 1.76 | 138 | 40.86 | p__Latescibacterota;c__Latescibacteria;o__Latescibacterales;f__Latescibacteraceae;g__Latescibacter;s__ | 43.99 | 0.00 |
| Leptospirillum ferriphilum DSM 14647 | GCF_000755505.1 | 2.41 | 18 | 54.05 | p__Nitrospirota_A;c__Leptospirillia;o__Leptospirillales;f__Leptospirillaceae;g__Leptospirillum_A;s__Leptospirillum_A ferriphilum | 68.81 | 0.62 |
| Leptospirillum ferriphilum ML-04 | GCF_000299235.1 | 2.41 | 1 | 54.55 | p__Nitrospirota_A;c__Leptospirillia;o__Leptospirillales;f__Leptospirillaceae;g__Leptospirillum_A;s__Leptospirillum_A rubarum | 98.58 | 0.91 |
| Leptospirillum ferrooxidans C2-3 | GCF_000284315.1 | 2.56 | 1 | 50.05 | p__Nitrospirota_A;c__Leptospirillia;o__Leptospirillales;f__Leptospirillaceae;g__Leptospirillum;s__Leptospirillum ferrooxidans | 55.23 | 0.00 |
| Leptotrichia buccalis C-1013-b | GCA_000023905.1 | 2.47 | 1 | 29.65 | p__Fusobacteriota;c__Fusobacteriia;o__Fusobacteriales;f__Leptotrichiaceae;g__Leptotrichia;s__Leptotrichia buccalis | 62.13 | 5.10 |
| Magnetococcales bacterium DC0425bin3 | GCA_002753665.1 | 3.70 | 230 | 65.4 | p__Proteobacteria;c__Magnetococcia;o__Magnetococcales;f__DC0425bin3;g__DC0425bin3;s__DC0425bin3 sp002753665 | 99.35 | 0.65 |
| Magnetococcales bacterium DCbin2 | GCA_002753615.1 | 3.36 | 201 | 51.85 | p__Proteobacteria;c__Magnetococcia;o__Magnetococcales;f__UBA8363;g__UBA8363;s__UBA8363 sp002753615 | 99.40 | 0.00 |
| Magnetococcales bacterium DCbin4 | GCA_002753735.1 | 4.52 | 148 | 54.25 | p__Proteobacteria;c__Magnetococcia;o__Magnetococcales;f__UBA8363;g__UBA8363;s__UBA8363 sp002753735 | 99.35 | 0.65 |
| Magnetococcales bacterium DH2_bin20 | GCA_011089865.1 | 5.01 | 525 | 52.96 | p__Proteobacteria;c__Magnetococcia;o__Magnetococcales;f__;g__;s__ | 100.00 | 0.00 |
| Magnetococcales bacterium DH2_bin6 | GCA_011089965.1 | 3.66 | 80 | 56.76 | p__Proteobacteria;c__Magnetococcia;o__Magnetococcales;f__Magnetaquicoccaceae;g__JAANAU01;s__JAANAU01 sp011089965 | 99.35 | 3.39 |
| Magnetococcales bacterium ER1bin7 | GCA_002753565.1 | 3.87 | 92 | 52.3 | p__Proteobacteria;c__Magnetococcia;o__Magnetococcales;f__UBA8363;g__GCA-2753565;s__GCA-2753565 sp002753565 | 98.98 | 0.00 |
| Magnetococcales bacterium HA3dbin1 | GCA_002753515.1 | 4.33 | 134 | 53.32 | p__Proteobacteria;c__Magnetococcia;o__Magnetococcales;f__UBA8363;g__UBA8363;s__UBA8363 sp002753515 | 100.00 | 0.00 |
| Magnetococcales bacterium HA3dbin3 | GCA_002753495.1 | 2.90 | 316 | 61.72 | p__Proteobacteria;c__Magnetococcia;o__Magnetococcales;f__DC0425bin3;g__HA3dbin3;s__HA3dbin3 sp002753495 | 98.70 | 0.65 |
| Magnetococcales bacterium HAa3bin1 | GCA_002753595.1 | 4.35 | 118 | 53.21 | p__Proteobacteria;c__Magnetococcia;o__Magnetococcales;f__UBA8363;g__UBA8363;s__UBA8363 sp002753515 | 99.50 | 1.24 |
| Magnetococcales bacterium HCHbin5 | GCA_002753505.1 | 4.19 | 200 | 56.97 | p__Proteobacteria;c__Magnetococcia;o__Magnetococcales;f__Magnetaquicoccaceae;g__HCHbin5;s__HCHbin5 sp002753505 | 99.88 | 0.00 |
| Magnetococcales bacterium MAG_21055_mgc_1 | GCA_013349385.1 | 3.59 | 930 | 52.41 | p__Proteobacteria;c__Magnetococcia;o__Magnetococcales;f__UBA8363;g__UBA8363;s__UBA8363 sp013349385 | 100.00 | 0.59 |
| Magnetococcales bacterium nARSLQbin1 | GCA_015234115.1 | 3.31 | 161 | 58.83 | p__Proteobacteria;c__Magnetococcia;o__Magnetococcales;f__Magnetaquicoccaceae;g__JAANAU01;s__ | 99.50 | 0.00 |
| Magnetococcales bacterium nCLbin10 | GCA_015233785.1 | 3.36 | 364 | 55.62 | p__Proteobacteria;c__Magnetococcia;o__Magnetococcales;f__;g__;s__ | 99.17 | 1.65 |
| Magnetococcales bacterium nCLbin6 | GCA_015234045.1 | 3.65 | 117 | 47.56 | p__Proteobacteria;c__Magnetococcia;o__Magnetococcales;f__;g__;s__ | 100.00 | 0.00 |
| Magnetococcales bacterium nDC0425bin2 | GCA_015233945.1 | 3.04 | 364 | 55.86 | p__Proteobacteria;c__Magnetococcia;o__Magnetococcales;f__Magnetaquicoccaceae;g__HCHbin5;s__HCHbin5 sp002753505 | 100.00 | 0.75 |
| Magnetococcales bacterium nDC0425bin4 | GCA_015233965.1 | 3.89 | 165 | 65.39 | p__Proteobacteria;c__Magnetococcia;o__Magnetococcales;f__DC0425bin3;g__DC0425bin3;s__DC0425bin3 sp002753665 | 100.00 | 0.00 |
| Magnetococcales bacterium nDCbin4 | GCA_015234015.1 | 4.49 | 149 | 54.13 | p__Proteobacteria;c__Magnetococcia;o__Magnetococcales;f__UBA8363;g__UBA8363;s__UBA8363 sp002753735 | 99.35 | 1.61 |
| Magnetococcales bacterium nDH2bin6 | GCA_015233935.1 | 4.54 | 414 | 52.7 | p__Proteobacteria;c__Magnetococcia;o__Magnetococcales;f__;g__;s__ | 99.41 | 0.00 |
| Magnetococcales bacterium nDH2bin7 | GCA_015233905.1 | 1.95 | 74 | 56.31 | p__Proteobacteria;c__Magnetococcia;o__Magnetococcales;f__Magnetaquicoccaceae;g__JAANAU01;s__JAANAU01 sp011089965 | 100.00 | 1.39 |
| Magnetococcales bacterium nDJH13bin19 | GCA_015233895.1 | 3.71 | 65 | 41.57 | p__Nitrospirota;c__Thermodesulfovibrionia;o__Thermodesulfovibrionales;f__Magnetobacteriaceae;g__;s__ | 79.81 | 1.71 |
| Magnetococcales bacterium nDJH15bin4 | GCA_015233515.1 | 4.20 | 701 | 51.2 | p__Desulfobacterota;c__Desulfarculia_A;o__Adiutricales;f__;g__;s__ | 94.87 | 3.85 |
| Magnetococcales bacterium nDJH8bin2 | GCA_015233035.1 | 4.53 | 434 | 57.45 | p__Proteobacteria;c__Magnetococcia;o__Magnetococcales;f__Magnetaquicoccaceae;g__HCHbin5;s__ | 98.82 | 0.30 |
| Magnetococcales bacterium nER1bin1 | GCA_015232915.1 | 3.66 | 47 | 52.32 | p__Proteobacteria;c__Magnetococcia;o__Magnetococcales;f__UBA8363;g__GCA-2753565;s__GCA-2753565 sp002753565 | 98.51 | 0.50 |
| Magnetococcales bacterium nER1bin6 | GCA_015232865.1 | 3.09 | 102 | 51.04 | p__Proteobacteria;c__Magnetococcia;o__Magnetococcales;f__UBA8363;g__GCA-2753565;s__ | 99.41 | 0.07 |
| Magnetococcales bacterium nGRbin4 | GCA_015232795.1 | 2.87 | 466 | 42 | p__Proteobacteria;c__Magnetococcia;o__Magnetococcales;f__UBA8363;g__GCA-2753565;s__ | 100.00 | 0.00 |
| Magnetococcales bacterium nHA1bin2 | GCA_015232825.1 | 3.03 | 270 | 50.29 | p__Proteobacteria;c__Magnetococcia;o__Magnetococcales;f__Magnetaquicoccaceae;g__JAANAU01;s__ | 99.35 | 2.15 |
| Magnetococcales bacterium nHA3dbin1 | GCA_015232815.1 | 4.47 | 203 | 52.89 | p__Proteobacteria;c__Magnetococcia;o__Magnetococcales;f__UBA8363;g__UBA8363;s__UBA8363 sp002753515 | 99.35 | 0.65 |
| Magnetococcales bacterium nHA3dbin2 | GCA_015232755.1 | 3.55 | 358 | 61.56 | p__Proteobacteria;c__Magnetococcia;o__Magnetococcales;f__DC0425bin3;g__HA3dbin3;s__HA3dbin3 sp002753495 | 99.33 | 0.00 |
| Magnetococcales bacterium nHA4bin1 | GCA_015232765.1 | 2.44 | 414 | 50.34 | p__Proteobacteria;c__Magnetococcia;o__Magnetococcales;f__Magnetaquicoccaceae;g__JAANAU01;s__ | 99.35 | 0.65 |
| Magnetococcales bacterium nHA4bin8 | GCA_015232705.1 | 2.05 | 382 | 58.19 | p__Proteobacteria;c__Magnetococcia;o__Magnetococcales;f__Magnetaquicoccaceae;g__JAANAU01;s__ | 99.70 | 0.00 |
| Magnetococcales bacterium nHA5abin2 | GCA_015232685.1 | 3.24 | 229 | 52.36 | p__Proteobacteria;c__Magnetococcia;o__Magnetococcales;f__UBA8363;g__UBA8363;s__UBA8363 sp002753515 | 99.68 | 0.00 |
| Magnetococcales bacterium nHA5abin3 | GCA_015232645.1 | 3.39 | 426 | 54.67 | p__Proteobacteria;c__Magnetococcia;o__Magnetococcales;f__UBA8363;g__UBA8363;s__ | 100.00 | 0.75 |
| Magnetococcales bacterium nHAa3bin1 | GCA_015232675.1 | 3.00 | 97 | 53.21 | p__Proteobacteria;c__Magnetococcia;o__Magnetococcales;f__UBA8363;g__UBA8363;s__UBA8363 sp002753515 | 98.71 | 0.00 |
| Magnetococcales bacterium nHCHbin1 | GCA_015232635.1 | 3.81 | 295 | 56.77 | p__Proteobacteria;c__Magnetococcia;o__Magnetococcales;f__Magnetaquicoccaceae;g__HCHbin5;s__HCHbin5 sp002753505 | 100.00 | 1.09 |
| Magnetococcales bacterium nJSWbin1 | GCA_015232405.1 | 5.86 | 335 | 42.2 | p__Proteobacteria;c__Magnetococcia;o__Magnetococcales;f__UBA8363;g__GCA-2753565;s__ | 100.00 | 0.00 |
| Magnetococcales bacterium nJSWbin2 | GCA_015232395.1 | 4.94 | 170 | 55.33 | p__Proteobacteria;c__Magnetococcia;o__Magnetococcales;f__;g__;s__ | 99.35 | 0.00 |
| Magnetococcales bacterium nJSWbin3 | GCA_015232365.1 | 2.84 | 103 | 44.6 | p__Proteobacteria;c__Magnetococcia;o__Magnetococcales;f__UBA8363;g__GCA-2753565;s__ | 99.45 | 0.82 |
| Magnetococcales bacterium nKLKbin4 | GCA_015232265.1 | 4.03 | 225 | 54.24 | p__Proteobacteria;c__Magnetococcia;o__Magnetococcales;f__UBA8363;g__UBA8363;s__UBA8363 sp002753735 | 100.00 | 0.62 |
| Magnetococcales bacterium nMBPbin6 | GCA_015232245.1 | 3.70 | 481 | 59.23 | p__Proteobacteria;c__Magnetococcia;o__Magnetococcales;f__Magnetaquicoccaceae;g__JAANAU01;s__ | 94.17 | 2.73 |
| Magnetococcales bacterium nMYbin5 | GCA_015232135.1 | 2.25 | 376 | 56.15 | p__Proteobacteria;c__Magnetococcia;o__Magnetococcales;f__DC0425bin3;g__HA3dbin3;s__ | 99.35 | 0.65 |
| Magnetococcales bacterium nNGHbin13 | GCA_015231965.1 | 3.82 | 314 | 63.09 | p__Proteobacteria;c__Magnetococcia;o__;f__;g__;s__ | 99.50 | 0.25 |
| Magnetococcales bacterium nNGHbin14 | GCA_015231925.1 | 4.51 | 358 | 60.22 | p__Proteobacteria;c__Magnetococcia;o__Magnetococcales;f__;g__;s__ | 100.00 | 0.20 |
| Magnetococcales bacterium nNGHbin2 | GCA_015231915.1 | 4.02 | 133 | 61.98 | p__Proteobacteria;c__Magnetococcia;o__Magnetococcales;f__Magnetaquicoccaceae;g__WMHbin3;s__ | 96.82 | 4.77 |
| Magnetococcales bacterium nQXH1bin1 | GCA_015231795.1 | 3.63 | 23 | 59.95 | p__Proteobacteria;c__Alphaproteobacteria;o__Rhodospirillales;f__WMHbin7;g__WMHbin7;s__ | 99.42 | 0.00 |
| Magnetococcales bacterium nQXH2bin1 | GCA_015231775.1 | 3.82 | 389 | 61.35 | p__Proteobacteria;c__Magnetococcia;o__Magnetococcales;f__Magnetaquicoccaceae;g__JAANAU01;s__ | 96.13 | 0.65 |
| Magnetococcales bacterium nQXH2bin5 | GCA_015231755.1 | 3.84 | 44 | 59.16 | p__Proteobacteria;c__Magnetococcia;o__Magnetococcales;f__Magnetaquicoccaceae;g__JAANAU01;s__ | 99.17 | 0.00 |
| Magnetococcales bacterium nW5bin1 | GCA_015231375.1 | 4.10 | 114 | 59.91 | p__Proteobacteria;c__Magnetococcia;o__Magnetococcales;f__Magnetaquicoccaceae;g__JAANAU01;s__ | 92.42 | 0.91 |

| Genome name | NCBI/IMG accession | Size, Mbp | Scaff, no. | GC, % | GTDB Taxonomy | Completeness, % | Contamination, % |
|---|---|---|---|---|---|---|---|
| Magnetococcales bacterium nW5bin3 | GCA_015231295.1 | 3.79 | 353 | 59.17 | p_Proteobacteria;c_Magnetococcia;o_Magnetococcales;f_Magnetaquicoccaceae;g_JAANAU01;s_ | 98.74 | 0.00 |
| Magnetococcales bacterium nwagbin5 | GCA_015231305.1 | 3.96 | 238 | 38.49 | p_Proteobacteria;c_Magnetococcia;o_Magnetococcales;f_UBA8363;g_GCA-2753565;s_ | 96.75 | 4.27 |
| Magnetococcales bacterium nwalbin5 | GCA_015231265.1 | 3.82 | 459 | 55.87 | p_Proteobacteria;c_Magnetococcia;o_;f_;g_;s_ | 99.35 | 1.29 |
| Magnetococcales bacterium nWMHbin1 | GCA_015231215.1 | 4.37 | 116 | 54.97 | p_Proteobacteria;c_Magnetococcia;o_Magnetococcales;f_Magnetaquicoccaceae;g_Magnetaquicoccus;s_Magnetaquicoccus sp002753135 | 97.66 | 0.71 |
| Magnetococcales bacterium nWMHbin2 | GCA_015231175.1 | 3.81 | 175 | 57.17 | p_Proteobacteria;c_Magnetococcia;o_Magnetococcales;f_DC0425bin3;g_HA3dbin3;s_ | 100.00 | 0.00 |
| Magnetococcales bacterium nWMHbin3 | GCA_015231235.1 | 4.97 | 167 | 61.57 | p_Proteobacteria;c_Magnetococcia;o_Magnetococcales;f_Magnetaquicoccaceae;g_WMHbin3;s_WMHbin3 sp002753185 | 100.00 | 0.91 |
| Magnetococcales bacterium nWMHbin4 | GCA_015231205.1 | 4.16 | 207 | 54.29 | p_Proteobacteria;c_Magnetococcia;o_Magnetococcales;f_UBA8363;g_UBA8363;s_UBA8363 sp002753735 | 99.35 | 2.26 |
| Magnetococcales bacterium nWRX1bin1 | GCA_015229095.1 | 2.63 | 200 | 59.27 | p_Proteobacteria;c_Magnetococcia;o_Magnetococcales;f_Magnetaquicoccaceae;g_HCHbin5;s_ | 100.00 | 0.00 |
| Magnetococcales bacterium nWRX1bin11 | GCA_015229135.1 | 3.56 | 254 | 60.94 | p_Proteobacteria;c_Magnetococcia;o_Magnetococcales;f_Magnetaquicoccaceae;g_JAANAU01;s_ | 100.00 | 0.00 |
| Magnetococcales bacterium nWRX1bin12 | GCA_015229105.1 | 2.77 | 199 | 62.96 | p_Proteobacteria;c_Magnetococcia;o_Magnetococcales;f_;g_;s_ | 95.27 | 1.78 |
| Magnetococcales bacterium nWRX1bin3 | GCA_015229115.1 | 3.15 | 184 | 54.88 | p_Proteobacteria;c_Magnetococcia;o_Magnetococcales;f_;g_;s_ | 100.00 | 0.07 |
| Magnetococcales bacterium nWRX1bin5 | GCA_015229055.1 | 4.55 | 173 | 54.4 | p_Proteobacteria;c_Magnetococcia;o_Magnetococcales;f_UBA8363;g_UBA8363;s_UBA8363 sp002753735 | 100.00 | 1.19 |
| Magnetococcales bacterium nWRX1bin6 | GCA_015229045.1 | 4.08 | 283 | 57.46 | p_Proteobacteria;c_Magnetococcia;o_Magnetococcales;f_DC0425bin3;g_HA3dbin3;s_ | 98.11 | 0.00 |
| Magnetococcales bacterium nWRX2bin6 | GCA_015229035.1 | 4.72 | 210 | 54.25 | p_Proteobacteria;c_Magnetococcia;o_Magnetococcales;f_UBA8363;g_UBA8363;s_UBA8363 sp002753735 | 99.91 | 0.07 |
| Magnetococcales bacterium nWRX3bin10 | GCA_015229005.1 | 3.28 | 173 | 62.87 | p_Proteobacteria;c_Magnetococcia;o_Magnetococcales;f_;g_;s_ | 99.96 | 0.00 |
| Magnetococcales bacterium nWRX3bin11 | GCA_015228995.1 | 1.96 | 287 | 53.76 | p_Proteobacteria;c_Magnetococcia;o_Magnetococcales;f_;g_;s_ | 99.50 | 0.50 |
| Magnetococcales bacterium nWRX3bin12 | GCA_015228975.1 | 3.75 | 234 | 52.93 | p_Proteobacteria;c_Magnetococcia;o_Magnetococcales;f_UBA8363;g_UBA8363;s_ | 99.35 | 0.65 |
| Magnetococcales bacterium nWRX3bin2 | GCA_015228935.1 | 4.71 | 197 | 54.61 | p_Proteobacteria;c_Magnetococcia;o_Magnetococcales;f_DC0425bin3;g_HA3dbin3;s_ | 99.17 | 0.62 |
| Magnetococcales bacterium nWRX3bin6 | GCA_015228925.1 | 4.25 | 219 | 52.02 | p_Proteobacteria;c_Magnetococcia;o_Magnetococcales;f_UBA8363;g_UBA8363;s_ | 100.00 | 0.78 |
| Magnetococcales bacterium nWRX3bin7 | GCA_015228895.1 | 2.89 | 501 | 61.11 | p_Proteobacteria;c_Magnetococcia;o_Magnetococcales;f_Magnetaquicoccaceae;g_WMHbin3;s_ | 94.35 | 2.80 |
| Magnetococcales bacterium nYD0423bin2 | GCA_015228815.1 | 4.22 | 97 | 56.86 | p_Proteobacteria;c_Magnetococcia;o_Magnetococcales;f_UBA8363;g_UBA8363;s_ | 98.39 | 1.94 |
| Magnetococcales bacterium nYD0423bin3 | GCA_015228775.1 | 4.50 | 146 | 55.48 | p_Proteobacteria;c_Magnetococcia;o_Magnetococcales;f_Magnetaquicoccaceae;g_Magnetaquicoccus;s_Magnetaquicoccus sp002753095 | 98.06 | 4.54 |
| Magnetococcales bacterium nYD0425bin13 | GCA_015228825.1 | 4.22 | 231 | 55.36 | p_Proteobacteria;c_Magnetococcia;o_Magnetococcales;f_Magnetaquicoccaceae;g_Magnetaquicoccus;s_Magnetaquicoccus sp002753095 | 95.91 | 6.55 |
| Magnetococcales bacterium WMHbin1 | GCA_002753215.1 | 4.38 | 242 | 54.3 | p_Proteobacteria;c_Magnetococcia;o_Magnetococcales;f_UBA8363;g_UBA8363;s_UBA8363 sp002753735 | 100.00 | 0.00 |
| Magnetococcales bacterium WMHbin3 | GCA_002753185.1 | 4.60 | 157 | 61.61 | p_Proteobacteria;c_Magnetococcia;o_Magnetococcales;f_Magnetaquicoccaceae;g_WMHbin3;s_WMHbin3 sp002753185 | 96.76 | 2.73 |
| Magnetococcales bacterium WMHbinv6 | GCA_002753135.1 | 3.84 | 80 | 55.32 | p_Proteobacteria;c_Magnetococcia;o_Magnetococcales;f_Magnetaquicoccaceae;g_Magnetaquicoccus;s_Magnetaquicoccus sp002753135 | 99.03 | 3.87 |
| Magnetococcales bacterium YD0425bin7 | GCA_002753095.1 | 3.58 | 177 | 55.67 | p_Proteobacteria;c_Magnetococcia;o_Magnetococcales;f_Magnetaquicoccaceae;g_Magnetaquicoccus;s_Magnetaquicoccus sp002753095 | 95.51 | 0.37 |
| Magnetococcus marinus MC-1 | GCA_000014865.1 | 4.72 | 1 | 54.17 | p_Proteobacteria;c_Magnetococcia;o_Magnetococcales;f_Magnetococcaceae;g_Magnetococcus;s_Magnetococcus marinus | 97.94 | 1.13 |
| Magnetofaba australis IT-1 | GCA_002109495.1 | 4.99 | 21 | 61.3 | p_Proteobacteria;c_Magnetococcia;o_Magnetococcales;f_Magnetococcaceae;g_Magnetofaba;s_Magnetofaba australis | 99.41 | 0.00 |
| Magnetospira sp. QH-2 | GCF_000968135.1 | 4.05 | 2 | 59.44 | p_Proteobacteria;c_Alphaproteobacteria;o_Rhodospirillales;f_Magnetospiraceae;g_Magnetospira;s_Magnetospira sp000968135 | 98.32 | 1.68 |
| Magnetospirillum caucaseum SO-1 | GCF_000342045.1 | 4.86 | 236 | 65.99 | p_Proteobacteria;c_Alphaproteobacteria;o_Rhodospirillales;f_Magnetospirillaceae;g_Phaeospirillum;s_Phaeospirillum caucaseum | 96.06 | 2.73 |
| Magnetospirillum gryphiswaldense MSR-1 v2 | GCF_000513295.1 | 4.37 | 1 | 63.28 | p_Proteobacteria;c_Alphaproteobacteria;o_Rhodospirillales;f_Magnetospirillaceae;g_Magnetospirillum;s_Magnetospirillum gryphiswaldense | 83.95 | 3.23 |
| Magnetospirillum kuznetsovii LBB-42 | GCA_003284725.1 | 4.41 | 69 | 63.44 | p_Proteobacteria;c_Alphaproteobacteria;o_Rhodospirillales;f_Magnetospirillaceae;g_Phaeospirillum;s_Phaeospirillum kuznetsovii | 99.94 | 1.82 |
| Magnetospirillum magneticum AMB-1 | GCF_000009985.1 | 4.97 | 1 | 65.09 | p_Proteobacteria;c_Alphaproteobacteria;o_Rhodospirillales;f_Magnetospirillaceae;g_Phaeospirillum;s_Phaeospirillum magneticum | 97.05 | 2.58 |
| Magnetospirillum magnetotacticum MS-1 | GCF_000829825.1 | 4.52 | 36 | 63.56 | p_Proteobacteria;c_Alphaproteobacteria;o_Rhodospirillales;f_Magnetospirillaceae;g_Phaeospirillum;s_Phaeospirillum magnetotacticum | 99.35 | 1.29 |
| Magnetospirillum marisnigri SP-1 | GCF_001650715.1 | 4.62 | 131 | 64.73 | p_Proteobacteria;c_Alphaproteobacteria;o_Rhodospirillales;f_Magnetospirillaceae;g_Phaeospirillum;s_Phaeospirillum marisnigri | 98.06 | 2.74 |
| Magnetospirillum moscoviense BB-1 | GCF_001650635.1 | 4.16 | 207 | 65.18 | p_Proteobacteria;c_Alphaproteobacteria;o_Rhodospirillales;f_Magnetospirillaceae;g_Magnetospirillum;s_Magnetospirillum moscoviense | 70.17 | 0.00 |
| Magnetovibrio blakemorei MV-1 | GCF_001746755.1 | 3.64 | 91 | 54.29 | p_Proteobacteria;c_Alphaproteobacteria;o_Rhodospirillales;f_Magnetovibrionaceae;g_Magnetovibrio;s_Magnetovibrio blakemorei | 67.54 | 1.75 |
| Magnetovibrio sp. ARS8 | GCA_002686765.1 | 2.02 | 197 | 59.64 | p_Proteobacteria;c_Alphaproteobacteria;o_Rhodospirillales;f_2-02-FULL-58-16;g_GCA-2686765;s_GCA-2686765 sp002686765 | 99.03 | 0.91 |
| Mailhella massiliensis Marseille-P3199 | GCF_900155525.1 | 3.47 | 28 | 59.08 | p_Desulfobacterota;c_Desulfovibrionia;o_Desulfovibrionales;f_Desulfovibrionaceae;g_Mailhella;s_Mailhella massiliensis | 98.32 | 2.94 |
| Malonomonas rubra DSM 5091 | GCF_900142125.1 | 3.96 | 42 | 52.46 | p_Desulfobacterota_F;c_Desulfuromonadia;o_Desulfuromonadales;f_Geopsychrobacteraceae;g_Malonomonas;s_Malonomonas rubra | 70.43 | 2.27 |
| Malonomonas rubra SZUA-375 | GCA_003247215.1 | 3.13 | 149 | 52.49 | p_Desulfobacterota_F;c_Desulfuromonadia;o_Desulfuromonadales;f_Geopsychrobacteraceae;g_Malonomonas;s_Malonomonas rubra_A | 56.35 | 0.97 |
| Mariprofundus ferrinatatus CP-8 | GCA_002795825.1 | 2.30 | 1 | 53.66 | p_Proteobacteria;c_Zetaproteobacteria;o_Mariprofundales;f_Mariprofundaceae;g_Mariprofundus;s_Mariprofundus ferrinatatus | 100.00 | 1.45 |
| Mariprofundus ferrooxydans M34 | GCA_000379405.1 | 2.74 | 36 | 53.93 | p_Proteobacteria;c_Zetaproteobacteria;o_Mariprofundales;f_Mariprofundaceae;g_Mariprofundus;s_Mariprofundus ferrooxydans | 100.00 | 0.46 |
| Mariprofundus micogutta ET2 | GCA_001895085.1 | 2.50 | 59 | 48.76 | p_Proteobacteria;c_Zetaproteobacteria;o_Mariprofundales;f_Mariprofundaceae;g_Mariprofundus;s_Mariprofundus micogutta | 99.39 | 0.00 |
| Nitrospina gracilis 3211 | GCA_000341545.2 | 3.08 | 4 | 56.21 | p_Nitrospinota;c_Nitrospinia;o_Nitrospinales;f_Nitrospinaceae;g_Nitrospina;s_Nitrospina gracilis | 85.45 | 0.50 |
| Nitrospina sp. NAT278 | GCA_002697825.1 | 1.78 | 90 | 37.06 | p_Nitrospinota;c_Nitrospinia;o_Nitrospinales;f_Nitrospinaceae;g_UBA8687;s_UBA8687 sp002697825 | 55.59 | 2.41 |
| Nitrospina sp. SCGC AAA288-L16 | GCA_000372225.1 | 2.08 | 136 | 39.47 | p_Nitrospinota;c_Nitrospinia;o_Nitrospinales;f_Nitrospinaceae;g_SCGCAAA288-L16;s_SCGCAAA288-L16 sp000372225 | 89.28 | 5.86 |
| Nitrospina sp. SP265 | GCA_002721515.1 | 1.64 | 51 | 36.86 | p_Nitrospinota;c_Nitrospinia;o_Nitrospinales;f_Nitrospinaceae;g_UBA8687;s_UBA8687 sp002721515 | 95.73 | 5.34 |
| Nitrospinae bacterium bin107 | GCA_002238965.1 | 4.37 | 60 | 59.46 | p_Nitrospinota_A;c_UBA8248;o_UBA8248;f_UBA8248;g_Bin107;s_Bin107 sp002238965 | 60.69 | 3.87 |
| Nitrospinae bacterium MAG_09705_ntspn_70 | GCA_013349585.1 | 2.02 | 120 | 42.63 | p_Nitrospinota;c_Nitrospinia;o_Nitrospinales;f_Nitrospinaceae;g_UBA8687;s_UBA8687 sp013349585 | 78.43 | 5.08 |
| Nitrospinae bacterium nARSLQbin3 | GCA_015234085.1 | 2.64 | 440 | 54.88 | p_Nitrospinota;c_UBA7883;o_UBA7883;f_;g_;s_ | 98.17 | 3.96 |
| Nitrospinae bacterium nNGHbin12 | GCA_015231995.1 | 3.23 | 358 | 54.84 | p_Nitrospinota;c_UBA7883;o_UBA7883;f_;g_;s_ | 99.39 | 4.88 |
| Nitrospinae bacterium NP1026 | GCA_002731985.1 | 1.58 | 114 | 46.05 | p_Nitrospinota;c_Nitrospinia;o_Nitrospinales;f_Nitrospinaceae;g_SCGCAAA288-L16;s_SCGCAAA288-L16 sp002731985 | 87.64 | 2.12 |
| Nitrospinae bacterium nPCRbin9 | GCA_015231815.1 | 2.65 | 410 | 37.05 | p_Nitrospinota;c_UBA7883;o_;f_;g_;s_ | 96.95 | 3.05 |
| Nitrospinae bacterium nWMHbin6 | GCA_015229165.1 | 2.31 | 346 | 63.95 | p_Nitrospinota;c_UBA7883;o_UBA7883;f_;g_;s_ | 95.12 | 4.19 |
| Nitrospira bacterium HGW-Nitrospira-1 | GCA_002839535.1 | 1.66 | 182 | 46.15 | p_Nitrospirota;c_Thermodesulfovibrionia;o_Thermodesulfovibrionales;f_UBA6898;g_GW-Nitrospira-1;s_GW-Nitrospira-1 sp002839535 | 84.69 | 0.50 |
| Nitrospira defluvii | GCA_000196815.1 | 4.32 | 1 | 59.03 | p_Nitrospirota;c_Nitrospiria;o_Nitrospirales;f_Nitrospiraceae;g_Nitrospira_A;s_Nitrospira_A defluvii_A | 93.50 | 2.19 |
| Nitrospira japonica NJ11 | GCF_900169565.1 | 4.08 | 1 | 58.96 | p_Nitrospirota;c_Nitrospiria;o_Nitrospirales;f_Nitrospiraceae;g_Nitrospira_C;s_Nitrospira_C japonica | 79.91 | 1.17 |

| Genome name | NCBI/IMG accession | Size, Mbp | Scaff, no. | GC, % | GTDB Taxonomy | Completeness, % | Contamination, % |
|---|---|---|---|---|---|---|---|
| Nitrospira lenta BS10 | GCF_900403705.1 | 3.76 | 22 | 57.88 | p_Nitrospirota;c_Nitrospiria;o_Nitrospirales;f_Nitrospiraceae;g_Nitrospira_D;s_Nitrospira_D lenta | 98.10 | 1.26 |
| Nitrospira moscoviensis NSP M-1 | GCF_001273775.1 | 4.59 | 1 | 61.99 | p_Nitrospirota;c_Nitrospiria;o_Nitrospirales;f_Nitrospiraceae;g_Nitrospira_E;s_Nitrospira_E moscoviensis | 61.21 | 1.72 |
| Nitrospira sp. Bin_34_1 | GCA_005239745.1 | 1.63 | 145 | 52.91 | p_Nitrospirota;c_Nitrospiria;o_SBBL01;f_SBBL01;g_SBBL01;s_SBBL01 sp005239745 | 73.76 | 1.03 |
| Nitrospira sp. Bin_6_1_1 | GCA_005239595.1 | 1.88 | 236 | 50.25 | p_Nitrospirota;c_Nitrospiria;o_SBBL01;f_SBBT01;g_SBBT01;s_SBBT01 sp005239595 | 90.84 | 1.79 |
| Nitrospira sp. bin75 | GCA_002238765.1 | 3.04 | 115 | 56.24 | p_Nitrospirota;c_Nitrospiria;o_Nitrospirales;f_UBA8639;g_Bin75;s_Bin75 sp002238765 | 92.43 | 0.44 |
| Nitrospira sp. CG24A | GCA_002869925.2 | 3.36 | 40 | 55.68 | p_Nitrospirota;c_Nitrospiria;o_Nitrospirales;f_Nitrospiraceae;g_Palsa-1315;s_Palsa-1315 sp002869925 | 96.89 | 2.94 |
| Nitrospira sp. CG24B | GCA_002869845.2 | 3.14 | 20 | 55.28 | p_Nitrospirota;c_Nitrospiria;o_Nitrospirales;f_Nitrospiraceae;g_Nitrospira_F;s_Nitrospira_F sp002869845 | 94.12 | 2.10 |
| Nitrospira sp. CG24C | GCA_002869885.2 | 2.99 | 22 | 56.12 | p_Nitrospirota;c_Nitrospiria;o_Nitrospirales;f_Nitrospiraceae;g_Palsa-1315;s_Palsa-1315 sp002869885 | 97.48 | 2.10 |
| Nitrospira sp. CG24D | GCA_002869855.2 | 3.41 | 87 | 57.84 | p_Nitrospirota;c_Nitrospiria;o_Nitrospirales;f_Nitrospiraceae;g_Nitrospira_D;s_Nitrospira_D sp002869855 | 93.70 | 0.05 |
| Nitrospira sp. CG24E | GCA_002869895.2 | 3.45 | 99 | 55.84 | p_Nitrospirota;c_Nitrospiria;o_Nitrospirales;f_Nitrospiraceae;g_Palsa-1315;s_Palsa-1315 sp002869895 | 96.69 | 2.63 |
| Nitrospira sp. ND1 | GCF_900170025.1 | 4.45 | 6 | 58.87 | p_Nitrospirota;c_Nitrospiria;o_Nitrospirales;f_Nitrospiraceae;g_Nitrospira_A;s_Nitrospira_A sp900170025 | 81.68 | 0.50 |
| Nitrospira sp. OLB3 | GCA_001567445.1 | 3.75 | 79 | 60.36 | p_Nitrospirota;c_Nitrospiria;o_Nitrospirales;f_Nitrospiraceae;g_Nitrospira_A;s_Nitrospira_A sp001567445 | 96.77 | 0.52 |
| Nitrospira sp. palsa_1310 | GCA_003135435.1 | 4.34 | 254 | 56.94 | p_Nitrospirota;c_Nitrospiria;o_Nitrospirales;f_Nitrospiraceae;g_Palsa-1315;s_Palsa-1315 sp003135435 | 78.85 | 1.59 |
| Nitrospira sp. RCA | GCA_005239465.1 | 3.30 | 85 | 56.83 | p_Nitrospirota;c_Nitrospiria;o_Nitrospirales;f_Nitrospiraceae;g_Nitrospira_F;s_Nitrospira_F sp005239465 | 73.74 | 0.85 |
| Nitrospira sp. RCB | GCA_005239475.1 | 3.56 | 289 | 57.07 | p_Nitrospirota;c_Nitrospiria;o_Nitrospirales;f_Nitrospiraceae;g_Palsa-1315;s_Palsa-1315 sp005239475 | 60.70 | 0.00 |
| Nitrospira sp. RSF1 | GCA_005116965.1 | 3.73 | 27 | 55.24 | p_Nitrospirota;c_Nitrospiria;o_Nitrospirales;f_Nitrospiraceae;g_Nitrospira_F;s_Nitrospira_F sp005116965 | 75.33 | 5.04 |
| Nitrospira sp. RSF12 | GCA_005116955.1 | 3.65 | 8 | 55.15 | p_Nitrospirota;c_Nitrospiria;o_Nitrospirales;f_Nitrospiraceae;g_Nitrospira_F;s_Nitrospira_F sp005116955 | 91.04 | 3.36 |
| Nitrospira sp. RSF3 | GCA_005116835.1 | 3.94 | 24 | 55.56 | p_Nitrospirota;c_Nitrospiria;o_Nitrospirales;f_Nitrospiraceae;g_Palsa-1315;s_Palsa-1315 sp005116835 | 88.72 | 0.84 |
| Nitrospira sp. RSF5 | GCA_005116895.1 | 3.31 | 35 | 55.18 | p_Nitrospirota;c_Nitrospiria;o_Nitrospirales;f_Nitrospiraceae;g_Nitrospira_F;s_Nitrospira_F sp005116895 | 58.40 | 1.26 |
| Nitrospira sp. RSF6 | GCA_005116885.1 | 3.19 | 49 | 55.43 | p_Nitrospirota;c_Nitrospiria;o_Nitrospirales;f_Nitrospiraceae;g_Palsa-1315;s_Palsa-1315 sp005116885 | 87.17 | 1.26 |
| Nitrospira sp. RSF9 | GCA_005116745.1 | 3.86 | 32 | 54.96 | p_Nitrospirota;c_Nitrospiria;o_Nitrospirales;f_Nitrospiraceae;g_Nitrospira_F;s_Nitrospira_F sp005116745 | 75.18 | 1.68 |
| Nitrospira sp. Ru_enrich_NS | GCA_002634385.1 | 4.31 | 77 | 55.93 | p_Nitrospirota;c_Thermodesulfovibrionia;o_Thermodesulfovibrionales;f_UBA9935;g_GCA-2634385;s_GCA-2634385 sp002634385 | 94.79 | 2.10 |
| Nitrospira sp. SG-bin1 | GCA_002083365.1 | 4.42 | 48 | 56.08 | p_Nitrospirota;c_Nitrospiria;o_Nitrospirales;f_Nitrospiraceae;g_Nitrospira_F;s_Nitrospira_F sp002083365 | 95.63 | 0.00 |
| Nitrospira sp. SG-bin2 | GCA_002083405.1 | 3.66 | 63 | 56.77 | p_Nitrospirota;c_Nitrospiria;o_Nitrospirales;f_Nitrospiraceae;g_Nitrospira_F;s_Nitrospira_F sp002083405 | 78.88 | 3.85 |
| Nitrospira sp. ST-bin4 | GCA_002083565.1 | 2.94 | 117 | 57.02 | p_Nitrospirota;c_Nitrospiria;o_Nitrospirales;f_Nitrospiraceae;g_Nitrospira_F;s_Nitrospira_F sp002083565 | 65.43 | 0.00 |
| Nitrospira sp. ST-bin5 | GCA_002083555.1 | 4.01 | 29 | 58.02 | p_Nitrospirota;c_Nitrospiria;o_Nitrospirales;f_Nitrospiraceae;g_Nitrospira_D;s_Nitrospira_D sp002083555 | 57.14 | 0.00 |
| Nitrospira sp. UBA2082 | GCA_002331335.1 | 3.56 | 37 | 57.89 | p_Nitrospirota;c_Nitrospiria;o_Nitrospirales;f_Nitrospiraceae;g_Nitrospira_F;s_Nitrospira_F sp002331335 | 97.48 | 2.10 |
| Nitrospira sp. UBA5698 | GCA_002420115.1 | 4.38 | 73 | 55.06 | p_Nitrospirota;c_Nitrospiria;o_Nitrospirales;f_Nitrospiraceae;g_Nitrospira_F;s_Nitrospira_F sp002420115 | 83.19 | 5.04 |
| Nitrospira sp. UBA5699 | GCA_002420105.1 | 3.44 | 124 | 62.41 | p_Nitrospirota;c_Nitrospiria;o_Nitrospirales;f_Nitrospiraceae;g_Nitrospira_C;s_Nitrospira_C sp002420105 | 96.64 | 2.10 |
| Nitrospira sp. UW-LDO-01 | GCA_002254365.1 | 3.91 | 230 | 54.93 | p_Nitrospirota;c_Nitrospiria;o_Nitrospirales;f_Nitrospiraceae;g_Nitrospira_F;s_Nitrospira_F sp002254365 | 58.62 | 0.00 |
| Nitrospiraceae bacterium fen_1308 | GCA_003170655.1 | 2.83 | 254 | 49.38 | p_Nitrospirota;c_Thermodesulfovibrionia;o_Thermodesulfovibrionales;f_UBA9935;g_Fen-1308;s_Fen-1308 sp003170655 | 88.29 | 2.15 |
| Nitrospiraceae bacterium SURF_11 | GCA_003599505.1 | 2.64 | 87 | 47.98 | p_Nitrospirota;c_Thermodesulfovibrionia;o_UBA6902;f_UBA6902;g_SURF-11;s_SURF-11 sp003599505 | 76.53 | 3.36 |
| Nitrospiraceae bacterium SURF_45 | GCA_003599275.1 | 3.05 | 74 | 47.56 | p_Nitrospirota;c_Thermodesulfovibrionia;o_UBA6902;f_UBA6902;g_SURF-45;s_SURF-45 sp003599275 | 88.69 | 1.10 |
| Nitrospiraceae bacterium UBA2194 | GCA_002328665.1 | 2.20 | 132 | 47.95 | p_Nitrospirota;c_Thermodesulfovibrionia;o_Thermodesulfovibrionales;f_SM23-35;g_UBA2194;s_UBA2194 sp002328665 | 90.48 | 0.00 |
| Nitrospiraceae bacterium UBA2600 | GCA_002339325.1 | 1.90 | 53 | 36.78 | p_Nitrospirota;c_Thermodesulfovibrionia;o_Thermodesulfovibrionales;f_Thermodesulfovibrionaceae;g_Thermodesulfovibrio;s_sp002339325 | 90.55 | 1.68 |
| Nitrospiraceae bacterium UBA665 | GCA_002299835.1 | 2.02 | 173 | 40.35 | p_Nitrospirota;c_Thermodesulfovibrionia;o_Thermodesulfovibrionales;f_UBA9935;g_UBA665;s_UBA665 sp002299835 | 86.97 | 3.78 |
| Nitrospiraceae bacterium UBA6902 | GCA_002451135.1 | 3.05 | 160 | 46.5 | p_Nitrospirota;c_Thermodesulfovibrionia;o_UBA6902;f_UBA6902;g_UBA6902;s_UBA6902 sp002451135 | 73.79 | 1.72 |
| Nitrospiraceae bacterium UBA9159 | GCA_003453735.1 | 4.66 | 540 | 48.26 | p_Nitrospirota;c_Thermodesulfovibrionia;o_Thermodesulfovibrionales;f_UBA9159;g_UBA9159;s_UBA9159 sp003453735 | 95.65 | 1.97 |
| Nitrospiraceae bacterium UBA9217 | GCA_003454665.1 | 3.01 | 388 | 54.65 | p_Nitrospirota;c_UBA9217;o_UBA9217;f_UBA9217;g_UBA9217;s_UBA9217 sp003454665 | 73.13 | 0.97 |
| Nitrospirae bacterium Baikal-G1 | GCA_002737345.1 | 1.67 | 38 | 57.92 | p_Nitrospirota;c_Nitrospiria;o_Nitrospirales;f_Nitrospiraceae;g_Palsa-1315;s_Palsa-1315 sp002737345 | 95.38 | 0.84 |
| Nitrospirae bacterium BMS3Abin08 | GCA_002897935.1 | 2.61 | 123 | 47.88 | p_Nitrospirota;c_Thermodesulfovibrionia;o_Thermodesulfovibrionales;f_JdFR-85;g_BMS3Abin08;s_BMS3Abin08 sp002897935 | 96.77 | 1.75 |
| Nitrospirae bacterium BMS3Bbin05 | GCA_002897855.1 | 2.46 | 89 | 44.67 | p_Nitrospirota;c_Thermodesulfovibrionia;o_Thermodesulfovibrionales;f_BMS3Bbin05;g_BMS3Bbin05;s_BMS3Bbin05 sp002897855 | 94.65 | 0.33 |
| Nitrospirae bacterium BMS3Bbin08 | GCA_002897775.1 | 2.71 | 67 | 45.24 | p_Nitrospirota;c_Thermodesulfovibrionia;o_UBA6902;f_BMS3Bbin08;g_BMS3Bbin08;s_BMS3Bbin08 sp002897775 | 56.19 | 0.00 |
| Nitrospirae bacterium CG02_land_8_20_14_3_00_41_53 | GCA_002780895.1 | 2.01 | 159 | 40.63 | p_Nitrospirota;c_Thermodesulfovibrionia;o_Thermodesulfovibrionales;f_SM23-35;g_0-14-3-00-41-53;s_0-14-3-00-41-53 sp002780895 | 78.36 | 1.75 |
| Nitrospirae bacterium CG1_02_44_142 | GCA_001871685.1 | 1.79 | 138 | 43.94 | p_Nitrospirota;c_Thermodesulfovibrionia;o_Thermodesulfovibrionales;f_UBA1546;g_UBA1546;s_UBA1546 sp001871685 | 95.67 | 1.29 |
| Nitrospirae bacterium DC0425bin1 | GCA_002753685.1 | 4.02 | 107 | 49.04 | p_Nitrospirota;c_Thermodesulfovibrionia;o_Thermodesulfovibrionales;f_Magnetobacteriaceae;g_Magnetobacterium;s_Magnetobacterium sp002753685 | 97.06 | 0.91 |
| Nitrospirae bacterium DP16D_bin.35 | GCA_004321915.1 | 3.63 | 145 | 48.51 | p_Nitrospirota;c_Thermodesulfovibrionia;o_Thermodesulfovibrionales;f_UBA9935;g_MYbin3;s_MYbin3 sp004321915 | 53.73 | 0.05 |
| Nitrospirae bacterium DP16D_bin.48 | GCA_004321925.1 | 3.23 | 120 | 54.48 | p_Nitrospirota;c_Thermodesulfovibrionia;o_Thermodesulfovibrionales;f_UBA6898;g_PALSA-1316;s_PALSA-1316 sp004321925 | 82.29 | 0.81 |
| Nitrospirae bacterium FW300_bin.22 | GCA_004298625.1 | 2.34 | 416 | 45.48 | p_Nitrospirota;c_Thermodesulfovibrionia;o_Thermodesulfovibrionales;f_UBA1546;g_SCSY01;s_SCSY01 sp004298625 | 81.55 | 0.25 |
| Nitrospirae bacterium FW300_bin.52 | GCA_004298335.1 | 2.66 | 250 | 63.01 | p_Nitrospirota;c_Nitrospiria;o_Nitrospirales;f_NS-4;g_SCTG01;s_SCTG01 sp004298335 | 97.42 | 1.29 |
| Nitrospirae bacterium Glo_13 | GCA_003354025.1 | 4.17 | 642 | 42.23 | p_Nitrospirota;c_Thermodesulfovibrionia;o_UBA6902;f_UBA6902;g_Glo-13;s_Glo-13 sp003354025 | 96.57 | 2.94 |
| Nitrospirae bacterium GW715_bin.13 | GCA_004297235.1 | 4.17 | 613 | 57.81 | p_Nitrospirota;c_Nitrospiria;o_SBBL01;f_SCUR01;g_Manganitrophus;s_Manganitrophus sp004297235 | 73.57 | 1.94 |
| Nitrospirae bacterium GW928_bin.16 | GCA_004296885.1 | 3.26 | 23 | 61.97 | p_Nitrospirota;c_Nitrospiria;o_Nitrospirales;f_Nitrospiraceae;g_SYGV01;s_SYGV01 sp004296885 | 61.77 | 1.47 |
| Nitrospirae bacterium GW928_bin.19 | GCA_004296865.1 | 2.71 | 613 | 61.63 | p_Nitrospirota;c_Nitrospiria;o_Nitrospirales;f_Nitrospiraceae;g_SCVQ01;s_SCVQ01 sp004296865 | 97.78 | 1.68 |
| Nitrospirae bacterium GWB2_47_37 | GCA_001803635.1 | 2.47 | 81 | 46.39 | p_Nitrospirota;c_Thermodesulfovibrionia;o_Thermodesulfovibrionales;f_UBA9935;g_GWB2-47-37;s_GWB2-47-37 sp001803635 | 70.28 | 1.68 |
| Nitrospirae bacterium GWC2_56_14 | GCA_001803705.1 | 3.16 | 259 | 56.44 | p_Nitrospirota;c_UBA9217;o_UBA9217;f_UBA9217;g_GWC2-56-14;s_GWC2-56-14 sp001803705 | 93.82 | 0.00 |
| Nitrospirae bacterium GWC2_57_13 | GCA_001805055.1 | 3.32 | 105 | 57.15 | p_Nitrospirota;c_UBA9217;o_UBA9217;f_UBA9217;g_GWC2-57-13;s_GWC2-57-13 sp001805055 | 99.11 | 1.68 |
| Nitrospirae bacterium GWF2_44_13 | GCA_001805105.1 | 1.99 | 18 | 43.69 | p_Nitrospirota;c_Thermodesulfovibrionia;o_Thermodesulfovibrionales;f_UBA1546;g_UBA1546;s_UBA1546 sp001805105 | 99.68 | 1.61 |
| Nitrospirae bacterium HCH-1 | GCA_001541255.1 | 3.59 | 152 | 45.37 | p_Nitrospirota;c_Thermodesulfovibrionia;o_Thermodesulfovibrionales;f_Magnetobacteriaceae;g_HCH-1;s_HCH-1 sp001541255 | 91.07 | 2.68 |

| Genome name | NCBI/IMG accession | Size, Mbp | Scaff, no. | GC, % | GTDB Taxonomy | Completeness, % | Contamination, % |
|---|---|---|---|---|---|---|---|
| Nitrospirae bacterium HCHbin1 | GCA_002753435.1 | 3.69 | 94 | 45.24 | p_Nitrospirota;c_Thermodesulfovibrionia;o_Thermodesulfovibrionales;f_Magnetobacteriaceae;g_HCH-1;s_HCH-1 sp001541255 | 95.66 | 0.00 |
| Nitrospirae bacterium J017 | GCA_003695915.1 | 3.71 | 135 | 54.77 | p_Nitrospirota;c_Nitrospiria;o_Nitrospirales;f_UBA8639;g_J017;s_J017 sp003695915 | 78.06 | 5.22 |
| Nitrospirae bacterium J031 | GCA_003696975.1 | 2.77 | 273 | 53.77 | p_Nitrospirota;c_Nitrospiria;o_Nitrospirales;f_UBA8639;g_J031;s_J031 sp003696975 | 58.65 | 0.00 |
| Nitrospirae bacterium JdFR-81 | GCA_002011735.1 | 2.05 | 95 | 48.33 | p_Nitrospirota;c_Thermodesulfovibrionia;o_UBA6902;f_JdFR-81;g_JdFR-81;s_JdFR-81 sp002011735 | 61.47 | 1.68 |
| Nitrospirae bacterium JdFR-85 | GCA_002011745.1 | 2.33 | 47 | 41.54 | p_Nitrospirota;c_Thermodesulfovibrionia;o_Thermodesulfovibrionales;f_JdFR-85;g_JdFR-85;s_JdFR-85 sp002011745 | 68.97 | 0.00 |
| Nitrospirae bacterium JdFR-86 | GCA_002011815.1 | 2.10 | 25 | 45.26 | p_Nitrospirota;c_Thermodesulfovibrionia;o_Thermodesulfovibrionales;f_JdFR-86;g_JdFR-86;s_JdFR-86 sp002011815 | 63.44 | 1.08 |
| Nitrospirae bacterium JdFR-88 | GCA_002011795.1 | 1.86 | 22 | 62.9 | p_Nitrospirota;c_Thermodesulfovibrionia;o_Thermodesulfovibrionales;f_JdFR-88;g_JdFR-88;s_JdFR-88 sp002011795 | 84.62 | 0.16 |
| Nitrospirae bacterium MAG_10313_ntr_31 | GCA_013349595.1 | 1.93 | 344 | 35.33 | p_Nitrospirota;c_Thermodesulfovibrionia;o_Thermodesulfovibrionales;f_DUZI01;g_DUZI01;s_DUZI01 sp013349595 | 88.39 | 0.32 |
| Nitrospirae bacterium MYbin2 | GCA_002753455.1 | 3.07 | 89 | 48.79 | p_Nitrospirota;c_Thermodesulfovibrionia;o_Thermodesulfovibrionales;f_Magnetobacteriaceae;g_Magnetobacterium;s_Magnetobacterium casensis | 85.64 | 5.45 |
| Nitrospirae bacterium MYbin3 | GCA_002753335.1 | 2.93 | 66 | 44.36 | p_Nitrospirota;c_Thermodesulfovibrionia;o_Thermodesulfovibrionales;f_UBA9935;g_MYbin3;s_MYbin3 sp002753335 | 81.86 | 1.68 |
| Nitrospirae bacterium MYbin6 | GCA_002753305.1 | 3.60 | 218 | 47.78 | p_Nitrospirota;c_Thermodesulfovibrionia;o_Thermodesulfovibrionales;f_Magnetobacteriaceae;g_HCH-1;s_HCH-1 sp002753305 | 83.77 | 1.79 |
| Nitrospirae bacterium MYbinv3 | GCA_002753395.1 | 3.71 | 175 | 44.44 | p_Nitrospirota;c_Thermodesulfovibrionia;o_Thermodesulfovibrionales;f_Magnetobacteriaceae;g_Magnetobacterium;s_Magnetobacterium sp002753395 | 59.65 | 0.00 |
| Nitrospirae bacterium nDC0425bin1 | GCA_015233985.1 | 4.04 | 178 | 49.09 | p_Nitrospirota;c_Thermodesulfovibrionia;o_Thermodesulfovibrionales;f_Magnetobacteriaceae;g_Magnetobacterium;s_Magnetobacterium sp002753685 | 67.50 | 1.08 |
| Nitrospirae bacterium nDJH13bin1 | GCA_015233725.1 | 3.32 | 168 | 49.07 | p_Nitrospirota;c_Thermodesulfovibrionia;o_Thermodesulfovibrionales;f_Magnetobacteriaceae;g_Magnetobacterium;s_ | 99.50 | 0.50 |
| Nitrospirae bacterium nDJH13bin15 | GCA_015233685.1 | 2.03 | 121 | 48.07 | p_Nitrospirota;c_Thermodesulfovibrionia;o_Thermodesulfovibrionales;f_Magnetobacteriaceae;g_HCH-1;s_ | 94.12 | 2.10 |
| Nitrospirae bacterium nDJH13bin21 | GCA_015233655.1 | 3.29 | 262 | 47.98 | p_Nitrospirota;c_Thermodesulfovibrionia;o_Thermodesulfovibrionales;f_UBA9935;g_GCA-2634385;s_ | 98.18 | 0.91 |
| Nitrospirae bacterium nDJH13bin3 | GCA_015233615.1 | 4.01 | 220 | 47.81 | p_Nitrospirota;c_Thermodesulfovibrionia;o_Thermodesulfovibrionales;f_Magnetobacteriaceae;g_;s_ | 98.51 | 0.50 |
| Nitrospirae bacterium nDJH14bin5 | GCA_015233555.1 | 3.59 | 69 | 41.5 | p_Nitrospirota;c_Thermodesulfovibrionia;o_Thermodesulfovibrionales;f_Magnetobacteriaceae;g_;s_ | 92.24 | 1.68 |
| Nitrospirae bacterium nDJH14bin7 | GCA_015233485.1 | 4.24 | 565 | 44.94 | p_Nitrospirota;c_Thermodesulfovibrionia;o_Thermodesulfovibrionales;f_Magnetobacteriaceae;g_HCH-1;s_ | 96.47 | 3.36 |
| Nitrospirae bacterium nDJH14bin9 | GCA_015233465.1 | 3.13 | 75 | 35.21 | p_Nitrospirota;c_Thermodesulfovibrionia;o_Thermodesulfovibrionales;f_;g_;s_ | 98.71 | 1.36 |
| Nitrospirae bacterium nDJH15bin2 | GCA_015233455.1 | 3.58 | 245 | 49.18 | p_Nitrospirota;c_Thermodesulfovibrionia;o_Thermodesulfovibrionales;f_Magnetobacteriaceae;g_Magnetobacterium;s_ | 76.25 | 0.50 |
| Nitrospirae bacterium nDJH15bin8 | GCA_015233365.1 | 2.18 | 62 | 48.65 | p_Nitrospirota;c_Thermodesulfovibrionia;o_Thermodesulfovibrionales;f_UBA9935;g_GCA-2634385;s_ | 96.00 | 0.00 |
| Nitrospirae bacterium nDJH5bin4 | GCA_015233265.1 | 2.54 | 344 | 47.41 | p_Nitrospirota;c_Thermodesulfovibrionia;o_Thermodesulfovibrionales;f_Magnetobacteriaceae;g_HCH-1;s_ | 90.84 | 0.00 |
| Nitrospirae bacterium nDJH6bin1 | GCA_015233245.1 | 3.47 | 318 | 41.13 | p_Nitrospirota;c_Thermodesulfovibrionia;o_Thermodesulfovibrionales;f_Magnetobacteriaceae;g_;s_ | 98.62 | 2.52 |
| Nitrospirae bacterium nDJH8bin13 | GCA_015233045.1 | 2.61 | 190 | 47.66 | p_Nitrospirota;c_Thermodesulfovibrionia;o_Thermodesulfovibrionales;f_Magnetobacteriaceae;g_HCH-1;s_ | 93.38 | 0.75 |
| Nitrospirae bacterium nDJH8bin6 | GCA_015233015.1 | 3.78 | 149 | 46.6 | p_Nitrospirota;c_Thermodesulfovibrionia;o_Thermodesulfovibrionales;f_Magnetobacteriaceae;g_HCH-1;s_ | 99.58 | 0.84 |
| Nitrospirae bacterium nDJH8bin7 | GCA_015232945.1 | 1.91 | 354 | 35.14 | p_Nitrospirota;c_Thermodesulfovibrionia;o_Thermodesulfovibrionales;f_;g_;s_ | 88.80 | 0.56 |
| Nitrospirae bacterium nDJH8bin8 | GCA_015232995.1 | 4.19 | 111 | 41.49 | p_Nitrospirota;c_Thermodesulfovibrionia;o_Thermodesulfovibrionales;f_Magnetobacteriaceae;g_;s_ | 51.04 | 0.00 |
| Nitrospirae bacterium nHCHbin2 | GCA_015232615.1 | 3.70 | 99 | 45.24 | p_Nitrospirota;c_Thermodesulfovibrionia;o_Thermodesulfovibrionales;f_Magnetobacteriaceae;g_HCH-1;s_HCH-1 sp001541255 | 88.82 | 1.20 |
| Nitrospirae bacterium nMYbin1 | GCA_015232195.1 | 2.91 | 48 | 44.36 | p_Nitrospirota;c_Thermodesulfovibrionia;o_Thermodesulfovibrionales;f_UBA9935;g_MYbin3;s_MYbin3 sp002753335 | 84.03 | 4.62 |
| Nitrospirae bacterium nMYbin2 | GCA_015232185.1 | 3.49 | 127 | 49 | p_Nitrospirota;c_Thermodesulfovibrionia;o_Thermodesulfovibrionales;f_Magnetobacteriaceae;g_Magnetobacterium;s_Magnetobacterium casensis | 52.67 | 0.42 |
| Nitrospirae bacterium nMYbin3 | GCA_015232165.1 | 1.92 | 278 | 49.89 | p_Nitrospirota;c_Thermodesulfovibrionia;o_Thermodesulfovibrionales;f_Magnetobacteriaceae;g_Magnetobacterium;s_ | 80.97 | 0.65 |
| Nitrospirae bacterium nMYbin4 | GCA_015232145.1 | 3.93 | 213 | 44.04 | p_Nitrospirota;c_Thermodesulfovibrionia;o_Thermodesulfovibrionales;f_Magnetobacteriaceae;g_Magnetobacterium;s_Magnetobacterium sp002753395 | 90.08 | 2.73 |
| Nitrospirae bacterium nMYbin6 | GCA_015232115.1 | 3.40 | 171 | 47.77 | p_Nitrospirota;c_Thermodesulfovibrionia;o_Thermodesulfovibrionales;f_Magnetobacteriaceae;g_HCH-1;s_HCH-1 sp002753305 | 70.68 | 0.75 |
| Nitrospirae bacterium NS_7 | GCA_005877775.1 | 3.06 | 309 | 58.5 | p_Nitrospirota;c_Nitrospiria;o_Nitrospirales;f_Nitrospiraceae;g_NS-7;s_NS-7 sp005877775 | 97.01 | 0.00 |
| Nitrospirae bacterium NS_8 | GCA_005877565.1 | 2.23 | 265 | 62.11 | p_Nitrospirota;c_Nitrospiria;o_Nitrospirales;f_NS-4;g_NS-12;s_NS-12 sp005877565 | 93.01 | 1.08 |
| Nitrospirae bacterium NS_9 | GCA_005887945.1 | 3.54 | 366 | 56.73 | p_Nitrospirota;c_Nitrospiria;o_Nitrospirales;f_Nitrospiraceae;g_Palsa-1315;s_Palsa-1315 sp005887945 | 68.82 | 0.18 |
| Nitrospirae bacterium RBG_13_39_12 | GCA_001805125.1 | 2.93 | 176 | 39.49 | p_Nitrospirota;c_Thermodesulfovibrionia;o_Thermodesulfovibrionales;f_SM23-35;g_RBG-13-39-12;s_RBG-13-39-12 sp001805125 | 80.66 | 2.94 |
| Nitrospirae bacterium RIFCSPLOWO2_02_FULL_62_14 | GCA_001805245.1 | 2.66 | 76 | 61.66 | p_Nitrospirota;c_Nitrospiria;o_Nitrospirales;f_Nitrospiraceae;g_2-02-FULL-62-14;s_2-02-FULL-62-14 sp001805245 | 100.00 | 0.91 |
| Nitrospirae bacterium SZUA-452 | GCA_003252075.1 | 2.86 | 146 | 52.15 | p_Nitrospirota;c_Thermodesulfovibrionia;o_Thermodesulfovibrionales;f_UBA6898;g_UBA6898;s_UBA6898 sp003252075 | 73.59 | 2.21 |
| Oligoflexales bacterium nHA4bin4 | GCA_015232735.1 | 4.29 | 942 | 43.67 | p_Bdellovibrionota;c_Oligoflexia;o_Oligoflexales;f_Oligoflexaceae;g_;s_ | 100.00 | 0.00 |
| Oligoflexia bacterium nKLKbin12 | GCA_015232355.1 | 5.63 | 96 | 32.16 | p_Bdellovibrionota;c_Bacteriovoracia;o_Bacteriovoracales;f_;g_;s_ | 98.88 | 1.12 |
| Oligoflexia bacterium nKLKbin5 | GCA_015232255.1 | 4.59 | 212 | 41.45 | p_Bdellovibrionota;c_Bacteriovoracia;o_Bacteriovoracales;f_;g_;s_ | 100.00 | 1.69 |
| Oligoflexia bacterium nN3bin16 | GCA_015232015.1 | 4.91 | 413 | 31.52 | p_Bdellovibrionota;c_Bacteriovoracia;o_Bacteriovoracales;f_;g_;s_ | 100.00 | 1.14 |
| Oligoflexia bacterium nN3bin31 | GCA_015231955.1 | 5.61 | 114 | 40.9 | p_Bdellovibrionota;c_Bacteriovoracia;o_Bacteriovoracales;f_;g_;s_ | 98.32 | 0.84 |
| Oligoflexia bacterium nW3bin19 | GCA_015231325.1 | 4.37 | 470 | 38.41 | p_Bdellovibrionota;c_Bacteriovoracia;o_Bacteriovoracales;f_Bacteriovoracaceae;g_;s_ | 98.78 | 0.00 |
| Omnitrophica bacterium GWA2_52_8 | GCA_001804025.1 | 2.20 | 146 | 51.7 | p_Omnitrophota;c_Omnitrophia;o_Omnitrophales;f_GWA2-52-8;g_GWA2-52-8;s_GWA2-52-8 sp001804025 | 85.56 | 0.45 |
| Omnitrophica bacterium SCGC AG-290-C17 | 3300015153 | 1.71 | 171 | 48.60 | p_Omnitrophota;c_Omnitrophia;o_Omnitrophales;f_GWA2-52-8;g_;s_ | 62.84 | 0.00 |
| Omnitrophica WOR_2 bacterium GWA2_45_18 | GCA_001804395.1 | 2.34 | 24 | 46.36 | p_Omnitrophota;c_Koll11;o_UBA10015;f_Kpj58rc;g_UBA10174;s_UBA10174 sp003528115 | 62.54 | 1.08 |
| Omnitrophica WOR_2 bacterium GWC2_45_7 | GCA_001805405.1 | 1.34 | 98 | 45.06 | p_Omnitrophota;c_Koll11;o_UBA10015;f_Kpj58rc;g_UBA10174;s_UBA10174 sp003528115 | 72.63 | 1.06 |
| Pelobacter propionicus DSM 2379 | GCF_000015045.1 | 4.24 | 3 | 58.48 | p_Desulfobacterota_F;c_Desulfuromonadia;o_Geobacterales;f_Pseudopelobacteraceae;g_Pseudopelobacter;s_Pseudopelobacter propionicus | 94.03 | 2.73 |
| Pelobacter seleniigenes DSM 18267 | GCA_000711225.1 | 5.09 | 6 | 54.19 | p_Desulfobacterota_F;c_Desulfuromonadia;o_Desulfuromonadales;f_Geopsychrobacteraceae;g_Seleniibacterium;s_Seleniibacterium seleniigenes | 96.48 | 3.18 |
| Pelobacter sp. UBA5610 | GCA_002424645.1 | 2.70 | 168 | 53.45 | p_Desulfobacterota_F;c_Desulfuromonadia;o_Desulfuromonadales;f_Syntrophotaleaceae;g_Syntrophotalea;s_Syntrophotalea sp002424645 | 63.39 | 1.22 |
| Pelobacter sp. UBA6812 | GCA_002452265.1 | 3.02 | 90 | 52.83 | p_Desulfobacterota_F;c_Desulfuromonadia;o_Desulfuromonadales;f_Syntrophotaleaceae;g_Syntrophotalea;s_Syntrophotalea sp002452265 | 91.27 | 2.38 |
| Phaeospirillum molischianum DSM 120 | GCF_000294655.1 | 3.81 | 61 | 61.51 | p_Proteobacteria;c_Alphaproteobacteria;o_Rhodospirillales;f_Magnetospirillaceae;g_Phaeospirillum;s_Phaeospirillum molischianum | 58.23 | 0.81 |
| Phycisphaera mikurensis NBRC 102666 | GCA_000284115.1 | 3.88 | 2 | 73.22 | p_Planctomycetota;c_Phycisphaerae;o_Phycisphaerales;f_Phycisphaeraceae;g_Phycisphaera;s_Phycisphaera mikurensis | 95.85 | 0.91 |
| Phycisphaerae bacterium ST-NAGAB-D1 | GCA_002007645.1 | 4.25 | 1 | 51.98 | p_Planctomycetota;c_Phycisphaerae;o_Sedimentisphaerales;f_Anaerohalosphaeraceae;g_Anaerohalosphaera;s_Anaerohalosphaera lusitana | 96.00 | 1.33 |
| Phycisphaerales bacterium UBA1845 | GCA_002338715.1 | 5.25 | 148 | 57 | p_Planctomycetota;c_Phycisphaerae;o_UBA1845;f_UBA1845;g_UBA1845;s_UBA1845 sp002338715 | 85.99 | 2.25 |
| Pirellula staleyi DSM 6068 | GCA_000025185.1 | 6.20 | 1 | 57.46 | p_Planctomycetota;c_Planctomycetes;o_Pirellulales;f_Pirellulaceae;g_Pirellula;s_Pirellula staleyi | 54.51 | 0.00 |

| Genome name | NCBI/IMG accession | Size, Mbp | Scaff , no. | GC, % | GTDB Taxonomy | Complete ness, % | Contamin ation, % |
|---|---|---|---|---|---|---|---|
| Planctomycetes bacterium MAG_11118_pl_115 | GCA_013349615.1 | 3.77 | 157 | 48.98 | p_Planctomycetota;c_Phycisphaerae;o_Sedimentisphaerales;f_SG8-4;g_CS2-K091;s_CS2-K091 sp013349615 | 56.21 | 0.00 |
| Planctomycetes bacterium MAG_17991_pl_60 | GCA_013349525.1 | 1.31 | 145 | 49.53 | p_Planctomycetota;c_Phycisphaerae;o_Sedimentisphaerales;f_Anaerohalophaeraceae;g_QNBT01;s_QNBT01 sp013349435 | 77.94 | 1.82 |
| Planctomycetes bacterium MAG_18080_pl_157 | GCA_013349435.1 | 3.14 | 139 | 48.44 | p_Planctomycetota;c_Phycisphaerae;o_Sedimentisphaerales;f_Anaerohalophaeraceae;g_QNBT01;s_QNBT01 sp013349435 | 73.86 | 1.33 |
| Planctomycetes bacterium nTSbin13 | GCA_015231495.1 | 3.00 | 642 | 56.47 | p_Planctomycetota;c_UBA11346;o_;f_;g_;s_ | 88.29 | 2.15 |
| Planctomycetes bacterium nwagbin3 | GCA_015231345.1 | 4.14 | 588 | 41.73 | p_Planctomycetota;c_UBA11346;o_;f_;g_;s_ | 75.50 | 1.29 |
| Planctomycetes bacterium SCGC JGI090-P21 | 2264265205 | 1.23 | 242 | 49.20 | p_Planctomycetota;c_Phycisphaerae;o_UBA1845;f_PWPN01;g_;s_ | 38.87 | 2.19 |
| Planctomycetes bacterium SM23_25 | GCA_001303605.1 | 3.77 | 418 | 66.68 | p_Planctomycetota;c_Phycisphaerae;o_FEN-1346;f_FEN-1346;g_;s_ | 60.62 | 0.32 |
| Pseudomonas aeruginosa JCM 5962 | GCA_000615485.1 | 6.07 | 1198 | 66.28 | p_Proteobacteria;c_Gammaproteobacteria;o_Pseudomonadales;f_Pseudomonadaceae;g_Pseudomonas;s_Pseudomonas aeruginosa | 99.04 | 0.00 |
| Rhodospirillaceae bacterium MAG_01419_mvb_30 | 3300001419 | 2.81 | 477 | 55.72 | p_Proteobacteria;c_Alphaproteobacteria;o_Rhodospirillales;f_Magnetovibrionaceae;g_Magnetovibrio;s_ | 94.58 | 4.10 |
| Rhodospirillaceae bacterium MAG_04806_tlms_2 | GCA_013349735.1 | 2.09 | 309 | 57.51 | p_Proteobacteria;c_Alphaproteobacteria;o_Rhodospirillales;f_Magnetospirillaceae;g_Telmatospirillum;s_Telmatospirillum sp013349735 | 94.56 | 1.08 |
| Rhodospirillaceae bacterium MAG_05422_2-02_14 | GCA_013349695.1 | 2.28 | 255 | 61.09 | p_Proteobacteria;c_Alphaproteobacteria;o_Rhodospirillales;f_2-02-FULL-58-16;g_GCA-2686765;s_GCA-2686765 sp013349695 | 71.32 | 1.79 |
| Rhodospirillaceae bacterium MAG_05596_2-02_51 | GCA_013349685.1 | 1.83 | 329 | 61.19 | p_Proteobacteria;c_Alphaproteobacteria;o_Rhodospirillales;f_2-02-FULL-58-16;g_GCA-2686765;s_GCA-2686765 sp013349695 | 80.77 | 2.10 |
| Rhodospirillaceae bacterium MAG_06104_tlms_034 | GCA_013349665.1 | 3.19 | 353 | 64.25 | p_Proteobacteria;c_Alphaproteobacteria;o_Rhodospirillales;f_Magnetospirillaceae;g_Telmatospirillum;s_Telmatospirillum sp013349665 | 99.41 | 0.59 |
| Rhodospirillaceae bacterium MAG_22225_2-02_112 | GCA_013349335.1 | 2.55 | 147 | 61.01 | p_Proteobacteria;c_Alphaproteobacteria;o_Rhodospirillales;f_2-02-FULL-58-16;g_GCA-2686765;s_GCA-2686765 sp013349695 | 54.48 | 0.00 |
| Rhodospirillum rubrum ATCC 11170 | GCF_000013085.1 | 4.41 | 2 | 65.38 | p_Proteobacteria;c_Alphaproteobacteria;o_Rhodospirillales;f_Rhodospirillaceae;g_Rhodospirillum;s_Rhodospirillum rubrum | 98.06 | 1.29 |
| SAR324 cluster bacterium nKLKbin6 | GCA_015232315.1 | 5.25 | 88 | 45.21 | p_SAR324;c_SAR324;o_SAR324;f_;g_;s_ | 100.00 | 0.65 |
| SAR324 cluster bacterium nPCRbin10 | GCA_015231885.1 | 5.13 | 241 | 42.00 | p_SAR324;c_SAR324;o_SAR324;f_GCA-2753255;g_GCA-2753255;s_GCA-2753255 sp002753255 | 95.97 | 2.15 |
| SAR324 cluster bacterium nPCRbin7 | GCA_015231825.1 | 5.80 | 81 | 44.77 | p_SAR324;c_SAR324;o_SAR324;f_GCA-2753255;g_;s_ | 96.82 | 3.92 |
| SAR324 cluster bacterium nTSbin10 | GCA_015231535.1 | 5.76 | 344 | 44.87 | p_SAR324;c_SAR324;o_SAR324;f_;g_;s_ | 100.00 | 0.00 |
| Sebaldella termitidis ATCC 33386 | GCA_000024405.1 | 4.49 | 3 | 33.38 | p_Fusobacteriota;c_Fusobacteriia;o_Fusobacteriales;f_Leptotrichiaceae;g_Sebaldella;s_Sebaldella termitidis | 64.20 | 0.00 |
| Smithella sp. D17 | GCA_000753945.1 | 1.64 | 271 | 43.19 | p_Desulfobacterota;c_Syntrophia;o_Syntrophales;f_Smithellaceae;g_Smithella;s_Smithella sp000753945 | 97.42 | 3.21 |
| Smithella sp. M82 | GCA_001683985.1 | 2.75 | 236 | 42.83 | p_Desulfobacterota;c_Syntrophia;o_Syntrophales;f_Smithellaceae;g_Smithella;s_Smithella sp001683985 | 98.71 | 2.72 |
| Smithella sp. PtaU1.Bin162 | GCA_002068015.1 | 3.60 | 191 | 45.33 | p_Desulfobacterota;c_Syntrophia;o_Syntrophales;f_Smithellaceae;g_Fen-1166;s_Fen-1166 sp002068015 | 96.59 | 0.91 |
| Smithella sp. SCADC | GCA_000747625.2 | 3.19 | 244 | 43.97 | p_Desulfobacterota;c_Syntrophia;o_Syntrophales;f_Smithellaceae;g_Smithella;s_Smithella sp000747625 | 96.97 | 0.00 |
| Smithella sp. SDB_sulfate1 SDB_sulfate2 | GCA_007244375.1 | 3.10 | 58 | 41.44 | p_Desulfobacterota;c_Syntrophia;o_Syntrophales;f_Smithellaceae;g_Smithella;s_Smithella sp001412345 | 96.77 | 1.94 |
| Smithella sp. UBA2180 | GCA_002327415.1 | 1.75 | 189 | 42.54 | p_Desulfobacterota;c_Syntrophia;o_Syntrophales;f_Smithellaceae;g_Smithella;s_Smithella sp002327415 | 96.45 | 0.97 |
| Streptobacillus moniliformis DSM 12112 | GCA_000024565.1 | 1.67 | 2 | 26.28 | p_Fusobacteriota;c_Fusobacteriia;o_Fusobacteriales;f_Leptotrichiaceae;g_Streptobacillus;s_Streptobacillus moniliformis | 60.01 | 1.16 |
| Sulfurivirga caldicuralii DSM 17737 | GCF_900141795.1 | 1.67 | 7 | 59.19 | p_Proteobacteria;c_Gammaproteobacteria;o_Thiomicrospirales;f_Thiomicrospiraceae;g_Sulfurivirga;s_Sulfurivirga caldicuralii | 99.50 | 0.75 |
| Syntrophaceae bacterium bog_1110 | GCA_003139775.1 | 3.42 | 206 | 43.25 | p_Desulfobacterota;c_Syntrophia;o_Syntrophales;f_Smithellaceae;g_Smithella;s_Smithella sp003139775 | 88.28 | 1.29 |
| Syntrophaceae bacterium bog_1155 | GCA_003165115.1 | 3.91 | 521 | 58.82 | p_Desulfobacterota_G;c_Syntrophorhabdia;o_Syntrophorhabdales;f_WCHB1-27;g_BOG-1155;s_BOG-1155 sp003165115 | 91.76 | 10.88 |
| Syntrophaceae bacterium CG2_30_49_12 | GCA_001873675.1 | 2.03 | 223 | 48.47 | p_Desulfobacterota;c_Syntrophia;o_Syntrophales;f_CG2-30-49-12;g_CG2-30-49-12;s_CG2-30-49-12 sp001873675 | 84.26 | 2.15 |
| Syntrophaceae bacterium CG2_30_58_14 | GCA_001873745.1 | 3.04 | 204 | 57.77 | p_Desulfobacterota;c_Syntrophia;o_Syntrophales;f_UBA5619;g_UBA5619;s_UBA5619 sp001873745 | 95.82 | 6.47 |
| Syntrophaceae bacterium fen_1087 | GCA_003161855.1 | 1.81 | 197 | 50.66 | p_Desulfobacterota;c_Syntrophia;o_Syntrophales;f_Fen-1087;g_Fen-1087;s_Fen-1087 sp003161855 | 95.81 | 0.65 |
| Syntrophaceae bacterium fen_1141 | GCA_003154025.1 | 5.73 | 574 | 56.46 | p_Desulfobacterota;c_SM23-61;o_SM23-61;f_SM23-61;g_SM23-61;s_SM23-61 sp003154025 | 96.97 | 1.82 |
| Syntrophaceae bacterium fen_1145 | GCA_003153775.1 | 3.13 | 98 | 49.99 | p_Desulfobacterota;c_Syntrophia;o_Syntrophales;f_UBA2185;g_Fen-1135;s_Fen-1135 sp003153775 | 91.82 | 0.91 |
| Syntrophaceae bacterium fen_1160 | GCA_003142835.1 | 2.49 | 188 | 44.35 | p_Desulfobacterota;c_Syntrophia;o_Syntrophales;f_Smithellaceae;g_FEN-1160;s_FEN-1160 sp003142835 | 87.74 | 3.44 |
| Syntrophaceae bacterium fen_1164 | GCA_003142635.1 | 3.47 | 260 | 49.56 | p_Desulfobacterota;c_Syntrophia;o_Syntrophales;f_UBA2185;g_Fen-1135;s_Fen-1135 sp003142635 | 91.82 | 0.00 |
| Syntrophaceae bacterium fen_1165 | GCA_003141615.1 | 3.54 | 218 | 43.67 | p_Desulfobacterota;c_Syntrophia;o_Syntrophales;f_Smithellaceae;g_Smithella;s_Smithella sp003141615 | 97.42 | 3.23 |
| Syntrophaceae bacterium fen_1166 | GCA_003142535.1 | 2.93 | 70 | 43.42 | p_Desulfobacterota;c_Syntrophia;o_Syntrophales;f_Smithellaceae;g_Fen-1166;s_Fen-1166 sp003142535 | 99.58 | 0.84 |
| Syntrophaceae bacterium fen_1168 | GCA_003142295.1 | 3.28 | 250 | 48 | p_Desulfobacterota;c_Syntrophia;o_Syntrophales;f_UBA2185;g_Fen-1135;s_Fen-1135 sp003142295 | 96.36 | 3.64 |
| Syntrophaceae bacterium fen_1170 | GCA_003141195.1 | 3.27 | 108 | 57.46 | p_Desulfobacterota;c_Syntrophia;o_Syntrophales;f_UBA5619;g_UBA5619;s_UBA5619 sp003141195 | 65.48 | 2.26 |
| Syntrophaceae bacterium PtaB.Bin038 | GCA_002067855.1 | 2.55 | 433 | 64.54 | p_Desulfobacterota;c_Syntrophia;o_Syntrophales;f_UBA4778;g_UBA2192;s_UBA2192 sp002067855 | 93.55 | 1.29 |
| Syntrophaceae bacterium UBA10514 | GCA_003508565.1 | 2.25 | 298 | 45.94 | p_Desulfobacterota;c_Syntrophia;o_Syntrophales;f_UBA2185;g_UBA2185;s_UBA2185 sp003508565 | 94.35 | 3.25 |
| Syntrophaceae bacterium UBA1062 | GCA_002316295.1 | 3.13 | 295 | 58.13 | p_Desulfobacterota;c_Desulfomonilia;o_UBA1062;f_UBA1062;g_UBA1062;s_UBA1062 sp002316295 | 80.27 | 4.03 |
| Syntrophaceae bacterium UBA1411 | GCA_002304995.1 | 3.42 | 148 | 57.94 | p_Desulfobacterota;c_Syntrophia;o_Syntrophales;f_Smithellaceae;g_UBA8904;s_UBA8904 sp002304995 | 93.44 | 7.74 |
| Syntrophaceae bacterium UBA2188 | GCA_002327985.1 | 3.98 | 186 | 45.72 | p_Desulfobacterota;c_BSN033;o_BSN033;f_UBA1163;g_UBA1163;s_UBA1163 sp002327985 | 100.00 | 0.00 |
| Syntrophaceae bacterium UBA2192 | GCA_002327315.1 | 3.66 | 227 | 59.42 | p_Desulfobacterota;c_Syntrophia;o_Syntrophales;f_UBA4778;g_UBA2192;s_UBA2192 sp002327315 | 92.74 | 2.17 |
| Syntrophaceae bacterium UBA2210 | GCA_002327865.1 | 2.21 | 60 | 48.11 | p_Desulfobacterota;c_Syntrophia;o_Syntrophales;f_UBA2210;g_UBA2210;s_UBA2210 sp002327865 | 96.45 | 1.33 |
| Syntrophaceae bacterium UBA2251 | GCA_002347815.1 | 2.73 | 85 | 62.98 | p_Desulfobacterota;c_Syntrophia;o_Syntrophales;f_UBA2251;g_UBA2251;s_UBA2251 sp002347815 | 93.91 | 5.22 |
| Syntrophaceae bacterium UBA2280 | GCA_002347685.1 | 3.87 | 71 | 51.55 | p_Desulfobacterota;c_Syntrophia;o_Syntrophales;f_Smithellaceae;g_UBA8904;s_UBA8904 sp002347685 | 99.35 | 0.65 |
| Syntrophaceae bacterium UBA3084 | GCA_002367335.1 | 2.10 | 237 | 42.36 | p_Desulfobacterota;c_Syntrophia;o_Syntrophales;f_UBA6807;g_UBA3084;s_UBA3084 sp002367335 | 91.17 | 8.06 |
| Syntrophaceae bacterium UBA4054 | GCA_002383495.1 | 2.66 | 67 | 53.14 | p_Desulfobacterota;c_Syntrophia;o_Syntrophales;f_UBA4778;g_UBA4054;s_UBA4054 sp002383495 | 98.06 | 1.29 |
| Syntrophaceae bacterium UBA4059 | GCA_002382045.1 | 2.95 | 229 | 49.21 | p_Desulfobacterota;c_Syntrophia;o_Syntrophales;f_Smithellaceae;g_UBA8904;s_UBA8904 sp002382045 | 95.38 | 1.94 |
| Syntrophaceae bacterium UBA4767 | GCA_002403385.1 | 1.83 | 82 | 45.92 | p_Desulfobacterota;c_Syntrophia;o_Syntrophales;f_UBA4767;g_UBA4767;s_UBA4767 sp002403385 | 92.42 | 4.30 |
| Syntrophaceae bacterium UBA4778 | GCA_002403175.1 | 3.44 | 342 | 46.45 | p_Desulfobacterota;c_Syntrophia;o_Syntrophales;f_UBA4778;g_UBA4778;s_UBA4778 sp002403175 | 88.50 | 5.16 |
| Syntrophaceae bacterium UBA4810 | GCA_002402545.1 | 2.31 | 109 | 44.61 | p_Desulfobacterota;c_Syntrophia;o_Syntrophales;f_Smithellaceae;g_UBA4810;s_UBA4810 sp002402545 | 94.03 | 1.51 |
| Syntrophaceae bacterium UBA5619 | GCA_002424545.1 | 2.60 | 135 | 58.82 | p_Desulfobacterota;c_Syntrophia;o_Syntrophales;f_UBA5619;g_UBA5619;s_UBA5619 sp002424545 | 96.22 | 0.84 |
| Syntrophaceae bacterium UBA5744 | GCA_002419215.1 | 3.47 | 154 | 60.36 | p_Desulfobacterota;c_Syntrophia;o_Syntrophales;f_UBA4778;g_UBA2192;s_UBA2192 sp002419215 | 86.03 | 1.61 |
| Syntrophaceae bacterium UBA5761 | GCA_002418885.1 | 2.03 | 89 | 55.6 | p_Desulfobacterota;c_Syntrophia;o_Syntrophales;f_UBA4778;g_UBA2207;s_UBA2207 sp002418885 | 87.13 | 0.65 |

| Genome name | NCBI/IMG accession | Size, Mbp | Scaff, no. | GC, % | GTDB Taxonomy | Completeness, % | Contamination, % |
|---|---|---|---|---|---|---|---|
| Syntrophaceae bacterium UBA5764 | GCA_002418825.1 | 2.12 | 98 | 45.52 | p__Desulfobacterota;c__Syntrophia;o__Syntrophales;f__Smithellaceae;g__UBA4810;s__UBA4810 sp002418825 | 98.81 | 0.00 |
| Syntrophaceae bacterium UBA6078 | GCA_002428735.1 | 3.00 | 82 | 54.29 | p__Desulfobacterota;c__Syntrophia;o__Syntrophales;f__UBA2210;g__UBA6078;s__UBA6078 sp002428735 | 95.48 | 3.44 |
| Syntrophaceae bacterium UBA6109 | GCA_002428445.1 | 4.19 | 98 | 58.99 | p__Desulfobacterota;c__Syntrophia;o__Syntrophales;f__UBA6109;g__UBA6109;s__UBA6109 sp002428445 | 95.81 | 3.15 |
| Syntrophaceae bacterium UBA6112 | GCA_002423465.1 | 3.18 | 87 | 56.34 | p__Desulfobacterota;c__Syntrophia;o__Syntrophales;f__UBA5619;g__UBA5619;s__UBA5619 sp002423465 | 91.72 | 0.91 |
| Syntrophaceae bacterium UBA6252 | GCA_002441265.1 | 3.31 | 70 | 60.78 | p__Desulfobacterota;c__Syntrophia;o__Syntrophales;f__UBA6807;g__UBA6807;s__UBA6807 sp002441265 | 80.16 | 2.41 |
| Syntrophaceae bacterium UBA6255 | GCA_002441205.1 | 3.19 | 304 | 43.49 | p__Desulfobacterota;c__Syntrophia;o__Syntrophales;f__Smithellaceae;g__Smithella;s__Smithella sp002441205 | 96.13 | 1.94 |
| Syntrophaceae bacterium UBA7520 | GCA_002478265.1 | 2.34 | 40 | 47.73 | p__Desulfobacterota;c__Syntrophia;o__Syntrophales;f__Smithellaceae;g__UBA4810;s__UBA4810 sp002478265 | 98.13 | 1.64 |
| Syntrophaceae bacterium UBA7544 | GCA_002479215.1 | 2.98 | 305 | 43.29 | p__Desulfobacterota;c__Syntrophia;o__Syntrophales;f__Smithellaceae;g__UBA4810;s__UBA4810 sp002479215 | 84.52 | 1.81 |
| Syntrophaceae bacterium UBA8930 | GCA_003506745.1 | 1.81 | 308 | 51.47 | p__Desulfobacterota;c__Syntrophia;o__Syntrophales;f__Smithellaceae;g__UBA8904;s__UBA8904 sp003506745 | 93.07 | 2.73 |
| Syntrophobacter fumaroxidans MPOB | GCF_000014965.1 | 4.99 | 1 | 59.95 | p__Desulfobacterota;c__Syntrophobacteria;o__Syntrophobacterales;f__Syntrophobacteraceae;g__Syntrophobacter;s__Syntrophobacter fumaroxidans | 85.47 | 0.97 |
| Syntrophobacter sp. DG_60 | GCA_001304365.1 | 1.36 | 128 | 39.12 | p__Desulfobacterota;c__Desulfofervidia;o__Desulfofervidales;f__DG-60;g__DG-60;s__DG-60 sp001304365 | 93.55 | 2.96 |
| Syntrophobacter sp. SbD1 | GCA_900290365.1 | 4.67 | 182 | 52.45 | p__Desulfobacterota;c__Syntrophobacteria;o__Syntrophobacterales;f__Syntrophobacteraceae;g__SbD1;s__SbD1 sp900290365 | 92.06 | 5.06 |
| Syntrophobacter sp. SbD2 | GCA_900290385.1 | 1.81 | 216 | 53.27 | p__Desulfobacterota;c__Syntrophobacteria;o__Syntrophobacterales;f__Syntrophobacteraceae;g__SbD1;s__SbD1 sp900290385 | 77.51 | 2.26 |
| Syntrophobacteraceae bacterium bog_1154 | GCA_003165235.1 | 7.07 | 424 | 52.86 | p__Desulfobacterota;c__Syntrophobacteria;o__Syntrophobacterales;f__Syntrophobacteraceae;g__SbD1;s__SbD1 sp003165235 | 65.50 | 0.65 |
| Syntrophobacteraceae bacterium CSSed162cmB_493R1 | GCA_007125915.1 | 2.35 | 269 | 54.87 | p__Desulfobacterota;c__Syntrophobacteria;o__Syntrophobacterales;f__Syntrophobacteraceae;g__SLCH01;s__SLCH01 sp007125915 | 82.46 | 1.75 |
| Syntrophobacteraceae bacterium T3Sed10_239 | GCA_003566995.1 | 3.04 | 159 | 55.34 | p__Desulfobacterota;c__Syntrophobacteria;o__Syntrophobacterales;f__Syntrophobacteraceae;g__SLCH01;s__SLCH01 sp003566995 | 76.12 | 0.00 |
| Syntrophobacterales bacterium Delta_01 | GCA_001603845.1 | 4.29 | 525 | 59.72 | p__Desulfobacterota;c__Syntrophobacteria;o__Syntrophobacterales;f__Syntrophobacteraceae;g__Delta-01;s__Delta-01 sp001603845 | 90.74 | 0.77 |
| Syntrophobacterales bacterium GWC2_56_13 | GCA_001830835.1 | 2.60 | 119 | 56.37 | p__Desulfobacterota;c__Syntrophia;o__Syntrophales;f__UBA5619;g__UBA5619;s__UBA5619 sp001830835 | 77.98 | 3.36 |
| Syntrophorhabdaceae bacterium PtaU1.Bin034 | GCA_002067405.1 | 5.02 | 407 | 53.82 | p__Desulfobacterota_G;c__Syntrophorhabdia;o__Syntrophorhabdales;f__WCHB1-27;g__BOG-1155;s__BOG-1155 sp002067405 | 75.74 | 0.00 |
| Syntrophorhabdus sp. PtaB.Bin047 | GCA_002067235.1 | 3.64 | 269 | 57.63 | p__Desulfobacterota_G;c__Syntrophorhabdia;o__Syntrophorhabdales;f__Syntrophorhabdaceae;g__Delta-02;s__Delta-02 sp002067235 | 99.19 | 1.45 |
| Syntrophus aciditrophicus SB | GCF_000013405.1 | 3.18 | 1 | 51.46 | p__Desulfobacterota;c__Syntrophia;o__Syntrophales;f__Syntrophaceae;g__Syntrophus;s__Syntrophus aciditrophicus | 92.09 | 2.73 |
| Syntrophus gentianae DSM 8423 | GCA_900109885.1 | 3.72 | 71 | 52.59 | p__Desulfobacterota;c__Syntrophia;o__Syntrophales;f__Syntrophaceae;g__Syntrophus;s__Syntrophus gentianae | 93.98 | 4.09 |
| Syntrophus sp. (in: Bacteria) UBA8958 | GCA_003451635.1 | 2.79 | 449 | 52.27 | p__Desulfobacterota;c__Syntrophia;o__Syntrophales;f__UBA8958;g__UBA8958;s__UBA8958 sp003451635 | 89.09 | 2.73 |
| Syntrophus sp. GWC2_56_31 | GCA_001829875.1 | 2.06 | 136 | 55.77 | p__Desulfobacterota;c__Syntrophia;o__Syntrophales;f__UBA5619;g__UBA5619;s__UBA5619 sp001829875 | 82.20 | 0.22 |
| Syntrophus sp. PtaB.Bin001 | GCA_002067815.1 | 2.84 | 582 | 48.91 | p__Desulfobacterota;c__Syntrophia;o__Syntrophales;f__Syntrophaceae;g__Syntrophus;s__Syntrophus sp002067815 | 89.30 | 2.67 |
| Syntrophus sp. PtaU1.Bin005 | GCA_002067745.1 | 2.90 | 165 | 55.11 | p__Desulfobacterota;c__Syntrophia;o__Syntrophales;f__Syntrophaceae;g__Syntrophus;s__Syntrophus sp002067745 | 94.03 | 2.73 |
| Telmatospirillum siberiense 26-4b1 | GCF_002845745.1 | 6.20 | 81 | 62.33 | p__Proteobacteria;c__Alphaproteobacteria;o__Rhodospirillales;f__Magnetospirillaceae;g__Telmatospirillum;s__Telmatospirillum siberiense | 89.28 | 2.09 |
| Terasakiella pusilla DSM 6293 | GCF_000688235.1 | 4.05 | 85 | 50.02 | p__Proteobacteria;c__Alphaproteobacteria;o__Rhodospirillales;f__Terasakiellaceae;g__Terasakiella;s__Terasakiella pusilla | 99.35 | 0.00 |
| Thalassospira mesophila JCM 18969 | GCF_002115755.1 | 4.93 | 94 | 54.25 | p__Proteobacteria;c__Alphaproteobacteria;o__Rhodospirillales;f__Thalassospiraceae;g__Thalassospira;s__Thalassospira mesophila | 99.00 | 0.00 |
| Thermodesulfatator autotrophicus S606 | GCF_001642325.1 | 2.27 | 129 | 43.08 | p__Desulfobacterota;c__Thermodesulfobacteria;o__Thermodesulfobacteriales;f__Thermodesulfatatoraceae;g__Thermodesulfatator;s__autotrophicus | 74.37 | 1.11 |
| Thermodesulfatator indicus DSM 15286 | GCF_000217795.1 | 2.32 | 1 | 42.43 | p__Desulfobacterota;c__Thermodesulfobacteria;o__Thermodesulfobacteriales;f__Thermodesulfatatoraceae;g__Thermodesulfatator;s__Thermodesulfatator indicus | 96.42 | 2.26 |
| Thermodesulfobacteriaceae bacterium UBA6232 | GCA_002441555.1 | 1.66 | 46 | 37.32 | p__Desulfobacterota;c__Thermodesulfobacteria;o__Thermodesulfobacteriales;f__Thermodesulfobacteriaceae;g__Caldimicrobium;s__Caldimicrobium sp002441555 | 95.84 | 1.29 |
| Thermodesulfobacterium commune DSM 2178 | GCF_000734015.1 | 1.76 | 1 | 36.98 | p__Desulfobacterota;c__Thermodesulfobacteria;o__Thermodesulfobacteriales;f__Thermodesulfobacteriaceae;g__Thermodesulfobacterium;s__commune | 70.48 | 0.00 |
| Thermodesulfobacterium hveragerdense DSM 12571 | GCF_000423845.1 | 1.73 | 42 | 37.1 | p__Desulfobacterota;c__Thermodesulfobacteria;o__Thermodesulfobacteriales;f__Thermodesulfobacteriaceae;g__Thermodesulfobacterium;s__hveragerdense | 78.55 | 2.58 |
| Thermodesulfobacterium thermophilum DSM 1276 | GCF_000421605.1 | 1.79 | 46 | 37.07 | p__Desulfobacterota;c__Thermodesulfobacteria;o__Thermodesulfobacteriales;f__Thermodesulfobacteriaceae;g__Thermodesulfobacterium;s__thermophilum | 90.80 | 4.09 |
| Thermodesulfovibrio aggregans TGE-P1 | GCF_001514535.1 | 2.00 | 3 | 34.89 | p__Nitrospirota;c__Thermodesulfovibrionia;o__Thermodesulfovibrionales;f__Thermodesulfovibrionaceae;g__Thermodesulfovibrio;s__aggregans | 94.53 | 0.00 |
| Thermodesulfovibrio hydrogeniphilus DSM_18151 | 2574179746 | 2.12 | 104 | 35.81 | p__Nitrospirota;c__Thermodesulfovibrionia;o__Thermodesulfovibrionales;f__Thermodesulfovibrionaceae;g__;s__ | 99.03 | 0.00 |
| Thermodesulfovibrio sp. N1 | GCF_001707915.1 | 1.92 | 116 | 32.99 | p__Nitrospirota;c__Thermodesulfovibrionia;o__Thermodesulfovibrionales;f__Thermodesulfovibrionaceae;g__Thermodesulfovibrio;s__sp001707915 | 90.62 | 0.89 |
| Thermodesulfovibrio thiophilus DSM 17215 | GCF_000423865.1 | 1.87 | 18 | 34.43 | p__Nitrospirota;c__Thermodesulfovibrionia;o__Thermodesulfovibrionales;f__Thermodesulfovibrionaceae;g__Thermodesulfovibrio;s__thiophilus | 99.16 | 3.64 |
| Thermodesulfovibrio yellowstonii DSM 11347 | GCF_000020985.1 | 2.00 | 1 | 34.13 | p__Nitrospirota;c__Thermodesulfovibrionia;o__Thermodesulfovibrionales;f__Thermodesulfovibrionaceae;g__Thermodesulfovibrio;s__yellowstonii | 85.14 | 1.68 |
| Thiomicrospira pelophila DSM 1534 | GCF_000711195.1 | 2.11 | 1 | 44.46 | p__Proteobacteria;c__Gammaproteobacteria;o__Thiomicrospirales;f__Thiomicrospiraceae;g__Thiomicrospira;s__Thiomicrospira pelophila | 94.14 | 1.82 |
| Unclassified Nitrospina Bin 25 | 2651870060 | 4.16 | 431 | 37.69 | p__Nitrospinota;c__Nitrospinia;o__Nitrospinales;f__;g__;s__ | 92.31 | 4.27 |
| uncultured Desulfobacteraceae bacterium CR-1 | GCA_900659855.1 | 3.25 | 52 | 54.03 | p__Desulfobacterota;c__Desulfobacteria;o__Desulfobacterales;f__CR-1;g__CR-1;s__CR-1 sp900659855 | 87.88 | 0.65 |
| uncultured Desulfobacterium sp. TRIP AH-1 | GCA_900258555.1 | 5.43 | 28 | 47.01 | p__Desulfobacterota;c__Desulfobacteria;o__Desulfatiglandales;f__Desulfatiglandaceae;g__UBA5623;s__UBA5623 sp900258555 | 83.55 | 2.26 |
| uncultured Desulfovibrio sp. RUG514 | GCA_900319575.1 | 2.14 | 253 | 59.19 | p__Desulfobacterota;c__Desulfovibrionia;o__Desulfovibrionales;f__Desulfovibrionaceae;g__Desulfovibrio;s__Desulfovibrio sp900319575 | 91.31 | 0.91 |
| uncultured Desulfovibrio sp. UMGS1330 | GCA_900550745.1 | 2.38 | 425 | 61.53 | p__Desulfobacterota;c__Desulfovibrionia;o__Desulfovibrionales;f__Desulfovibrionaceae;g__Bilophila;s__Bilophila sp900550745 | 88.39 | 0.65 |
| uncultured Desulfovibrio sp. UMGS1580 | GCA_900553065.1 | 1.92 | 70 | 58.43 | p__Desulfobacterota;c__Desulfovibrionia;o__Desulfovibrionales;f__Desulfovibrionaceae;g__Mailhella;s__Mailhella sp900553065 | 96.13 | 0.84 |
| uncultured Desulfovibrio sp. UMGS1890 | GCA_900555975.1 | 1.51 | 319 | 59.28 | p__Desulfobacterota;c__Desulfovibrionia;o__Desulfovibrionales;f__Desulfovibrionaceae;g__Mailhella;s__Mailhella sp900555975 | 77.14 | 0.96 |
| uncultured Desulfovibrio sp. UMGS1966 | GCA_900556755.1 | 1.38 | 32 | 62.94 | p__Desulfobacterota;c__Desulfovibrionia;o__Desulfovibrionales;f__Desulfovibrionaceae;g__Desulfovibrio;s__Desulfovibrio sp900556755 | 64.52 | 0.00 |
| uncultured Desulfovibrio sp. UMGS250 | GCA_900540515.1 | 3.43 | 68 | 61.58 | p__Desulfobacterota;c__Desulfovibrionia;o__Desulfovibrionales;f__Desulfovibrionaceae;g__Desulfovibrio;s__Desulfovibrio sp900540515 | 95.70 | 5.47 |
| uncultured Desulfovibrio sp. UMGS847 | GCA_900546145.1 | 2.46 | 72 | 58.52 | p__Desulfobacterota;c__Desulfovibrionia;o__Desulfovibrionales;f__Desulfovibrionaceae;g__Desulfovibrio;s__Desulfovibrio sp900546145 | 73.21 | 0.00 |
| Uncultured microorganism SbSrfc.SA12.01.D19 | 3300022116 | 2.50 | 175 | 52.60 | p__Desulfobacterota;c__Desulfobulbia;o__Desulfobulbales;f__BM004;g__;s__ | 49.13 | 0.00 |
| uncultured Nitrosospira sp. NOB2.fa.gz | GCA_900696505.1 | 3.25 | 413 | 58.19 | p__Nitrospirota;c__Nitrospiria;o__Nitrospirales;f__Nitrospiraceae;g__Nitrospira_A;s__Nitrospira_A sp900696505 | 98.94 | 1.26 |
| Zetaproteobacteria bacterium nPCbin1 | GCA_015231855.1 | 2.02 | 68 | 47.46 | p__Proteobacteria;c__Zetaproteobacteria;o__Mariprofundales;f__Mariprofundaceae;g__GCA-2753275;s__GCA-2753275 sp002753275 | 99.35 | 0.97 |
| Zetaproteobacteria bacterium PCbin4 | GCA_002753275.1 | 1.86 | 55 | 47.58 | p__Proteobacteria;c__Zetaproteobacteria;o__Mariprofundales;f__Mariprofundaceae;g__GCA-2753275;s__GCA-2753275 sp002753275 | 98.81 | 0.00 |
| Zetaproteobacteria bacterium SZUA-181 | GCA_003228735.1 | 1.88 | 100 | 55.29 | p__Proteobacteria;c__Zetaproteobacteria;o__Mariprofundales;f__Mariprofundaceae;g__SZUA-181;s__SZUA-181 sp003228735 | 99.35 | 0.00 |

**Supplementary table S2.** Results of reconciliations for protein trees and concatenated protein tree obtained by Notung and Ranger-DTL tools. LCMA – last common magnetotactic ancestor, LCDA – last common *Dissulfurispiraceae* ancestor, HGT – horizontal gene transfer.

| **Analysis** | ***Magnetobacteriaceae* group** | ***Dissulfurispiraceae* group** | ***man*-containing *Thermodesulfobacteriota* group** |
|---|---|---|---|
| Man1 Ranger-DTL | Vertical inheritance from LCMA | HGT from *Magnetobacteriaceae* group | - |
| Man1 Notung | Vertical inheritance from LCMA | HGT from *Magnetobacteriaceae* group | - |
| Man2 Ranger-DTL | Vertical inheritance from LCMA | HGT from *Magnetobacteriaceae* group | HGT from *Magnetobacteriaceae* group |
| Man2 Notung | Vertical inheritance from LCMA | HGT from *Magnetobacteriaceae* group | Vertical inheritance from LCMA |
| Man3 Ranger-DTL | A reconciliation was not done due to the sequences shortness, which makes it difficult to avoid misinterpreting the results | | |
| Man3 Notung | | | |
| Man4 Ranger-DTL | Vertical inheritance from LCMA | HGT from *Magnetobacteriaceae* group | - |
| Man4 Notung | Vertical inheritance from LCMA | HGT from *Magnetobacteriaceae* group | - |
| Man5 Ranger-DTL | Vertical inheritance from LCMA | HGT from *Magnetobacteriaceae* group | - |
| Man5 Notung | Vertical inheritance from LCMA | Vertical inheritance from LCMA | - |
| Man6 Ranger-DTL | HGT from LCDA | Vertical inheritance from LCMA | HGT from LCDA |
| Man6 Notung | HGT from LCDA | Vertical inheritance from LCMA | HGT from LCDA |
| Mad2 Ranger-DTL | Vertical inheritance from LCMA | HGT from *Magnetobacteriaceae* group | HGT from *Magnetobacteriaceae* group |
| Mad2 Notung | Vertical inheritance from LCMA | HGT from *Magnetobacteriaceae* group | HGT from LCDA |
| Mad10 Ranger-DTL | Vertical inheritance from LCMA | HGT from *Magnetobacteriaceae* group | HGT from *Magnetobacteriaceae* group |
| Mad10 Notung | Vertical inheritance from LCMA | HGT from *Magnetobacteriaceae* group | HGT from *Magnetobacteriaceae* group |
| Mad23 Ranger-DTL | HGT from *Bdellovibrionota* | HGT from *Magnetobacteriaceae* group | HGT from *Magnetobacteriaceae* group |
| Mad23 Notung | HGT from *Thermodesulfobacteriota* | HGT from *Magnetobacteriaceae* group | HGT from *Magnetobacteriaceae* group |
| Mad24 Ranger-DTL | Vertical inheritance from LCMA | HGT from *Magnetobacteriaceae* group | HGT from *Magnetobacteriaceae* group |
| Mad24 Notung | HGT from *Thermodesulfobacteriota* | HGT from *Magnetobacteriaceae* group | Vertical inheritance from LCMA |
| Mad25 Ranger-DTL | Vertical inheritance from LCMA | HGT from *Magnetobacteriaceae* group | HGT from *Magnetobacteriaceae* group |
| Mad25 Notung | Vertical inheritance from LCMA | HGT from *Magnetobacteriaceae* group | HGT from *Thermodesulfobacteriota* |
| Mad26 Ranger-DTL | HGT from *Thermodesulfobacteriota* | HGT from *Magnetobacteriaceae* group | HGT from *Magnetobacteriaceae* group |
| Mad26 Notung | HGT from *Thermodesulfobacteriota* | HGT from *Magnetobacteriaceae* group | HGT from *Magnetobacteriaceae* group |

| **Analysis** | ***Magnetobacteriaceae* group** | ***Dissulfurispiraceae* group** | ***man*-containing *Thermodesulfobacteriota* group** |
|---|---|---|---|
| Mad31 Ranger-DTL | Vertical inheritance from LCMA | HGT from *Magnetobacteriaceae* group | HGT from *Magnetobacteriaceae* group |
| Mad31 Notung | Vertical inheritance from LCMA | HGT from *Magnetobacteriaceae* group | HGT from *Magnetobacteriaceae* group |
| MamA Ranger-DTL | HGT from *Bdellovibrionota*/ *Hydrogenedentota* | HGT from *Magnetobacteriaceae* group | HGT from *Magnetobacteriaceae* group |
| MamA Notung | HGT from *Bdellovibrionota* | HGT from *Magnetobacteriaceae* group | HGT from *Magnetobacteriaceae* group |
| MamB Ranger-DTL | HGT from *man*-containing *Thermodesulfobacteriota* group | HGT from *Magnetobacteriaceae* group | HGT from *Riflebactetria* |
| MamB Notung | HGT from *Riflebactetria* | HGT from *Magnetobacteriaceae* group | HGT from *Magnetobacteriaceae* group |
| MamK Ranger-DTL | HGT from *Thermodesulfobacteriota* | HGT from *Magnetobacteriaceae* group | HGT from *Magnetobacteriaceae* group |
| MamK Notung | HGT from *Thermodesulfobacteriota* | HGT from *Magnetobacteriaceae* group | HGT from *Magnetobacteriaceae* group |
| MamM Ranger-DTL | HGT from *man*-containing *Thermodesulfobacteriota* group | HGT from *Magnetobacteriaceae* group | HGT from *Planctomycetota* |
| MamM Notung | HGT from *man*-containing *Thermodesulfobacteriota* group | HGT from *Magnetobacteriaceae* group | HGT from *Thermodesulfobacteriota* |
| MamP Ranger-DTL | HGT from *Placntomycetota* | HGT from *Magnetobacteriaceae* group | HGT from *Magnetobacteriaceae* group |
| MamP Notung | HGT from *Omnitrophota* | HGT from *Magnetobacteriaceae* group | HGT from *Magnetobacteriaceae* group |
| MamQ Ranger-DTL | HGT from *Riflebacteria* | HGT from *Magnetobacteriaceae* group | HGT from *Magnetobacteriaceae* group |
| MamQ Notung | HGT from *man*-containing *Thermodesulfobacteriota* group | HGT from *Magnetobacteriaceae* group | HGT from *Planctomycetota* |
| MamI Ranger-DTL | HGT from *man*-containing *Thermodesulfobacteriota* group | HGT from *Magnetobacteriaceae* group | HGT from *Omnitrophota* |
| MamI Notung | HGT from *Thermodesulfobacteriota* | HGT from *Magnetobacteriaceae* group | HGT from *Magnetobacteriaceae* group |
| MamE Ranger-DTL | HGT from *man*-containing *Thermodesulfobacteriota* group | HGT from *Magnetobacteriaceae* group | HGT from *Omnitrophota* |
| MamE Notung | HGT from *man*-containing *Thermodesulfobacteriota* group | HGT from *Magnetobacteriaceae* group | HGT from *Nitrospinota* |
| MamO-Cter Ranger-DTL | HGT from *Bdellovibrionota* | HGT from *Magnetobacteriaceae* group | HGT from *Magnetobacteriaceae* group |
| MamO-Cter Notung | HGT from *man*-containing *Thermodesulfobacteriota* group | HGT from *Magnetobacteriaceae* group | HGT from *Planctomycetota* |
| MamQ-2 Ranger-DTL | Vertical inheritance from LCMA | HGT from *Magnetobacteriaceae* group | HGT from *Magnetobacteriaceae* group |
| MamQ-2 Notung | Vertical inheritance from LCMA | HGT from *Magnetobacteriaceae* group | HGT from *Magnetobacteriaceae* group |
| **Concatenated Ranger-DTL** | HGT from *Bdellovibrionota* | HGT from *Magnetobacteriaceae* group | HGT from *Magnetobacteriaceae* group |
| **Concatenated Notung** | HGT from *man*-containing *Thermodesulfobacteriota* group | HGT from *Magnetobacteriaceae* group | HGT from *Bdellovibrionota* |

**Supplementary table S3.** Reconstructed genomes statistics.

| Attribute | LBB01 | | LBB02 | | LBB04 | |
|---|---|---|---|---|---|---|
| | **Value** | **% of Total** | **Value** | **% of Total** | **Value** | **% of Total** |
| Genome size, bp | 3,273,455 | 100.0 | 3,471,208 | 100.0 | 4,495,917 | 100.0 |
| DNA coding, bp | 2,939,340 | 89.8 | 3,089,132 | 89.0 | 3,786,226 | 84.2 |
| DNA G+C, bp | 1,373,588 | 42.0 | 1,629,784 | 47.0 | 2,267,373 | 50.4 |
| DNA scaffolds | 1 | 100.0 | 142 | 100.0 | 2,129 | 100.0 |
| Total genes | 3,132 | 100.0 | 3,355 | 100.0 | 5,601 | 100.0 |
| Protein coding genes | 3,021 | 96.5 | 3,298 | 98.3 | 5,379 | 96.0 |
| RNA genes | 56 | 1.8 | 37 | 1.1 | 34 | 0.6 |
| Pseudo genes | 55 | 1.8 | 20 | 0.6 | 188 | 3.4 |
| Genes with function prediction | 2,474 | 79.0 | 2,521 | 75.1 | 3,825 | 68.3 |
| Genes assigned to COGs | 2,373 | 75.8 | 2,445 | 72.9 | 4,051 | 72.3 |
| Genes with Pfam domains | 2,473 | 79.0 | 2,576 | 76.8 | 3,844 | 68.6 |
| Genes with signal peptides | 305 | 9.7 | 412 | 12.3 | 512 | 9.1 |
| Genes with transmembrane helices | 807 | 25.8 | 943 | 28.1 | 1284 | 22.9 |
| CRISPR repeats | 6 | - | 11 | - | 8 | - |
| $N_{50}$, bp | 3,273,455 | | 44,214 | | 2,222 | |
| CheckM completeness, % | 99.03 | | 91.52 | | 60.18 | |
| CheckM contamination, % | 0.91 | | 0 | | 0 | |
| **GTDB affiliations** | | | | | | |
| Phylum | *Nitrospirota* | | *Nitrospirota* | | *Desulfobacterota* | |
| Class | *Thermodesulfovibrionia* | | *Thermodesulfovibrionia* | | *Syntrophia* | |
| Order | *Thermodesulfovibrionales* | | *Thermodesulfovibrionales* | | *Syntrophales* | |
| Family | *Ca*. Magnetobacteriaceae | | *Ca*. Magnetobacteriaceae | | UBA2185 | |
| Genus | - | | HCH-1 | | - | |
| Species | - | | - | | - | |

**Supplementary table S4.** AAI and POCP values between *Nitrospirota* genomes.

| | | | POCP | | | | | | | | | | | | | | | | | | | | | | | | | | | | | | | | | | | | |
|---|---|---|---|---|---|---|---|---|---|---|---|---|---|---|---|---|---|---|---|---|---|---|---|---|---|---|---|---|---|---|---|---|---|---|---|---|---|---|---|
| | № | Genome | 1 | 2 | 3 | 4 | 5 | 6 | 7 | 8 | 9 | 10 | 11 | 12 | 13 | 14 | 15 | 16 | 17 | 18 | 19 | 20 | 21 | 22 | 23 | 24 | 25 | 26 | 27 | 28 | 29 | 30 | 31 | 32 | 33 | 34 | 35 | 36 | 37 |
| | 1 | *Ca* . Magnetomonas plexicatena LBB01 | 100.0 | 76.7 | 82.0 | 79.4 | 81.8 | 59.0 | 60.4 | 60.9 | 61.3 | 61.4 | 46.4 | 51.1 | 48.3 | 58.3 | 57.9 | 51.3 | 53.8 | 55.4 | 55.2 | 52.9 | 43.3 | 54.8 | 55.5 | 55.2 | 49.6 | 54.6 | 54.0 | 51.6 | 52.2 | 33.1 | 45.4 | 40.0 | 33.0 | 32.6 | 34.0 | 33.9 | 32.9 |
| | 2 | *Ca* . Magnetomonas sp. nDJH6bin1 | 85.4 | 100.0 | 82.4 | 79.5 | 81.9 | 55.4 | 57.2 | 56.5 | 56.7 | 56.8 | 44.0 | 48.1 | 45.7 | 52.7 | 52.2 | 48.0 | 50.2 | 51.3 | 51.5 | 49.7 | 40.3 | 50.6 | 51.0 | 50.7 | 45.8 | 50.4 | 49.5 | 49.1 | 48.0 | 31.7 | 42.2 | 37.2 | 31.1 | 30.4 | 32.2 | 31.5 | 30.8 |
| | 3 | *Ca* . Magnetomonas sp. nDJH13bin19 | 86.3 | 94.3 | 100.0 | 94.3 | 97.3 | 59.8 | 61.9 | 59.5 | 59.9 | 60.0 | 47.0 | 50.2 | 48.4 | 57.2 | 56.9 | 51.1 | 53.1 | 54.3 | 55.6 | 53.4 | 42.1 | 54.6 | 55.1 | 54.7 | 48.3 | 53.8 | 53.0 | 53.1 | 51.1 | 32.5 | 44.9 | 40.2 | 33.0 | 32.3 | 34.0 | 34.0 | 32.5 |
| | 4 | *Ca* . Magnetomonas sp. nDJH8bin8 | 86.2 | 94.1 | 99.9 | 100.0 | 93.4 | 58.4 | 61.0 | 58.6 | 58.8 | 59.0 | 50.4 | 48.7 | 46.8 | 55.8 | 55.8 | 49.3 | 52.9 | 53.2 | 54.1 | 52.0 | 40.7 | 54.2 | 54.0 | 53.6 | 47.0 | 52.5 | 51.7 | 52.7 | 49.8 | 31.3 | 43.5 | 38.3 | 31.4 | 30.5 | 32.2 | 32.1 | 30.6 |
| | 5 | *Ca* . Magnetomonas sp. nDJH14bin5 | 86.2 | 94.2 | 100.0 | 99.9 | 100.0 | 59.4 | 61.0 | 59.0 | 59.4 | 59.5 | 46.3 | 49.7 | 47.8 | 56.6 | 56.3 | 50.7 | 52.5 | 53.9 | 54.7 | 52.7 | 42.7 | 53.8 | 54.6 | 54.2 | 47.9 | 53.3 | 52.3 | 52.8 | 50.4 | 32.4 | 44.1 | 39.7 | 32.7 | 32.3 | 33.6 | 33.8 | 32.4 |
| | 6 | *Ca* . Magnetominusculus linsii LBB02 | 60.8 | 60.3 | 61.1 | 61.0 | 60.9 | 100.0 | 76.1 | 73.5 | 73.9 | 74.2 | 55.7 | 63.7 | 58.3 | 66.6 | 66.0 | 62.2 | 52.6 | 53.1 | 52.9 | 50.4 | 42.5 | 52.2 | 53.0 | 52.6 | 46.7 | 52.2 | 50.6 | 48.4 | 48.0 | 30.6 | 41.7 | 37.1 | 29.9 | 29.6 | 31.0 | 31.0 | 30.0 |
| | 7 | *Ca* . Magnetominusculus sp. nDJH13bin15 | 61.5 | 61.7 | 62.3 | 62.1 | 62.1 | 77.3 | 100.0 | 73.9 | 74.6 | 74.8 | 61.0 | 65.4 | 61.5 | 66.4 | 66.4 | 64.0 | 54.0 | 54.8 | 55.7 | 53.1 | 41.8 | 53.6 | 54.4 | 54.0 | 48.6 | 53.5 | 52.2 | 52.2 | 50.0 | 31.2 | 44.0 | 39.3 | 31.8 | 31.5 | 33.3 | 32.9 | 32.0 |
| | 8 | *Ca* . Magnetominusculus xianensis HCH-1 | 61.9 | 61.3 | 61.6 | 61.6 | 61.3 | 76.8 | 80.1 | 100.0 | 96.9 | 97.2 | 57.7 | 65.4 | 60.8 | 70.8 | 71.0 | 64.8 | 54.8 | 55.9 | 54.4 | 51.8 | 43.3 | 55.0 | 56.4 | 56.2 | 50.0 | 54.8 | 54.1 | 49.9 | 50.5 | 30.4 | 42.4 | 39.1 | 31.3 | 31.8 | 33.0 | 32.4 | 31.7 |
| | 9 | *Ca.* Magnetominusculus xianensis nHCHbin2 | 61.8 | 61.2 | 61.6 | 61.4 | 61.3 | 76.8 | 80.2 | 100.0 | 100.0 | 99.2 | 58.1 | 65.8 | 60.7 | 71.5 | 71.7 | 65.3 | 54.8 | 55.7 | 54.4 | 51.7 | 43.2 | 55.1 | 56.5 | 56.3 | 50.0 | 54.8 | 54.0 | 50.3 | 50.5 | 30.4 | 42.4 | 39.0 | 31.0 | 31.6 | 32.8 | 32.3 | 31.7 |
| | 10 | *Ca* . Magnetominusculus xianensis HCHbin1 | 61.7 | 61.1 | 61.5 | 61.4 | 61.3 | 76.8 | 80.1 | 100.0 | 100.0 | 100.0 | 58.2 | 66.0 | 61.0 | 71.5 | 71.9 | 65.6 | 54.8 | 55.8 | 54.4 | 51.7 | 43.3 | 55.2 | 56.6 | 56.3 | 50.0 | 54.9 | 54.1 | 50.2 | 50.7 | 30.4 | 42.5 | 38.9 | 31.1 | 31.6 | 32.8 | 32.3 | 31.6 |
| | 11 | *Ca* . Magnetominusculus sp. nDJH14bin7 | 61.5 | 61.5 | 62.1 | 65.2 | 61.9 | 76.7 | 81.4 | 79.7 | 79.8 | 79.7 | 100.0 | 51.6 | 49.5 | 51.0 | 51.8 | 49.7 | 42.1 | 42.0 | 42.7 | 40.7 | 30.4 | 42.2 | 41.8 | 41.5 | 36.8 | 40.5 | 39.9 | 40.9 | 38.7 | 22.8 | 32.5 | 30.2 | 24.1 | 23.3 | 24.8 | 24.9 | 23.6 |
| | 12 | *Ca* . Magnetominusculus sp. nDJH8bin13 | 60.7 | 60.4 | 60.9 | 60.8 | 60.8 | 76.1 | 80.0 | 78.7 | 78.9 | 78.9 | 86.5 | 100.0 | 60.9 | 58.6 | 59.4 | 58.3 | 44.9 | 45.7 | 45.5 | 44.5 | 39.0 | 43.8 | 44.8 | 44.6 | 40.8 | 44.8 | 44.4 | 43.0 | 40.8 | 27.8 | 35.9 | 33.4 | 26.3 | 26.4 | 27.4 | 27.7 | 26.5 |
| | 13 | *Ca* . Magnetominusculus sp. nDJH13bin15 | 61.8 | 61.1 | 62.2 | 62.0 | 61.9 | 76.9 | 81.4 | 79.6 | 79.6 | 79.6 | 87.5 | 92.0 | 100.0 | 53.3 | 53.2 | 54.5 | 42.5 | 43.9 | 44.6 | 43.1 | 36.8 | 42.0 | 43.1 | 42.8 | 40.8 | 43.3 | 43.6 | 40.6 | 40.4 | 27.2 | 35.0 | 33.8 | 27.4 | 27.4 | 28.8 | 28.9 | 27.6 |
| | 14 | *Ca* . Magnetominusculus sp. nMYbin6 | 61.5 | 60.5 | 61.2 | 60.9 | 61.0 | 73.9 | 76.6 | 77.1 | 77.2 | 77.2 | 76.6 | 75.8 | 76.5 | 100.0 | 92.8 | 58.8 | 53.3 | 54.2 | 51.4 | 48.7 | 42.5 | 52.7 | 53.6 | 53.2 | 47.7 | 53.3 | 52.2 | 47.6 | 46.7 | 29.2 | 39.2 | 36.9 | 29.8 | 29.4 | 31.1 | 30.7 | 29.7 |
| | 15 | *Ca* . Magnetominusculus sp. MYbin6 | 61.8 | 60.7 | 61.4 | 61.1 | 61.2 | 74.0 | 76.7 | 77.1 | 77.1 | 77.1 | 76.5 | 75.7 | 76.5 | 99.9 | 100.0 | 58.0 | 53.1 | 53.6 | 51.3 | 48.5 | 42.0 | 52.8 | 53.6 | 53.2 | 47.7 | 53.0 | 51.8 | 47.5 | 46.5 | 29.2 | 39.7 | 36.4 | 28.7 | 28.5 | 30.2 | 29.9 | 28.5 |
| | 16 | *Ca* . Magnetominusculus sp. nDJH5bin4 | 61.1 | 60.6 | 61.4 | 61.3 | 61.2 | 75.7 | 78.4 | 79.9 | 80.1 | 80.1 | 77.9 | 76.8 | 77.3 | 74.5 | 74.6 | 100.0 | 44.2 | 45.1 | 46.5 | 44.0 | 37.4 | 44.4 | 45.5 | 45.2 | 41.4 | 45.0 | 43.9 | 41.7 | 43.2 | 27.6 | 36.4 | 33.5 | 26.7 | 27.2 | 28.2 | 28.1 | 26.6 |
| | 17 | *Ca* . Magnetobacterium sp. nMYbin4 | 59.7 | 59.6 | 60.0 | 60.1 | 59.8 | 59.9 | 60.7 | 60.5 | 60.4 | 60.4 | 60.2 | 58.9 | 60.5 | 60.4 | 60.4 | 59.6 | 100.0 | 90.8 | 69.2 | 66.5 | 54.8 | 71.3 | 71.4 | 71.3 | 64.8 | 70.8 | 69.8 | 49.6 | 50.3 | 29.3 | 40.9 | 37.6 | 30.8 | 29.9 | 31.8 | 31.1 | 29.8 |
| | 18 | *Ca* . Magnetobacterium sp. MYbinv3 | 60.0 | 59.8 | 60.2 | 60.1 | 60.1 | 60.0 | 60.8 | 60.7 | 60.6 | 60.6 | 60.4 | 59.2 | 60.7 | 60.6 | 60.6 | 59.9 | 99.8 | 100.0 | 71.7 | 68.8 | 56.9 | 72.1 | 73.4 | 72.8 | 66.6 | 72.9 | 72.2 | 51.1 | 51.8 | 30.0 | 41.5 | 38.5 | 32.0 | 31.2 | 33.1 | 32.6 | 31.1 |
| AAI | 19 | *Ca* . Magnetobacterium sp. nDJH15bin2 | 59.8 | 59.5 | 59.9 | 59.9 | 59.5 | 59.6 | 60.8 | 60.3 | 60.3 | 60.2 | 60.6 | 59.1 | 60.7 | 59.8 | 59.9 | 60.1 | 75.7 | 75.8 | 100.0 | 88.1 | 58.9 | 73.3 | 75.2 | 74.5 | 67.9 | 73.8 | 71.7 | 53.1 | 51.4 | 30.5 | 41.6 | 37.9 | 31.2 | 29.9 | 32.2 | 31.4 | 30.6 |
| | 20 | *Ca* . Magnetobacterium sp. nDJH13bin1 | 60.0 | 59.7 | 60.1 | 60.2 | 59.9 | 59.7 | 61.0 | 60.7 | 60.6 | 60.6 | 60.6 | 59.3 | 61.0 | 60.0 | 60.0 | 60.1 | 75.5 | 75.7 | 99.2 | 100.0 | 57.9 | 70.5 | 72.2 | 71.6 | 64.7 | 70.7 | 69.2 | 50.8 | 49.2 | 28.9 | 39.6 | 36.8 | 30.0 | 28.6 | 31.2 | 30.4 | 29.7 |
| | 21 | *Ca* . Magnetobacterium sp. nMYbin3 | 58.1 | 58.2 | 58.0 | 58.1 | 57.8 | 58.9 | 59.0 | 59.1 | 59.1 | 59.1 | 58.7 | 58.4 | 58.5 | 59.0 | 59.2 | 57.7 | 74.2 | 74.4 | 80.9 | 80.5 | 100.0 | 57.7 | 59.4 | 59.1 | 56.6 | 61.6 | 58.0 | 38.9 | 39.6 | 25.7 | 30.8 | 28.9 | 25.0 | 24.8 | 25.1 | 24.5 | 24.6 |
| | 22 | *Ca* . Magnetobacterium cryptolimnobacter XYR | 60.0 | 59.3 | 59.8 | 59.8 | 59.6 | 60.0 | 60.7 | 60.7 | 60.6 | 60.6 | 60.2 | 59.2 | 60.7 | 60.2 | 60.4 | 59.8 | 74.8 | 75.1 | 80.5 | 80.5 | 83.7 | 100.0 | 94.3 | 93.7 | 75.2 | 80.8 | 79.4 | 49.6 | 50.1 | 29.0 | 39.7 | 36.8 | 28.9 | 28.7 | 30.5 | 30.0 | 29.0 |
| | 23 | *Ca* . Magnetobacterium sp. DC0425bin1 | 60.1 | 59.4 | 59.8 | 59.7 | 59.6 | 60.1 | 60.6 | 60.7 | 60.6 | 60.6 | 60.3 | 59.3 | 60.7 | 60.3 | 60.4 | 59.7 | 74.8 | 74.8 | 80.5 | 80.4 | 83.5 | 99.5 | 100.0 | 97.8 | 76.4 | 82.4 | 80.7 | 50.8 | 50.9 | 29.6 | 40.2 | 37.3 | 29.7 | 29.7 | 31.3 | 30.7 | 29.8 |
| | 24 | *Ca* . Magnetobacterium sp. nDC0425bin1 | 60.1 | 59.4 | 59.8 | 59.7 | 59.6 | 60.1 | 60.6 | 60.6 | 60.6 | 60.5 | 60.2 | 59.2 | 60.7 | 60.2 | 60.3 | 59.7 | 74.8 | 74.8 | 80.6 | 80.5 | 83.7 | 99.4 | 100.0 | 100.0 | 76.3 | 82.0 | 80.3 | 50.4 | 50.7 | 29.6 | 39.9 | 37.2 | 29.6 | 29.6 | 31.2 | 30.7 | 29.6 |
| | 25 | *Ca* . Magnetobacterium sp. MYbin2 | 60.2 | 59.2 | 59.9 | 59.7 | 59.6 | 60.1 | 60.5 | 60.5 | 60.4 | 60.4 | 60.6 | 59.0 | 60.6 | 60.0 | 59.9 | 59.8 | 75.0 | 74.9 | 80.7 | 80.4 | 83.1 | 92.1 | 92.0 | 92.0 | 100.0 | 85.0 | 84.7 | 44.9 | 45.7 | 28.3 | 36.8 | 32.6 | 26.9 | 26.3 | 27.6 | 27.5 | 26.8 |
| | 26 | *Ca* . Magnetobacterium sp. nMYbin2 | 59.8 | 59.2 | 59.6 | 59.5 | 59.4 | 59.8 | 60.5 | 60.4 | 60.3 | 60.3 | 60.1 | 59.2 | 60.2 | 60.0 | 59.9 | 59.5 | 75.0 | 75.1 | 80.8 | 80.5 | 83.7 | 92.5 | 92.5 | 92.5 | 99.5 | 100.0 | 89.5 | 49.6 | 49.5 | 29.9 | 39.9 | 35.8 | 29.1 | 28.9 | 30.3 | 29.7 | 28.8 |
| | 27 | *Ca* . Magnetobacterium casensis MYR-1 | 60.3 | 59.8 | 60.2 | 60.0 | 60.0 | 60.4 | 60.9 | 61.1 | 61.0 | 61.0 | 60.8 | 59.7 | 61.1 | 60.4 | 60.4 | 60.1 | 75.1 | 75.2 | 80.9 | 80.7 | 83.7 | 92.5 | 92.4 | 92.5 | 99.7 | 99.8 | 100.0 | 49.0 | 48.9 | 29.5 | 39.7 | 36.4 | 29.9 | 29.6 | 31.1 | 30.7 | 29.8 |
| | 28 | Magnetobacteriaceae bacterium nDJH13bin3 | 60.2 | 59.8 | 60.3 | 60.8 | 60.1 | 59.6 | 61.2 | 60.5 | 60.2 | 60.3 | 61.0 | 59.7 | 60.6 | 59.8 | 60.2 | 59.7 | 61.7 | 61.8 | 61.3 | 61.5 | 59.6 | 61.3 | 61.2 | 61.2 | 61.4 | 61.2 | 61.8 | 100.0 | 45.0 | 27.7 | 38.4 | 34.9 | 27.8 | 27.1 | 28.8 | 28.7 | 27.3 |
| | 29 | Ca. Magnetomicrobium cryptolimnococcus XYC | 57.8 | 57.0 | 57.6 | 57.4 | 57.3 | 56.6 | 57.6 | 57.5 | 57.5 | 57.4 | 57.1 | 56.2 | 57.4 | 56.8 | 56.9 | 56.9 | 57.5 | 57.8 | 57.4 | 57.4 | 55.3 | 57.2 | 57.2 | 57.2 | 57.2 | 57.1 | 57.5 | 57.1 | 100.0 | 31.2 | 42.1 | 38.1 | 31.9 | 31.7 | 33.4 | 33.1 | 32.6 |
| | 30 | Thermodesulfovibrionales bacterium nDJH8bin7 | 55.4 | 54.9 | 55.2 | 55.2 | 55.0 | 55.4 | 55.5 | 55.5 | 55.5 | 55.5 | 55.0 | 53.9 | 54.7 | 54.8 | 54.8 | 55.5 | 55.5 | 55.7 | 55.0 | 55.1 | 53.8 | 54.8 | 55.0 | 55.0 | 55.0 | 54.8 | 55.0 | 54.2 | 54.6 | 100.0 | 71.6 | 29.3 | 23.3 | 24.0 | 25.0 | 24.9 | 23.9 |
| | 31 | Thermodesulfovibrionales bacterium nDJH14bin9 | 56.1 | 55.9 | 56.2 | 56.1 | 55.9 | 55.9 | 56.5 | 56.6 | 56.5 | 56.5 | 56.0 | 54.6 | 55.8 | 55.5 | 55.5 | 55.9 | 56.3 | 56.6 | 56.4 | 56.3 | 54.3 | 56.0 | 56.1 | 56.1 | 56.1 | 55.8 | 56.2 | 55.6 | 55.5 | 99.6 | 100.0 | 39.7 | 32.1 | 32.1 | 33.8 | 33.6 | 32.9 |
| | 32 | *Dissulfurispira thermophila* T55J | 56.6 | 55.9 | 56.3 | 56.1 | 56.0 | 55.8 | 56.8 | 56.4 | 56.4 | 56.3 | 56.2 | 55.5 | 55.8 | 56.0 | 56.1 | 55.8 | 56.4 | 56.6 | 56.1 | 56.3 | 54.4 | 56.5 | 56.4 | 56.4 | 56.4 | 56.2 | 56.5 | 55.8 | 56.0 | 54.3 | 55.8 | 100.0 | 53.2 | 52.9 | 56.6 | 55.1 | 52.7 |
| | 33 | *Thermodesulfovibrio hydrogeniphilus* DSM 18151 | 53.1 | 53.2 | 53.2 | 53.1 | 53.0 | 52.3 | 53.2 | 53.2 | 53.0 | 53.1 | 52.6 | 51.7 | 53.1 | 52.2 | 52.1 | 52.0 | 52.9 | 53.0 | 52.5 | 52.8 | 51.2 | 52.6 | 52.5 | 52.4 | 52.8 | 52.4 | 52.9 | 52.2 | 52.7 | 51.4 | 52.7 | 58.8 | 100.0 | 77.2 | 80.2 | 80.8 | 74.7 |
| | 34 | *Thermodesulfovibrio* sp. N1 | 53.2 | 53.1 | 53.4 | 53.2 | 53.3 | 52.6 | 53.4 | 53.2 | 53.2 | 53.1 | 52.8 | 51.7 | 52.5 | 52.7 | 52.6 | 51.8 | 53.2 | 53.2 | 52.5 | 52.6 | 51.4 | 52.6 | 52.6 | 52.5 | 52.9 | 52.6 | 53.1 | 52.6 | 52.6 | 51.5 | 52.6 | 59.2 | 73.8 | 100.0 | 79.4 | 80.3 | 75.5 |
| | 35 | *Thermodesulfovibrio yellowstonii* DSM 11347 | 52.9 | 52.9 | 53.1 | 52.8 | 52.8 | 52.4 | 53.1 | 52.9 | 53.0 | 52.9 | 52.7 | 51.9 | 52.9 | 52.4 | 52.4 | 51.7 | 53.2 | 53.3 | 52.5 | 52.6 | 51.4 | 52.7 | 52.6 | 52.6 | 52.6 | 52.2 | 52.9 | 52.3 | 52.7 | 51.2 | 52.7 | 59.1 | 74.9 | 76.8 | 100.0 | 84.2 | 78.9 |
| | 36 | *Thermodesulfovibrio aggregans* TGE-P1 | 53.4 | 53.3 | 53.6 | 53.3 | 53.3 | 52.8 | 53.6 | 53.2 | 53.2 | 53.2 | 53.1 | 51.9 | 53.1 | 52.9 | 52.9 | 52.2 | 53.2 | 53.3 | 52.8 | 53.0 | 51.5 | 53.0 | 53.0 | 53.0 | 53.1 | 52.7 | 53.3 | 52.9 | 53.1 | 51.6 | 52.9 | 59.0 | 74.2 | 77.2 | 80.4 | 100.0 | 78.8 |
| | 37 | *Thermodesulfovibrio thiophilus* DSM 17215 | 52.6 | 52.6 | 52.8 | 52.5 | 52.6 | 52.0 | 52.6 | 52.5 | 52.6 | 52.5 | 51.9 | 51.0 | 52.1 | 52.0 | 51.9 | 51.4 | 52.3 | 52.5 | 52.1 | 52.3 | 51.0 | 52.3 | 52.3 | 52.2 | 52.6 | 52.0 | 52.7 | 51.9 | 52.4 | 51.2 | 52.2 | 58.6 | 72.6 | 74.6 | 77.7 | 76.3 | 100.0 |

**Supplementary table S5.** ANI and dDDH values between LBB01 and closely related genomes.

| Query genome | Reference genome | ANI, % | dDDH, % |
|---|---|---|---|
| *Ca.* Magnetomonas plexicatena LBB01 | *Ca.* Magnetomonas sp. nDJH13bin19 | 83.1 | 27.3 |
| *Ca.* Magnetomonas plexicatena LBB01 | *Ca.* Magnetomonas sp. nDJH14bin5 | 83.1 | 27.3 |
| *Ca.* Magnetomonas plexicatena LBB01 | *Ca.* Magnetomonas sp. nDJH8bin8 | 83.0 | 27.3 |
| *Ca.* Magnetomonas plexicatena LBB01 | *Ca.* Magnetomonas sp. nDJH6bin1 | 82.7 | 27.3 |
| *Ca.* Magnetomonas plexicatena LBB01 | *Ca.* Magnetominusculus xianensis nHCHbin2 | 76.8 | 19.4 |
| *Ca.* Magnetomonas plexicatena LBB01 | *Ca.* Magnetominusculus xianensis HCH-1 | 76.7 | 20.0 |
| *Ca.* Magnetomonas plexicatena LBB01 | *Ca.* Magnetominusculus sp. nDJH8bin6 | 76.5 | 27.5 |
| *Ca.* Magnetomonas plexicatena LBB01 | *Ca.* Magnetominusculus xianensis HCHbin1 | 76.4 | 18.6 |
| *Ca.* Magnetomonas plexicatena LBB01 | *Ca.* Magnetominusculus sp. nMYbin6 | 76.1 | 18.9 |
| *Ca.* Magnetomonas plexicatena LBB01 | *Ca.* Magnetominusculus sp. MYbin6 | 76.0 | 19.8 |
| *Ca.* Magnetomonas plexicatena LBB01 | *Ca.* Magnetominusculus linsii LBB02 | <75.0 | 17.1 |
| *Ca.* Magnetomonas plexicatena LBB01 | *Ca.* Magnetominusculus sp. nDJH14bin7 | <75.0 | 24.3 |
| *Ca.* Magnetomonas plexicatena LBB01 | *Ca.* Magnetominusculus sp. nDJH5bin4 | <75.0 | 23.3 |
| *Ca.* Magnetomonas plexicatena LBB01 | *Ca.* Magnetominusculus sp. nDJH13bin15 | <75.0 | 18.6 |
| *Ca.* Magnetomonas plexicatena LBB01 | *Ca.* Magnetominusculus sp. nDJH8bin13 | <75.0 | 17.9 |
| *Ca.* Magnetomonas plexicatena LBB01 | *Ca.* Magnetobacterium sp. MYbin2 | <75.0 | 36.5 |
| *Ca.* Magnetomonas plexicatena LBB01 | *Ca.* Magnetobacterium casensis MYR-1 | <75.0 | 31.0 |
| *Ca.* Magnetomonas plexicatena LBB01 | *Ca.* Magnetobacterium sp. nMYbin2 | <75.0 | 29.8 |
| *Ca.* Magnetomonas plexicatena LBB01 | *Ca.* Magnetobacterium sp. nDJH15bin2 | <75.0 | 28.8 |
| *Ca.* Magnetomonas plexicatena LBB01 | *Ca.* Magnetobacterium sp. nDC0425bin1 | <75.0 | 26.7 |
| *Ca.* Magnetomonas plexicatena LBB01 | *Ca.* Magnetobacterium sp. nDJH13bin1 | <75.0 | 26.7 |
| *Ca.* Magnetomonas plexicatena LBB01 | *Ca.* Magnetobacterium cryptolimnobacter XYR | <75.0 | 26.3 |
| *Ca.* Magnetomonas plexicatena LBB01 | *Ca.* Magnetobacterium sp. DC0425bin1 | <75.0 | 26.0 |
| *Ca.* Magnetomonas plexicatena LBB01 | *Ca.* Magnetobacterium sp. nMYbin4 | <75.0 | 20.9 |
| *Ca.* Magnetomonas plexicatena LBB01 | *Ca.* Magnetobacterium sp. MYbinv3 | <75.0 | 19.5 |
| *Ca.* Magnetomonas plexicatena LBB01 | *Ca.* Magnetobacterium sp. nMYbin3 | <75.0 | 16.5 |

**Supplementary table S6.** ANI and dDDH values between LBB02 and closely related genomes.

| Query genome | Reference genome | ANI, % | dDDH, % |
|---|---|---|---|
| *Ca*. Magnetominusculus linsii LBB02 | *Ca*. Magnetominusculus sp. nDJH8bin6 | 79.1 | 20.8 |
| *Ca*. Magnetominusculus linsii LBB02 | *Ca*. Magnetominusculus sp. nDJH13bin15 | 78.9 | 20.9 |
| *Ca*. Magnetominusculus linsii LBB02 | *Ca*. Magnetominusculus sp. nDJH14bin7 | 78.8 | 21.2 |
| *Ca*. Magnetominusculus linsii LBB02 | *Ca*. Magnetominusculus xianensis HCH-1 | 78.6 | 20.1 |
| *Ca*. Magnetominusculus linsii LBB02 | *Ca*. Magnetominusculus xianensis HCHbin1 | 78.6 | 20.1 |
| *Ca*. Magnetominusculus linsii LBB02 | *Ca*. Magnetominusculus xianensis nHCHbin2 | 78.5 | 20 |
| *Ca*. Magnetominusculus linsii LBB02 | *Ca*. Magnetominusculus sp. nDJH8bin13 | 78.5 | 20.3 |
| *Ca*. Magnetominusculus linsii LBB02 | *Ca*. Magnetominusculus sp. nDJH5bin4 | 78.0 | 19.8 |
| *Ca*. Magnetominusculus linsii LBB02 | *Ca*. Magnetominusculus sp. nMYbin6 | 77.3 | 18.9 |
| *Ca*. Magnetominusculus linsii LBB02 | *Ca*. Magnetominusculus sp. MYbin6 | 77.3 | 19.1 |
| *Ca*. Magnetominusculus linsii LBB02 | *Ca*. Magnetomonas sp. nDJH8bin8 | 76.2 | 18.5 |
| *Ca*. Magnetominusculus linsii LBB02 | *Ca*. Magnetomonas sp. nDJH14bin5 | 75.7 | 16.2 |
| *Ca*. Magnetominusculus linsii LBB02 | *Ca*. Magnetomonas sp. nDJH13bin19 | 75.7 | 16.3 |
| *Ca*. Magnetominusculus linsii LBB02 | *Ca*. Magnetomonas plexicatena LBB01 | <75.0 | 17.1 |
| *Ca*. Magnetominusculus linsii LBB02 | *Ca*. Magnetomonas sp. nDJH6bin1 | <75.0 | 17.4 |
| *Ca*. Magnetominusculus linsii LBB02 | *Ca*. Magnetobacterium sp. DC0425bin1 | <75.0 | 17.3 |
| *Ca*. Magnetominusculus linsii LBB02 | *Ca*. Magnetobacterium sp. MYbin2 | <75.0 | 19.4 |
| *Ca*. Magnetominusculus linsii LBB02 | *Ca*. Magnetobacterium sp. MYbinv3 | <75.0 | 19.6 |
| *Ca*. Magnetominusculus linsii LBB02 | *Ca*. Magnetobacterium casensis MYR-1 | <75.0 | 17 |
| *Ca*. Magnetominusculus linsii LBB02 | *Ca*. Magnetobacterium sp. nDC0425bin1 | <75.0 | 17.1 |
| *Ca*. Magnetominusculus linsii LBB02 | *Ca*. Magnetobacterium sp. nDJH13bin1 | <75.0 | 19 |
| *Ca*. Magnetominusculus linsii LBB02 | *Ca*. Magnetobacterium sp. nDJH15bin2 | <75.0 | 15.9 |
| *Ca*. Magnetominusculus linsii LBB02 | *Ca*. Magnetobacterium sp. nMYbin2 | <75.0 | 16.4 |
| *Ca*. Magnetominusculus linsii LBB02 | *Ca*. Magnetobacterium sp. nMYbin3 | <75.0 | 14.7 |
| *Ca*. Magnetominusculus linsii LBB02 | *Ca*. Magnetobacterium sp. nMYbin4 | <75.0 | 17.2 |
| *Ca*. Magnetominusculus linsii LBB02 | *Ca*. Magnetobacterium cryptolimnobacter XYR | <75.0 | 17.3 |

**Supplementary table S7.** Genomes used for the MGC genes search in the *Nitrospirota* phylum

| ID | NCBI Organism Name | GTDB Taxonomy | GTDB species representative | NCBI type material |
|---|---|---|---|---|
| GCF_000284315.1 | Leptospirillum ferrooxidans C2-3 | p_Nitrospirota; c_Leptospirillia; o_Leptospirillales; f_Leptospirillaceae; g_Leptospirillum; s_Leptospirillum ferrooxidans | yes | no |
| GCF_000755505.1 | Leptospirillum ferriphilum | p_Nitrospirota; c_Leptospirillia; o_Leptospirillales; f_Leptospirillaceae; g_Leptospirillum_A; s_Leptospirillum_A ferriphilum | yes | yes |
| GCF_000299235.1 | Leptospirillum ferriphilum ML-04 | p_Nitrospirota; c_Leptospirillia; o_Leptospirillales; f_Leptospirillaceae; g_Leptospirillum_A; s_Leptospirillum_A rubarum | yes | no |
| GCA_002387725.1 | Nitrospirae bacterium UBA4572 | p_Nitrospirota; c_Leptospirillia; o_Leptospirillales; f_Leptospirillaceae; g_UBA4572; s_UBA4572 sp002387725 | yes | no |
| GCA_001803875.1 | Nitrospirae bacterium RIFCSPHIGHO2_01_FULL_66_17 | p_Nitrospirota; c_Nitrospiria; o_2-01-FULL-66-17; f_2-01-FULL-66-17; g_2-01-FULL-66-17; s_2-01-FULL-66-17 sp001803875 | yes | no |
| GCA_001805245.1 | Nitrospirae bacterium RIFCSPLOWO2_02_FULL_62_14 | p_Nitrospirota; c_Nitrospiria; o_Nitrospirales; f_Nitrospiraceae; g_2-02-FULL-62-14; s_2-02-FULL-62-14 sp001805245 | yes | no |
| GCA_001914955.1 | Nitrospirae bacterium 13_2_20CM_2_62_8 | p_Nitrospirota; c_Nitrospiria; o_Nitrospirales; f_Nitrospiraceae; g_40CM-3-62-11; s_40CM-3-62-11 sp001914955 | yes | no |
| GCF_001458695.1 | Candidatus Nitrospira inopinata | p_Nitrospirota; c_Nitrospiria; o_Nitrospirales; f_Nitrospiraceae; g_Nitrospira; s_Nitrospira inopinata | yes | no |
| GCF_001273775.1 | Nitrospira moscoviensis | p_Nitrospirota; c_Nitrospiria; o_Nitrospirales; f_Nitrospiraceae; g_Nitrospira; s_Nitrospira moscoviensis | yes | yes |
| GCF_001458775.1 | Candidatus Nitrospira nitrificans | p_Nitrospirota; c_Nitrospiria; o_Nitrospirales; f_Nitrospiraceae; g_Nitrospira; s_Nitrospira nitrificans | yes | no |
| GCF_001458735.1 | Candidatus Nitrospira nitrosa | p_Nitrospirota; c_Nitrospiria; o_Nitrospirales; f_Nitrospiraceae; g_Nitrospira; s_Nitrospira nitrosa | yes | no |
| GCA_001464735.1 | Nitrospira sp. Ga0074138 | p_Nitrospirota; c_Nitrospiria; o_Nitrospirales; f_Nitrospiraceae; g_Nitrospira; s_Nitrospira sp001464735 | yes | no |
| GCA_002083365.1 | Nitrospira sp. SG-bin1 | p_Nitrospirota; c_Nitrospiria; o_Nitrospirales; f_Nitrospiraceae; g_Nitrospira; s_Nitrospira sp002083365 | yes | no |
| GCA_002083565.1 | Nitrospira sp. ST-bin4 | p_Nitrospirota; c_Nitrospiria; o_Nitrospirales; f_Nitrospiraceae; g_Nitrospira; s_Nitrospira sp002083565 | yes | no |
| GCA_002254365.1 | Nitrospira sp. UW-LDO-01 | p_Nitrospirota; c_Nitrospiria; o_Nitrospirales; f_Nitrospiraceae; g_Nitrospira; s_Nitrospira sp002254365 | yes | no |
| GCA_002331335.1 | Nitrospira sp. UBA2082 | p_Nitrospirota; c_Nitrospiria; o_Nitrospirales; f_Nitrospiraceae; g_Nitrospira; s_Nitrospira sp002331335 | yes | no |
| GCA_002331625.1 | Nitrospira sp. UBA2083 | p_Nitrospirota; c_Nitrospiria; o_Nitrospirales; f_Nitrospiraceae; g_Nitrospira; s_Nitrospira sp002331625 | yes | no |
| GCA_002420115.1 | Nitrospira sp. UBA5698 | p_Nitrospirota; c_Nitrospiria; o_Nitrospirales; f_Nitrospiraceae; g_Nitrospira; s_Nitrospira sp002420115 | yes | no |
| GCA_002451055.1 | Nitrospira sp. UBA6909 | p_Nitrospirota; c_Nitrospiria; o_Nitrospirales; f_Nitrospiraceae; g_Nitrospira; s_Nitrospira sp002451055 | yes | no |
| GCA_002869845.2 | Nitrospira sp. CG24B | p_Nitrospirota; c_Nitrospiria; o_Nitrospirales; f_Nitrospiraceae; g_Nitrospira; s_Nitrospira sp002869845 | yes | no |
| GCA_005116745.1 | Nitrospira sp. | p_Nitrospirota; c_Nitrospiria; o_Nitrospirales; f_Nitrospiraceae; g_Nitrospira; s_Nitrospira sp005116745 | yes | no |
| GCA_005116895.1 | Nitrospira sp. | p_Nitrospirota; c_Nitrospiria; o_Nitrospirales; f_Nitrospiraceae; g_Nitrospira; s_Nitrospira sp005116895 | yes | no |
| GCA_005116955.1 | Nitrospira sp. | p_Nitrospirota; c_Nitrospiria; o_Nitrospirales; f_Nitrospiraceae; g_Nitrospira; s_Nitrospira sp005116955 | yes | no |
| GCA_005116965.1 | Nitrospira sp. | p_Nitrospirota; c_Nitrospiria; o_Nitrospirales; f_Nitrospiraceae; g_Nitrospira; s_Nitrospira sp005116965 | yes | no |
| GCA_005239465.1 | Nitrospira sp. | p_Nitrospirota; c_Nitrospiria; o_Nitrospirales; f_Nitrospiraceae; g_Nitrospira; s_Nitrospira sp005239465 | yes | no |
| GCA_005793285.1 | Nitrospiraceae bacterium | p_Nitrospirota; c_Nitrospiria; o_Nitrospirales; f_Nitrospiraceae; g_Nitrospira; s_Nitrospira sp005793285 | yes | no |
| GCF_000196815.1 | Nitrospira defluvii | p_Nitrospirota; c_Nitrospiria; o_Nitrospirales; f_Nitrospiraceae; g_Nitrospira_A; s_Nitrospira_A defluvii | yes | no |
| GCA_001567445.1 | Nitrospira sp. OLB3 | p_Nitrospirota; c_Nitrospiria; o_Nitrospirales; f_Nitrospiraceae; g_Nitrospira_A; s_Nitrospira_A sp001567445 | yes | no |
| GCA_003456605.1 | Nitrospira sp. | p_Nitrospirota; c_Nitrospiria; o_Nitrospirales; f_Nitrospiraceae; g_Nitrospira_A; s_Nitrospira_A sp003456605 | yes | no |
| GCF_900170025.1 | Nitrospira sp. ND1 | p_Nitrospirota; c_Nitrospiria; o_Nitrospirales; f_Nitrospiraceae; g_Nitrospira_A; s_Nitrospira_A sp900170025 | yes | no |
| GCA_900696505.1 | uncultured Nitrosospira sp. | p_Nitrospirota; c_Nitrospiria; o_Nitrospirales; f_Nitrospiraceae; g_Nitrospira_A; s_Nitrospira_A sp900696505 | yes | no |
| GCF_900169565.1 | Nitrospira japonica | p_Nitrospirota; c_Nitrospiria; o_Nitrospirales; f_Nitrospiraceae; g_Nitrospira_C; s_Nitrospira_C japonica | yes | no |
| GCA_002420105.1 | Nitrospira sp. UBA5699 | p_Nitrospirota; c_Nitrospiria; o_Nitrospirales; f_Nitrospiraceae; g_Nitrospira_C; s_Nitrospira_C sp002420105 | yes | no |
| GCF_900403705.1 | Nitrospira lenta | p_Nitrospirota; c_Nitrospiria; o_Nitrospirales; f_Nitrospiraceae; g_Nitrospira_D; s_Nitrospira_D lenta | yes | no |
| GCA_002083555.1 | Nitrospira sp. ST-bin5 | p_Nitrospirota; c_Nitrospiria; o_Nitrospirales; f_Nitrospiraceae; g_Nitrospira_D; s_Nitrospira_D sp002083555 | yes | no |
| GCA_002435325.1 | Nitrospira sp. UBA6493 | p_Nitrospirota; c_Nitrospiria; o_Nitrospirales; f_Nitrospiraceae; g_Nitrospira_D; s_Nitrospira_D sp002435325 | yes | no |
| GCA_002869855.2 | Nitrospira sp. CG24D | p_Nitrospirota; c_Nitrospiria; o_Nitrospirales; f_Nitrospiraceae; g_Nitrospira_D; s_Nitrospira_D sp002869855 | yes | no |
| GCA_005877775.1 | Nitrospirae bacterium | p_Nitrospirota; c_Nitrospiria; o_Nitrospirales; f_Nitrospiraceae; g_NS-7; s_NS-7 sp005877775 | yes | no |
| GCA_002737345.1 | Nitrospirae bacterium | p_Nitrospirota; c_Nitrospiria; o_Nitrospirales; f_Nitrospiraceae; g_Palsa-1315; s_Palsa-1315 sp002737345 | yes | no |
| GCA_002869885.2 | Nitrospira sp. CG24C | p_Nitrospirota; c_Nitrospiria; o_Nitrospirales; f_Nitrospiraceae; g_Palsa-1315; s_Palsa-1315 sp002869885 | yes | no |
| GCA_002869895.2 | Nitrospira sp. CG24E | p_Nitrospirota; c_Nitrospiria; o_Nitrospirales; f_Nitrospiraceae; g_Palsa-1315; s_Palsa-1315 sp002869895 | yes | no |
| GCA_002869925.2 | Nitrospira sp. CG24A | p_Nitrospirota; c_Nitrospiria; o_Nitrospirales; f_Nitrospiraceae; g_Palsa-1315; s_Palsa-1315 sp002869925 | yes | no |
| GCA_003135435.1 | Nitrospira sp. | p_Nitrospirota; c_Nitrospiria; o_Nitrospirales; f_Nitrospiraceae; g_Palsa-1315; s_Palsa-1315 sp003135435 | yes | no |
| GCA_005116775.1 | Nitrospira sp. | p_Nitrospirota; c_Nitrospiria; o_Nitrospirales; f_Nitrospiraceae; g_Palsa-1315; s_Palsa-1315 sp005116775 | yes | no |
| GCA_005116815.1 | Nitrospira sp. | p_Nitrospirota; c_Nitrospiria; o_Nitrospirales; f_Nitrospiraceae; g_Palsa-1315; s_Palsa-1315 sp005116815 | yes | no |
| GCA_005116835.1 | Nitrospira sp. | p_Nitrospirota; c_Nitrospiria; o_Nitrospirales; f_Nitrospiraceae; g_Palsa-1315; s_Palsa-1315 sp005116835 | yes | no |
| GCA_005116885.1 | Nitrospira sp. | p_Nitrospirota; c_Nitrospiria; o_Nitrospirales; f_Nitrospiraceae; g_Palsa-1315; s_Palsa-1315 sp005116885 | yes | no |
| GCA_005116945.1 | Nitrospira sp. | p_Nitrospirota; c_Nitrospiria; o_Nitrospirales; f_Nitrospiraceae; g_Palsa-1315; s_Palsa-1315 sp005116945 | yes | no |
| GCA_005239475.1 | Nitrospira sp. | p_Nitrospirota; c_Nitrospiria; o_Nitrospirales; f_Nitrospiraceae; g_Palsa-1315; s_Palsa-1315 sp005239475 | yes | no |
| GCA_005800255.1 | Nitrospiraceae bacterium | p_Nitrospirota; c_Nitrospiria; o_Nitrospirales; f_Nitrospiraceae; g_Palsa-1315; s_Palsa-1315 sp005800255 | yes | no |
| GCA_005877535.1 | Nitrospirae bacterium | p_Nitrospirota; c_Nitrospiria; o_Nitrospirales; f_Nitrospiraceae; g_Palsa-1315; s_Palsa-1315 sp005877535 | yes | no |
| GCA_005887945.1 | Nitrospirae bacterium | p_Nitrospirota; c_Nitrospiria; o_Nitrospirales; f_Nitrospiraceae; g_Palsa-1315; s_Palsa-1315 sp005887945 | yes | no |
| GCA_005116795.1 | Nitrospira sp. | p_Nitrospirota; c_Nitrospiria; o_Nitrospirales; f_Nitrospiraceae; g_RSF151; s_RSF151 sp005116795 | yes | no |
| GCA_004296865.1 | Nitrospirae bacterium | p_Nitrospirota; c_Nitrospiria; o_Nitrospirales; f_Nitrospiraceae; g_SCVQ01; s_SCVQ01 sp004296865 | yes | no |
| GCA_004296885.1 | Nitrospirae bacterium | p_Nitrospirota; c_Nitrospiria; o_Nitrospirales; f_Nitrospiraceae; g_SYGV01; s_SYGV01 sp004296885 | yes | no |
| GCA_005799365.1 | Nitrospiraceae bacterium | p_Nitrospirota; c_Nitrospiria; o_Nitrospirales; f_Nitrospiraceae; g_SYGV01; s_SYGV01 sp005799365 | yes | no |
| GCA_005877525.1 | Nitrospirae bacterium | p_Nitrospirota; c_Nitrospiria; o_Nitrospirales; f_NS-4; g_NS-11; s_NS-11 sp005877525 | yes | no |
| GCA_005877505.1 | Nitrospirae bacterium | p_Nitrospirota; c_Nitrospiria; o_Nitrospirales; f_NS-4; g_NS-12; s_NS-12 sp005877505 | yes | no |
| GCA_005877565.1 | Nitrospirae bacterium | p_Nitrospirota; c_Nitrospiria; o_Nitrospirales; f_NS-4; g_NS-12; s_NS-12 sp005877565 | yes | no |
| GCA_005877615.1 | Nitrospirae bacterium | p_Nitrospirota; c_Nitrospiria; o_Nitrospirales; f_NS-4; g_NS-12; s_NS-12 sp005877615 | yes | no |

| ID | NCBI Organism Name | GTDB Taxonomy | GTDB species representative | NCBI type material |
|---|---|---|---|---|
| GCA_005877855.1 | Nitrospirae bacterium | p_Nitrospirota; c_Nitrospiria; o_Nitrospirales; f_NS-4; g_NS-12; s_NS-12 sp005877855 | yes | no |
| GCA_005877815.1 | Nitrospirae bacterium | p_Nitrospirota; c_Nitrospiria; o_Nitrospirales; f_NS-4; g_NS-4; s_NS-4 sp005877815 | yes | no |
| GCA_004298335.1 | Nitrospirae bacterium | p_Nitrospirota; c_Nitrospiria; o_Nitrospirales; f_NS-4; g_SCTG01; s_SCTG01 sp004298335 | yes | no |
| GCA_002328825.1 | Nitrospiraceae bacterium UBA2166 | p_Nitrospirota; c_Nitrospiria; o_Nitrospirales; f_UBA2166; g_UBA2166; s_UBA2166 sp002328825 | yes | no |
| GCA_002726235.1 | Nitrospiraceae bacterium | p_Nitrospirota; c_Nitrospiria; o_Nitrospirales; f_UBA2166; g_UBA2166; s_UBA2166 sp002726235 | yes | no |
| GCA_002238765.1 | Nitrospira sp. bin75 | p_Nitrospirota; c_Nitrospiria; o_Nitrospirales; f_UBA8639; g_Bin75; s_Bin75 sp002238765 | yes | no |
| GCA_003695915.1 | Nitrospirae bacterium | p_Nitrospirota; c_Nitrospiria; o_Nitrospirales; f_UBA8639; g_J017; s_J017 sp003695915 | yes | no |
| GCA_003696975.1 | Nitrospirae bacterium | p_Nitrospirota; c_Nitrospiria; o_Nitrospirales; f_UBA8639; g_J031; s_J031 sp003696975 | yes | no |
| GCA_001643555.1 | Nitrospirae bacterium SPGG5 | p_Nitrospirota; c_Nitrospiria; o_Nitrospirales; f_UBA8639; g_SPGG5; s_SPGG5 sp001643555 | yes | no |
| GCA_003228495.1 | Nitrospirae bacterium | p_Nitrospirota; c_Nitrospiria; o_Nitrospirales; f_UBA8639; g_UBA8639; s_UBA8639 sp003228495 | yes | no |
| GCA_003233615.1 | Nitrospirae bacterium | p_Nitrospirota; c_Nitrospiria; o_Nitrospirales; f_UBA8639; g_UBA8639; s_UBA8639 sp003233615 | yes | no |
| GCA_003235785.1 | Nitrospirae bacterium | p_Nitrospirota; c_Nitrospiria; o_Nitrospirales; f_UBA8639; g_UBA8639; s_UBA8639 sp003235785 | yes | no |
| GCA_003523945.1 | Nitrospiraceae bacterium | p_Nitrospirota; c_Nitrospiria; o_Nitrospirales; f_UBA8639; g_UBA8639; s_UBA8639 sp003523945 | yes | no |
| GCA_005239925.1 | Nitrospira sp. | p_Nitrospirota; c_Nitrospiria; o_SBBL01; f_SBBA01; g_SBBA01; s_SBBA01 sp005239925 | yes | no |
| GCA_005239745.1 | Nitrospira sp. | p_Nitrospirota; c_Nitrospiria; o_SBBL01; f_SBBL01; g_SBBL01; s_SBBL01 sp005239745 | yes | no |
| GCA_005239595.1 | Nitrospira sp. | p_Nitrospirota; c_Nitrospiria; o_SBBL01; f_SBBT01; g_SBBT01; s_SBBT01 sp005239595 | yes | no |
| GCA_004297235.1 | Nitrospirae bacterium | p_Nitrospirota; c_Nitrospiria; o_SBBL01; f_SCUR01; g_SCUR01; s_SCUR01 sp004297235 | yes | no |
| GCA_001803815.1 | Nitrospirae bacterium RBG_19FT_COMBO_42_15 | p_Nitrospirota; c_Nitrospiria_A; o_9FT-COMBO-42-15; f_9FT-COMBO-42-15; g_9FT-COMBO-42-15; s_9FT-COMBO-42-15 sp001803815 | yes | no |
| GCA_001805205.1 | Nitrospirae bacterium RIFCSPLOW2_12_42_9 | p_Nitrospirota; c_Nitrospiria_A; o_HDB-SIOI813; f_HDB-SIOI813; g_HDB-SIOI813; s_HDB-SIOI813 sp001805205 | yes | no |
| GCA_001805165.1 | Nitrospirae bacterium RBG_16_43_11 | p_Nitrospirota; c_Nitrospiria_A; o_HDB-SIOI813; f_HDB-SIOI813; g_RBG-16-43-11; s_RBG-16-43-11 sp001805165 | yes | no |
| GCA_001803795.1 | Nitrospirae bacterium RBG_16_64_22 | p_Nitrospirota; c_RBG-16-64-22; o_RBG-16-64-22; f_RBG-16-64-22; g_RBG-16-64-22; s_RBG-16-64-22 sp001803795 | yes | no |
| GCA_003235715.1 | Nitrospirae bacterium | p_Nitrospirota; c_Thermodesulfovibrionia; o_SZUA-242; f_SZUA-242; g_SZUA-242; s_SZUA-242 sp003235715 | yes | no |
| GCA_002897855.1 | bacterium BMS3Bbin05 | p_Nitrospirota; c_Thermodesulfovibrionia; o_Thermodesulfovibrionales; f_BMS3Bbin05; g_BMS3Bbin05; s_BMS3Bbin05 sp002897855 | yes | no |
| GCA_002897935.1 | bacterium BMS3Abin08 | p_Nitrospirota; c_Thermodesulfovibrionia; o_Thermodesulfovibrionales; f_JdFR-85; g_BMS3Abin08; s_BMS3Abin08 sp002897935 | yes | no |
| GCA_002898135.1 | bacterium BMS3Bbin07 | p_Nitrospirota; c_Thermodesulfovibrionia; o_Thermodesulfovibrionales; f_JdFR-85; g_BMS3Bbin07; s_BMS3Bbin07 sp002898135 | yes | no |
| GCA_003232715.1 | Nitrospirae bacterium | p_Nitrospirota; c_Thermodesulfovibrionia; o_Thermodesulfovibrionales; f_JdFR-85; g_BMS3Bbin07; s_BMS3Bbin07 sp003232715 | yes | no |
| GCA_002011745.1 | Nitrospirae bacterium JdFR-85 | p_Nitrospirota; c_Thermodesulfovibrionia; o_Thermodesulfovibrionales; f_JdFR-85; g_JdFR-85; s_JdFR-85 sp002011745 | yes | no |
| GCA_002011815.1 | Nitrospirae bacterium JdFR-86 | p_Nitrospirota; c_Thermodesulfovibrionia; o_Thermodesulfovibrionales; f_JdFR-86; g_JdFR-86; s_JdFR-86 sp002011815 | yes | no |
| GCA_002011795.1 | Nitrospirae bacterium JdFR-88 | p_Nitrospirota; c_Thermodesulfovibrionia; o_Thermodesulfovibrionales; f_JdFR-88; g_JdFR-88; s_JdFR-88 sp002011795 | yes | no |
| GCF_001541255.1 | Nitrospirae bacterium HCH-1 | p_Nitrospirota; c_Thermodesulfovibrionia; o_Thermodesulfovibrionales; f_Magnetobacteriaceae; g_HCH-1; s_HCH-1 sp001541255 | yes | no |
| GCA_002753305.1 | Nitrospirae bacterium MYbin6 | p_Nitrospirota; c_Thermodesulfovibrionia; o_Thermodesulfovibrionales; f_Magnetobacteriaceae; g_HCH-1; s_HCH-1 sp002753305 | yes | no |
| GCF_000714715.1 | Candidatus Magnetobacterium casensis | p_Nitrospirota; c_Thermodesulfovibrionia; o_Thermodesulfovibrionales; f_Magnetobacteriaceae; g_Magnetobacterium; s_Magnetobacterium casensis | yes | no |
| GCA_002753395.1 | Nitrospirae bacterium MYbinv3 | p_Nitrospirota; c_Thermodesulfovibrionia; o_Thermodesulfovibrionales; f_Magnetobacteriaceae; g_Magnetobacterium; s_Magnetobacterium sp002753395 | yes | no |
| GCA_002753685.1 | Nitrospirae bacterium | p_Nitrospirota; c_Thermodesulfovibrionia; o_Thermodesulfovibrionales; f_Magnetobacteriaceae; g_Magnetobacterium; s_Magnetobacterium sp002753685 | yes | no |
| GCA_002780895.1 | Nitrospirae bacterium CG02_land_8_20_14_3_00_41_53 | p_Nitrospirota; c_Thermodesulfovibrionia; o_Thermodesulfovibrionales; f_SM23-35; g_0-14-3-00-41-53; s_0-14-3-00-41-53 sp002780895 | yes | no |
| GCA_001805125.1 | Nitrospirae bacterium RBG_13_39_12 | p_Nitrospirota; c_Thermodesulfovibrionia; o_Thermodesulfovibrionales; f_SM23-35; g_RBG-13-39-12; s_RBG-13-39-12 sp001805125 | yes | no |
| GCA_002328665.1 | Nitrospiraceae bacterium UBA2194 | p_Nitrospirota; c_Thermodesulfovibrionia; o_Thermodesulfovibrionales; f_SM23-35; g_UBA2194; s_UBA2194 sp002328665 | yes | no |
| GCF_001514535.1 | Thermodesulfovibrio aggregans | p_Nitrospirota; c_Thermodesulfovibrionia; o_Thermodesulfovibrionales; f_Thermodesulfovibrionaceae; g_Thermodesulfovibrio; s_aggregans | yes | yes |
| GCA_002878055.1 | Thermodesulfovibrio aggregans | p_Nitrospirota; c_Thermodesulfovibrionia; o_Thermodesulfovibrionales; f_Thermodesulfovibrionaceae; g_Thermodesulfovibrio; s_aggregans_A | yes | no |
| GCF_001707915.1 | Thermodesulfovibrio sp. N1 | p_Nitrospirota; c_Thermodesulfovibrionia; o_Thermodesulfovibrionales; f_Thermodesulfovibrionaceae; g_Thermodesulfovibrio; s_sp001707915 | yes | no |
| GCA_002339325.1 | Nitrospiraceae bacterium UBA2600 | p_Nitrospirota; c_Thermodesulfovibrionia; o_Thermodesulfovibrionales; f_Thermodesulfovibrionaceae; g_Thermodesulfovibrio; s_sp002339325 | yes | no |
| GCF_000423865.1 | Thermodesulfovibrio thiophilus DSM 17215 | p_Nitrospirota; c_Thermodesulfovibrionia; o_Thermodesulfovibrionales; f_Thermodesulfovibrionaceae; g_Thermodesulfovibrio; s_thiophilus | yes | yes |
| GCF_000020985.1 | Thermodesulfovibrio yellowstonii DSM 11347 | p_Nitrospirota; c_Thermodesulfovibrionia; o_Thermodesulfovibrionales; f_Thermodesulfovibrionaceae; g_Thermodesulfovibrio; s_yellowstonii | yes | yes |
| GCA_004298625.1 | Nitrospirae bacterium | p_Nitrospirota; c_Thermodesulfovibrionia; o_Thermodesulfovibrionales; f_UBA1546; g_SCSY01; s_SCSY01 sp004298625 | yes | no |
| GCA_001805105.1 | Nitrospirae bacterium GWF2_44_13 | p_Nitrospirota; c_Thermodesulfovibrionia; o_Thermodesulfovibrionales; f_UBA1546; g_UBA1546; s_UBA1546 sp001805105 | yes | no |
| GCA_001871685.1 | Nitrospirae bacterium CG1_02_44_142 | p_Nitrospirota; c_Thermodesulfovibrionia; o_Thermodesulfovibrionales; f_UBA1546; g_UBA1546; s_UBA1546 sp001871685 | yes | no |
| GCA_003535475.1 | Nitrospiraceae bacterium | p_Nitrospirota; c_Thermodesulfovibrionia; o_Thermodesulfovibrionales; f_UBA1546; g_UBA1546; s_UBA1546 sp003535475 | yes | no |
| GCA_003599795.1 | Nitrospiraceae bacterium | p_Nitrospirota; c_Thermodesulfovibrionia; o_Thermodesulfovibrionales; f_UBA1546; g_UBA1546; s_UBA1546 sp003599795 | yes | no |
| GCA_001803645.1 | Nitrospirae bacterium GWC2_42_7 | p_Nitrospirota; c_Thermodesulfovibrionia; o_Thermodesulfovibrionales; f_UBA6898; g_GWC2-42-7; s_GWC2-42-7 sp001803645 | yes | no |
| GCA_002839535.1 | Nitrospira bacterium HGW-Nitrospira-1 | p_Nitrospirota; c_Thermodesulfovibrionia; o_Thermodesulfovibrionales; f_UBA6898; g_GW-Nitrospira-1; s_GW-Nitrospira-1 sp002839535 | yes | no |
| GCA_003153275.1 | Nitrospiraceae bacterium | p_Nitrospirota; c_Thermodesulfovibrionia; o_Thermodesulfovibrionales; f_UBA6898; g_PALSA-1316; s_PALSA-1316 sp003153275 | yes | no |
| GCA_004321925.1 | Nitrospirae bacterium | p_Nitrospirota; c_Thermodesulfovibrionia; o_Thermodesulfovibrionales; f_UBA6898; g_PALSA-1316; s_PALSA-1316 sp004321925 | yes | no |
| GCF_900302705.1 | Candidatus Sulfobium mesophilum | p_Nitrospirota; c_Thermodesulfovibrionia; o_Thermodesulfovibrionales; f_UBA6898; g_Sulfobium; s_Sulfobium mesophilum | yes | no |
| GCA_002448975.1 | Nitrospiraceae bacterium UBA6898 | p_Nitrospirota; c_Thermodesulfovibrionia; o_Thermodesulfovibrionales; f_UBA6898; g_UBA6898; s_UBA6898 sp002448975 | yes | no |
| GCA_002451165.1 | Nitrospiraceae bacterium UBA6905 | p_Nitrospirota; c_Thermodesulfovibrionia; o_Thermodesulfovibrionales; f_UBA6898; g_UBA6898; s_UBA6898 sp002451165 | yes | no |
| GCA_003252075.1 | Nitrospirae bacterium | p_Nitrospirota; c_Thermodesulfovibrionia; o_Thermodesulfovibrionales; f_UBA6898; g_UBA6898; s_UBA6898 sp003252075 | yes | no |
| GCA_003453735.1 | Nitrospiraceae bacterium | p_Nitrospirota; c_Thermodesulfovibrionia; o_Thermodesulfovibrionales; f_UBA9159; g_UBA9159; s_UBA9159 sp003453735 | yes | no |
| GCA_003170655.1 | Nitrospiraceae bacterium | p_Nitrospirota; c_Thermodesulfovibrionia; o_Thermodesulfovibrionales; f_UBA9935; g_Fen-1308; s_Fen-1308 sp003170655 | yes | no |
| GCA_002634385.1 | Nitrospira sp. | p_Nitrospirota; c_Thermodesulfovibrionia; o_Thermodesulfovibrionales; f_UBA9935; g_GCA-2634385; s_GCA-2634385 sp002634385 | yes | no |
| GCA_001803635.1 | Nitrospirae bacterium GWB2_47_37 | p_Nitrospirota; c_Thermodesulfovibrionia; o_Thermodesulfovibrionales; f_UBA9935; g_GWB2-47-37; s_GWB2-47-37 sp001803635 | yes | no |
| GCA_002753335.1 | Nitrospirae bacterium MYbin3 | p_Nitrospirota; c_Thermodesulfovibrionia; o_Thermodesulfovibrionales; f_UBA9935; g_MYbin3; s_MYbin3 sp002753335 | yes | no |
| GCA_004321915.1 | Nitrospirae bacterium | p_Nitrospirota; c_Thermodesulfovibrionia; o_Thermodesulfovibrionales; f_UBA9935; g_MYbin3; s_MYbin3 sp004321915 | yes | no |

| ID | NCBI Organism Name | GTDB Taxonomy | GTDB species representative | NCBI type material |
|---|---|---|---|---|
| GCA_002299835.1 | Nitrospiraceae bacterium UBA665 | p_Nitrospirota; c_Thermodesulfovibrionia; o_Thermodesulfovibrionales; f_UBA9935; g_UBA665; s_UBA665 sp002299835 | yes | no |
| GCA_002897775.1 | bacterium BMS3Bbin08 | p_Nitrospirota; c_Thermodesulfovibrionia; o_UBA6902; f_BMS3Bbin08; g_BMS3Bbin08; s_BMS3Bbin08 sp002897775 | yes | no |
| GCA_002011735.1 | Nitrospirae bacterium JdFR-81 | p_Nitrospirota; c_Thermodesulfovibrionia; o_UBA6902; f_JdFR-81; g_JdFR-81; s_JdFR-81 sp002011735 | yes | no |
| GCA_002897895.1 | bacterium BMS3Abin06 | p_Nitrospirota; c_Thermodesulfovibrionia; o_UBA6902; f_UBA6902; g_BMS3ABIN06; s_BMS3ABIN06 sp002897895 | yes | no |
| GCA_002897915.1 | bacterium BMS3Bbin09 | p_Nitrospirota; c_Thermodesulfovibrionia; o_UBA6902; f_UBA6902; g_BMS3Abin09; s_BMS3Abin09 sp002897915 | yes | no |
| GCA_003354025.1 | Nitrospirae bacterium | p_Nitrospirota; c_Thermodesulfovibrionia; o_UBA6902; f_UBA6902; g_Glo-13; s_Glo-13 sp003354025 | yes | no |
| GCA_003599505.1 | Nitrospiraceae bacterium | p_Nitrospirota; c_Thermodesulfovibrionia; o_UBA6902; f_UBA6902; g_SURF-11; s_SURF-11 sp003599505 | yes | no |
| GCA_003599425.1 | Nitrospiraceae bacterium | p_Nitrospirota; c_Thermodesulfovibrionia; o_UBA6902; f_UBA6902; g_SURF-23; s_SURF-23 sp003599425 | yes | no |
| GCA_003599275.1 | Nitrospiraceae bacterium | p_Nitrospirota; c_Thermodesulfovibrionia; o_UBA6902; f_UBA6902; g_SURF-45; s_SURF-45 sp003599275 | yes | no |
| GCA_002451135.1 | Nitrospiraceae bacterium UBA6902 | p_Nitrospirota; c_Thermodesulfovibrionia; o_UBA6902; f_UBA6902; g_UBA6902; s_UBA6902 sp002451135 | yes | no |
| GCA_003599025.1 | Nitrospiraceae bacterium | p_Nitrospirota; c_Thermodesulfovibrionia; o_UBA6902; f_UBA6902; g_UBA6902; s_UBA6902 sp003599025 | yes | no |
| GCA_001803705.1 | Nitrospirae bacterium GWC2_56_14 | p_Nitrospirota; c_UBA9217; o_UBA9217; f_UBA9217; g_GWC2-56-14; s_GWC2-56-14 sp001803705 | yes | no |
| GCA_001805055.1 | Nitrospirae bacterium GWC2_57_13 | p_Nitrospirota; c_UBA9217; o_UBA9217; f_UBA9217; g_GWC2-57-13; s_GWC2-57-13 sp001805055 | yes | no |
| GCA_003454665.1 | Nitrospiraceae bacterium | p_Nitrospirota; c_UBA9217; o_UBA9217; f_UBA9217; g_UBA9217; s_UBA9217 sp003454665 | yes | no |
| GCF_900198525.1 | Leptospirillum ferriphilum | p_Nitrospirota; c_Leptospirillia; o_Leptospirillales; f_Leptospirillaceae; g_Leptospirillum_A; s_Leptospirillum_A ferriphilum | no | no |
| GCA_000205145.2 | Leptospirillum rubarum | p_Nitrospirota; c_Leptospirillia; o_Leptospirillales; f_Leptospirillaceae; g_Leptospirillum_A; s_Leptospirillum_A rubarum | no | no |
| GCA_000262365.1 | Leptospirillum sp. Group II 'C75' | p_Nitrospirota; c_Leptospirillia; o_Leptospirillales; f_Leptospirillaceae; g_Leptospirillum_A; s_Leptospirillum_A rubarum | no | no |
| GCA_002386985.1 | Leptospirillum ferriphilum | p_Nitrospirota; c_Leptospirillia; o_Leptospirillales; f_Leptospirillaceae; g_Leptospirillum_A; s_Leptospirillum_A rubarum | no | no |
| GCA_002420195.1 | Leptospirillum ferriphilum | p_Nitrospirota; c_Leptospirillia; o_Leptospirillales; f_Leptospirillaceae; g_Leptospirillum_A; s_Leptospirillum_A rubarum | no | no |
| GCA_002455015.1 | Leptospirillum ferriphilum | p_Nitrospirota; c_Leptospirillia; o_Leptospirillales; f_Leptospirillaceae; g_Leptospirillum_A; s_Leptospirillum_A rubarum | no | no |
| GCA_002455035.1 | Leptospirillum ferriphilum | p_Nitrospirota; c_Leptospirillia; o_Leptospirillales; f_Leptospirillaceae; g_Leptospirillum_A; s_Leptospirillum_A rubarum | no | no |
| GCA_002455075.1 | Leptospirillum ferriphilum | p_Nitrospirota; c_Leptospirillia; o_Leptospirillales; f_Leptospirillaceae; g_Leptospirillum_A; s_Leptospirillum_A rubarum | no | no |
| GCA_002455095.1 | Leptospirillum ferriphilum | p_Nitrospirota; c_Leptospirillia; o_Leptospirillales; f_Leptospirillaceae; g_Leptospirillum_A; s_Leptospirillum_A rubarum | no | no |
| GCA_002455445.1 | Leptospirillum ferriphilum | p_Nitrospirota; c_Leptospirillia; o_Leptospirillales; f_Leptospirillaceae; g_Leptospirillum_A; s_Leptospirillum_A rubarum | no | no |
| GCA_002469705.1 | Leptospirillum ferriphilum | p_Nitrospirota; c_Leptospirillia; o_Leptospirillales; f_Leptospirillaceae; g_Leptospirillum_A; s_Leptospirillum_A rubarum | no | no |
| GCA_002470195.1 | Leptospirillum ferriphilum | p_Nitrospirota; c_Leptospirillia; o_Leptospirillales; f_Leptospirillaceae; g_Leptospirillum_A; s_Leptospirillum_A rubarum | no | no |
| GCA_002470665.1 | Leptospirillum ferriphilum | p_Nitrospirota; c_Leptospirillia; o_Leptospirillales; f_Leptospirillaceae; g_Leptospirillum_A; s_Leptospirillum_A rubarum | no | no |
| GCA_002500075.1 | Leptospirillum ferriphilum | p_Nitrospirota; c_Leptospirillia; o_Leptospirillales; f_Leptospirillaceae; g_Leptospirillum_A; s_Leptospirillum_A rubarum | no | no |
| GCA_002500715.1 | Leptospirillum ferriphilum | p_Nitrospirota; c_Leptospirillia; o_Leptospirillales; f_Leptospirillaceae; g_Leptospirillum_A; s_Leptospirillum_A rubarum | no | no |
| GCF_000695975.1 | Leptospirillum ferriphilum YSK | p_Nitrospirota; c_Leptospirillia; o_Leptospirillales; f_Leptospirillaceae; g_Leptospirillum_A; s_Leptospirillum_A rubarum | no | no |
| GCF_001186405.1 | Leptospirillum sp. Group II 'CF-1' | p_Nitrospirota; c_Leptospirillia; o_Leptospirillales; f_Leptospirillaceae; g_Leptospirillum_A; s_Leptospirillum_A rubarum | no | no |
| GCF_001280545.1 | Leptospirillum ferriphilum | p_Nitrospirota; c_Leptospirillia; o_Leptospirillales; f_Leptospirillaceae; g_Leptospirillum_A; s_Leptospirillum_A rubarum | no | no |
| GCF_002002505.1 | Leptospirillum ferriphilum | p_Nitrospirota; c_Leptospirillia; o_Leptospirillales; f_Leptospirillaceae; g_Leptospirillum_A; s_Leptospirillum_A rubarum | no | no |
| GCF_002002665.1 | Leptospirillum ferriphilum | p_Nitrospirota; c_Leptospirillia; o_Leptospirillales; f_Leptospirillaceae; g_Leptospirillum_A; s_Leptospirillum_A rubarum | no | no |
| GCA_002386965.1 | Nitrospirae bacterium UBA4574 | p_Nitrospirota; c_Leptospirillia; o_Leptospirillales; f_Leptospirillaceae; g_UBA4572; s_UBA4572 sp002387725 | no | no |
| GCA_002420165.1 | Nitrospirae bacterium UBA5696 | p_Nitrospirota; c_Leptospirillia; o_Leptospirillales; f_Leptospirillaceae; g_UBA4572; s_UBA4572 sp002387725 | no | no |
| GCA_002454995.1 | Nitrospirae bacterium UBA6678 | p_Nitrospirota; c_Leptospirillia; o_Leptospirillales; f_Leptospirillaceae; g_UBA4572; s_UBA4572 sp002387725 | no | no |
| GCA_002455055.1 | Nitrospirae bacterium UBA6675 | p_Nitrospirota; c_Leptospirillia; o_Leptospirillales; f_Leptospirillaceae; g_UBA4572; s_UBA4572 sp002387725 | no | no |
| GCA_002470685.1 | bacterium UBA7390 | p_Nitrospirota; c_Leptospirillia; o_Leptospirillales; f_Leptospirillaceae; g_UBA4572; s_UBA4572 sp002387725 | no | no |
| GCA_002500645.1 | Nitrospirae bacterium UBA7871 | p_Nitrospirota; c_Leptospirillia; o_Leptospirillales; f_Leptospirillaceae; g_UBA4572; s_UBA4572 sp002387725 | no | no |
| GCA_001805225.1 | Nitrospirae bacterium RIFCSPLOWO2_01_FULL_62_17 | p_Nitrospirota; c_Nitrospiria; o_Nitrospirales; f_Nitrospiraceae; g_2-02-FULL-62-14; s_2-02-FULL-62-14 sp001805245 | no | no |
| GCA_001914545.1 | Nitrospirae bacterium 13_2_20CM_62_7 | p_Nitrospirota; c_Nitrospiria; o_Nitrospirales; f_Nitrospiraceae; g_40CM-3-62-11; s_40CM-3-62-11 sp001914955 | no | no |
| GCA_001918075.1 | Nitrospirae bacterium 13_1_40CM_4_62_6 | p_Nitrospirota; c_Nitrospiria; o_Nitrospirales; f_Nitrospiraceae; g_40CM-3-62-11; s_40CM-3-62-11 sp001914955 | no | no |
| GCA_001918505.1 | Nitrospirae bacterium 13_1_40CM_3_62_11 | p_Nitrospirota; c_Nitrospiria; o_Nitrospirales; f_Nitrospiraceae; g_40CM-3-62-11; s_40CM-3-62-11 sp001914955 | no | no |
| GCA_001920515.1 | Nitrospirae bacterium 13_1_20CM_2_62_14 | p_Nitrospirota; c_Nitrospiria; o_Nitrospirales; f_Nitrospiraceae; g_40CM-3-62-11; s_40CM-3-62-11 sp001914955 | no | no |
| GCA_005877575.1 | Nitrospirae bacterium | p_Nitrospirota; c_Nitrospiria; o_Nitrospirales; f_Nitrospiraceae; g_40CM-3-62-11; s_40CM-3-62-11 sp001914955 | no | no |
| GCA_005877605.1 | Nitrospirae bacterium | p_Nitrospirota; c_Nitrospiria; o_Nitrospirales; f_Nitrospiraceae; g_40CM-3-62-11; s_40CM-3-62-11 sp001914955 | no | no |
| GCA_002420045.1 | Nitrospira sp. UBA5702 | p_Nitrospirota; c_Nitrospiria; o_Nitrospirales; f_Nitrospiraceae; g_Nitrospira; s_Nitrospira sp002420115 | no | no |
| GCA_003529185.1 | Nitrospira sp. | p_Nitrospirota; c_Nitrospiria; o_Nitrospirales; f_Nitrospiraceae; g_Nitrospira; s_Nitrospira sp002420115 | no | no |
| GCA_002083355.1 | Nitrospira sp. HN-bin3 | p_Nitrospirota; c_Nitrospiria; o_Nitrospirales; f_Nitrospiraceae; g_Nitrospira; s_Nitrospira sp900078515 | no | no |
| GCA_002483475.1 | Nitrospira sp. UBA7655 | p_Nitrospirota; c_Nitrospiria; o_Nitrospirales; f_Nitrospiraceae; g_Nitrospira; s_Nitrospira sp900078515 | no | no |
| GCA_900696515.1 | uncultured Nitrosospira sp. | p_Nitrospirota; c_Nitrospiria; o_Nitrospirales; f_Nitrospiraceae; g_Nitrospira; s_Nitrospira sp900078515 | no | no |
| GCA_002083405.1 | Nitrospira sp. SG-bin2 | p_Nitrospirota; c_Nitrospiria; o_Nitrospirales; f_Nitrospiraceae; g_Nitrospira; s_Nitrospira sp900078535 | no | no |
| GCA_003576955.1 | Nitrospira sp. | p_Nitrospirota; c_Nitrospiria; o_Nitrospirales; f_Nitrospiraceae; g_Nitrospira_A; s_Nitrospira_A sp001567445 | no | no |
| GCA_900696565.1 | uncultured Nitrosospira sp. | p_Nitrospirota; c_Nitrospiria; o_Nitrospirales; f_Nitrospiraceae; g_Nitrospira_A; s_Nitrospira_A sp001567445 | no | no |
| GCA_002254325.1 | Nitrospira sp. UW-LDO-02 | p_Nitrospirota; c_Nitrospiria; o_Nitrospirales; f_Nitrospiraceae; g_Nitrospira_A; s_Nitrospira_A sp900170025 | no | no |
| GCA_002299405.1 | Nitrospira sp. UBA667 | p_Nitrospirota; c_Nitrospiria; o_Nitrospirales; f_Nitrospiraceae; g_Nitrospira_A; s_Nitrospira_A sp900170025 | no | no |
| GCA_002380995.1 | Nitrospira sp. UBA4129 | p_Nitrospirota; c_Nitrospiria; o_Nitrospirales; f_Nitrospiraceae; g_Nitrospira_A; s_Nitrospira_A sp900170025 | no | no |
| GCA_002473245.1 | Nitrospira sp. UBA7240 | p_Nitrospirota; c_Nitrospiria; o_Nitrospirales; f_Nitrospiraceae; g_Nitrospira_A; s_Nitrospira_A sp900170025 | no | no |
| GCA_003456185.1 | Nitrospira sp. | p_Nitrospirota; c_Nitrospiria; o_Nitrospirales; f_Nitrospiraceae; g_Nitrospira_A; s_Nitrospira_A sp900170025 | no | no |
| GCA_005116935.1 | Nitrospira sp. | p_Nitrospirota; c_Nitrospiria; o_Nitrospirales; f_Nitrospiraceae; g_Nitrospira_A; s_Nitrospira_A sp900170025 | no | no |
| GCA_002435405.1 | Nitrospira sp. UBA6488 | p_Nitrospirota; c_Nitrospiria; o_Nitrospirales; f_Nitrospiraceae; g_Nitrospira_D; s_Nitrospira_D sp002435325 | no | no |

| ID | NCBI Organism Name | GTDB Taxonomy | GTDB species representative | NCBI type material |
|---|---|---|---|---|
| GCA_005116865.1 | Nitrospira sp. | p_Nitrospirota; c_Nitrospiria; o_Nitrospirales; f_Nitrospiraceae; g_Nitrospira_D; s_Nitrospira_D sp002869855 | no | no |
| GCA_003569385.1 | Nitrospirae bacterium | p_Nitrospirota; c_Nitrospiria; o_Nitrospirales; f_Nitrospiraceae; g_Palsa-1315; s_Palsa-1315 sp002737345 | no | no |
| GCA_005788665.1 | Nitrospiraceae bacterium | p_Nitrospirota; c_Nitrospiria; o_Nitrospirales; f_Nitrospiraceae; g_Palsa-1315; s_Palsa-1315 sp002737345 | no | no |
| GCA_005793845.1 | Nitrospiraceae bacterium | p_Nitrospirota; c_Nitrospiria; o_Nitrospirales; f_Nitrospiraceae; g_Palsa-1315; s_Palsa-1315 sp002737345 | no | no |
| GCA_005798745.1 | Nitrospiraceae bacterium | p_Nitrospirota; c_Nitrospiria; o_Nitrospirales; f_Nitrospiraceae; g_Palsa-1315; s_Palsa-1315 sp002737345 | no | no |
| GCA_005116825.1 | Nitrospira sp. | p_Nitrospirota; c_Nitrospiria; o_Nitrospirales; f_Nitrospiraceae; g_Palsa-1315; s_Palsa-1315 sp002869885 | no | no |
| GCA_003152135.1 | Nitrospira sp. | p_Nitrospirota; c_Nitrospiria; o_Nitrospirales; f_Nitrospiraceae; g_Palsa-1315; s_Palsa-1315 sp003135435 | no | no |
| GCA_001803925.1 | Nitrospirae bacterium RIFCSPHIGHO2_02_FULL_42_12 | p_Nitrospirota; c_Nitrospiria_A; o_HDB-SIOI813; f_HDB-SIOI813; g_HDB-SIOI813; s_HDB-SIOI813 sp001805205 | no | no |
| GCA_001805025.1 | Nitrospirae bacterium GWA2_42_11 | p_Nitrospirota; c_Nitrospiria_A; o_HDB-SIOI813; f_HDB-SIOI813; g_HDB-SIOI813; s_HDB-SIOI813 sp001805205 | no | no |
| GCA_001805235.1 | Nitrospirae bacterium RIFCSPLOWO2_02_42_7 | p_Nitrospirota; c_Nitrospiria_A; o_HDB-SIOI813; f_HDB-SIOI813; g_HDB-SIOI813; s_HDB-SIOI813 sp001805205 | no | no |
| GCA_003477265.1 | Nitrospiraceae bacterium | p_Nitrospirota; c_Nitrospiria_A; o_HDB-SIOI813; f_HDB-SIOI813; g_HDB-SIOI813; s_HDB-SIOI813 sp001805205 | no | no |
| GCA_003508715.1 | Nitrospiraceae bacterium | p_Nitrospirota; c_Nitrospiria_A; o_HDB-SIOI813; f_HDB-SIOI813; g_HDB-SIOI813; s_HDB-SIOI813 sp001805205 | no | no |
| GCA_002897815.1 | bacterium BMS3Abin07 | p_Nitrospirota; c_Thermodesulfovibrionia; o_Thermodesulfovibrionales; f_BMS3Bbin05; g_BMS3Bbin05; s_BMS3Bbin05 sp002897855 | no | no |
| GCA_002897875.1 | bacterium BMS3Bbin06 | p_Nitrospirota; c_Thermodesulfovibrionia; o_Thermodesulfovibrionales; f_JdFR-85; g_BMS3Abin08; s_BMS3Abin08 sp002897935 | no | no |
| GCA_002010755.1 | Nitrospirae bacterium JdFR-87 | p_Nitrospirota; c_Thermodesulfovibrionia; o_Thermodesulfovibrionales; f_JdFR-88; g_JdFR-88; s_JdFR-88 sp002011795 | no | no |
| GCA_002376155.1 | Nitrospiraceae bacterium UBA3562 | p_Nitrospirota; c_Thermodesulfovibrionia; o_Thermodesulfovibrionales; f_JdFR-88; g_JdFR-88; s_JdFR-88 sp002011795 | no | no |
| GCA_002376445.1 | Nitrospiraceae bacterium UBA3568 | p_Nitrospirota; c_Thermodesulfovibrionia; o_Thermodesulfovibrionales; f_JdFR-88; g_JdFR-88; s_JdFR-88 sp002011795 | no | no |
| GCA_002753435.1 | Nitrospirae bacterium | p_Nitrospirota; c_Thermodesulfovibrionia; o_Thermodesulfovibrionales; f_Magnetobacteriaceae; g_HCH-1; s_HCH-1 sp001541255 | no | no |
| GCA_002753455.1 | Nitrospirae bacterium | p_Nitrospirota; c_Thermodesulfovibrionia; o_Thermodesulfovibrionales; f_Magnetobacteriaceae; g_Magnetobacterium; s_Magnetobacterium casensis | no | no |
| GCA_001873265.1 | Nitrospirae bacterium CG2_30_41_42 | p_Nitrospirota; c_Thermodesulfovibrionia; o_Thermodesulfovibrionales; f_SM23-35; g_0-14-3-00-41-53; s_0-14-3-00-41-53 sp002780895 | no | no |
| GCA_002782165.1 | Nitrospirae bacterium CG_4_8_14_3_um_filter_41_47 | p_Nitrospirota; c_Thermodesulfovibrionia; o_Thermodesulfovibrionales; f_SM23-35; g_0-14-3-00-41-53; s_0-14-3-00-41-53 sp002780895 | no | no |
| GCA_002785245.1 | Nitrospirae bacterium CG_4_10_14_0_8_um_filter_41_23 | p_Nitrospirota; c_Thermodesulfovibrionia; o_Thermodesulfovibrionales; f_SM23-35; g_0-14-3-00-41-53; s_0-14-3-00-41-53 sp002780895 | no | no |
| GCA_002787195.1 | Nitrospirae bacterium CG11_big_fil_rev_8_21_14_0_20_41_14 | p_Nitrospirota; c_Thermodesulfovibrionia; o_Thermodesulfovibrionales; f_SM23-35; g_0-14-3-00-41-53; s_0-14-3-00-41-53 sp002780895 | no | no |
| GCA_002790775.1 | Nitrospirae bacterium CG_4_9_14_3_um_filter_41_27 | p_Nitrospirota; c_Thermodesulfovibrionia; o_Thermodesulfovibrionales; f_SM23-35; g_0-14-3-00-41-53; s_0-14-3-00-41-53 sp002780895 | no | no |
| GCA_001312085.1 | Thermodesulfovibrio aggregans JCM 13213 | p_Nitrospirota; c_Thermodesulfovibrionia; o_Thermodesulfovibrionales; f_Thermodesulfovibrionaceae; g_Thermodesulfovibrio; s_aggregans | no | yes |
| GCF_000482825.1 | Thermodesulfovibrio islandicus DSM 12570 | p_Nitrospirota; c_Thermodesulfovibrionia; o_Thermodesulfovibrionales; f_Thermodesulfovibrionaceae; g_Thermodesulfovibrio; s_yellowstonii | no | yes |
| GCF_006538445.1 | Thermodesulfovibrio sp. Kuro-1 | p_Nitrospirota; c_Thermodesulfovibrionia; o_Thermodesulfovibrionales; f_Thermodesulfovibrionaceae; g_Thermodesulfovibrio; s_yellowstonii | no | no |
| GCA_001803715.1 | Nitrospirae bacterium GWD2_44_7 | p_Nitrospirota; c_Thermodesulfovibrionia; o_Thermodesulfovibrionales; f_UBA1546; g_UBA1546; s_UBA1546 sp001805105 | no | no |
| GCA_001803945.1 | Nitrospirae bacterium RIFOXYA2_FULL_44_9 | p_Nitrospirota; c_Thermodesulfovibrionia; o_Thermodesulfovibrionales; f_UBA1546; g_UBA1546; s_UBA1546 sp001805105 | no | no |
| GCA_003500245.1 | Nitrospiraceae bacterium | p_Nitrospirota; c_Thermodesulfovibrionia; o_Thermodesulfovibrionales; f_UBA1546; g_UBA1546; s_UBA1546 sp001805105 | no | no |
| GCA_003510545.1 | Nitrospiraceae bacterium | p_Nitrospirota; c_Thermodesulfovibrionia; o_Thermodesulfovibrionales; f_UBA1546; g_UBA1546; s_UBA1546 sp001805105 | no | no |
| GCA_002323715.1 | Nitrospiraceae bacterium UBA1546 | p_Nitrospirota; c_Thermodesulfovibrionia; o_Thermodesulfovibrionales; f_UBA1546; g_UBA1546; s_UBA1546 sp001871685 | no | no |
| GCA_002771505.1 | Nitrospirae bacterium CG22 | p_Nitrospirota; c_Thermodesulfovibrionia; o_Thermodesulfovibrionales; f_UBA1546; g_UBA1546; s_UBA1546 sp001871685 | no | no |
| GCA_002780905.1 | Nitrospirae bacterium CG02_land_8_20_14_3_00_44_33 | p_Nitrospirota; c_Thermodesulfovibrionia; o_Thermodesulfovibrionales; f_UBA1546; g_UBA1546; s_UBA1546 sp001871685 | no | no |
| GCA_002781345.1 | Nitrospirae bacterium CG01_land_8_20_14_3_00_44_22 | p_Nitrospirota; c_Thermodesulfovibrionia; o_Thermodesulfovibrionales; f_UBA1546; g_UBA1546; s_UBA1546 sp001871685 | no | no |
| GCA_002782185.1 | Nitrospirae bacterium CG_4_8_14_3_um_filter_44_28 | p_Nitrospirota; c_Thermodesulfovibrionia; o_Thermodesulfovibrionales; f_UBA1546; g_UBA1546; s_UBA1546 sp001871685 | no | no |
| GCA_002783085.1 | Nitrospirae bacterium CG_4_10_14_3_um_filter_44_29 | p_Nitrospirota; c_Thermodesulfovibrionia; o_Thermodesulfovibrionales; f_UBA1546; g_UBA1546; s_UBA1546 sp001871685 | no | no |
| GCA_002790755.1 | Nitrospirae bacterium CG_4_9_14_3_um_filter_44_28 | p_Nitrospirota; c_Thermodesulfovibrionia; o_Thermodesulfovibrionales; f_UBA1546; g_UBA1546; s_UBA1546 sp001871685 | no | no |
| GCA_003234985.1 | Nitrospirae bacterium | p_Nitrospirota; c_Thermodesulfovibrionia; o_Thermodesulfovibrionales; f_UBA6898; g_UBA6898; s_UBA6898 sp003252075 | no | no |
| GCA_003156875.1 | Nitrospiraceae bacterium | p_Nitrospirota; c_Thermodesulfovibrionia; o_Thermodesulfovibrionales; f_UBA9935; g_Fen-1308; s_Fen-1308 sp003170655 | no | no |
| GCA_003156895.1 | Nitrospiraceae bacterium | p_Nitrospirota; c_Thermodesulfovibrionia; o_Thermodesulfovibrionales; f_UBA9935; g_Fen-1308; s_Fen-1308 sp003170655 | no | no |
| GCA_003157305.1 | Nitrospiraceae bacterium | p_Nitrospirota; c_Thermodesulfovibrionia; o_Thermodesulfovibrionales; f_UBA9935; g_Fen-1308; s_Fen-1308 sp003170655 | no | no |
| GCA_003158615.1 | Nitrospiraceae bacterium | p_Nitrospirota; c_Thermodesulfovibrionia; o_Thermodesulfovibrionales; f_UBA9935; g_Fen-1308; s_Fen-1308 sp003170655 | no | no |
| GCA_003161685.1 | Nitrospiraceae bacterium | p_Nitrospirota; c_Thermodesulfovibrionia; o_Thermodesulfovibrionales; f_UBA9935; g_Fen-1308; s_Fen-1308 sp003170655 | no | no |
| GCA_003162155.1 | Nitrospiraceae bacterium | p_Nitrospirota; c_Thermodesulfovibrionia; o_Thermodesulfovibrionales; f_UBA9935; g_Fen-1308; s_Fen-1308 sp003170655 | no | no |
| GCA_003162365.1 | Nitrospiraceae bacterium | p_Nitrospirota; c_Thermodesulfovibrionia; o_Thermodesulfovibrionales; f_UBA9935; g_Fen-1308; s_Fen-1308 sp003170655 | no | no |
| GCA_003168405.1 | Nitrospiraceae bacterium | p_Nitrospirota; c_Thermodesulfovibrionia; o_Thermodesulfovibrionales; f_UBA9935; g_Fen-1308; s_Fen-1308 sp003170655 | no | no |
| GCA_003168475.1 | Nitrospiraceae bacterium | p_Nitrospirota; c_Thermodesulfovibrionia; o_Thermodesulfovibrionales; f_UBA9935; g_Fen-1308; s_Fen-1308 sp003170655 | no | no |
| GCA_003170055.1 | Nitrospiraceae bacterium | p_Nitrospirota; c_Thermodesulfovibrionia; o_Thermodesulfovibrionales; f_UBA9935; g_Fen-1308; s_Fen-1308 sp003170655 | no | no |
| GCA_001803605.1 | Nitrospirae bacterium GWA2_46_11 | p_Nitrospirota; c_Thermodesulfovibrionia; o_Thermodesulfovibrionales; f_UBA9935; g_GWB2-47-37; s_GWB2-47-37 sp001803635 | no | no |
| GCA_003453275.1 | Nitrospiraceae bacterium | p_Nitrospirota; c_Thermodesulfovibrionia; o_Thermodesulfovibrionales; f_UBA9935; g_GWB2-47-37; s_GWB2-47-37 sp001803635 | no | no |
| GCA_003538695.1 | Nitrospiraceae bacterium | p_Nitrospirota; c_Thermodesulfovibrionia; o_Thermodesulfovibrionales; f_UBA9935; g_GWB2-47-37; s_GWB2-47-37 sp001803635 | no | no |
| GCA_002897755.1 | bacterium BMS3Abin10 | p_Nitrospirota; c_Thermodesulfovibrionia; o_UBA6902; f_BMS3Bbin08; g_BMS3Bbin08; s_BMS3Bbin08 sp002897775 | no | no |
| GCA_002898055.1 | bacterium BMS3Abin09 | p_Nitrospirota; c_Thermodesulfovibrionia; o_UBA6902; f_UBA6902; g_BMS3Abin09; s_BMS3Abin09 sp002897915 | no | no |
| GCA_002451095.1 | Nitrospiraceae bacterium UBA6907 | p_Nitrospirota; c_Thermodesulfovibrionia; o_UBA6902; f_UBA6902; g_UBA6902; s_UBA6902 sp002451135 | no | no |
| GCA_003508235.1 | Nitrospiraceae bacterium | p_Nitrospirota; c_Thermodesulfovibrionia; o_UBA6902; f_UBA6902; g_UBA6902; s_UBA6902 sp002451135 | no | no |
| GCA_001803725.1 | Nitrospirae bacterium GWD2_57_8 | p_Nitrospirota; c_UBA9217; o_UBA9217; f_UBA9217; g_GWC2-57-13; s_GWC2-57-13 sp001805055 | no | no |
| GCA_003483085.1 | Nitrospiraceae bacterium | p_Nitrospirota; c_UBA9217; o_UBA9217; f_UBA9217; g_GWC2-57-13; s_GWC2-57-13 sp001805055 | no | no |
